# Chromatin boundary permeability is controlled by CTCF conformational ensembles

**DOI:** 10.1101/2025.11.25.690553

**Authors:** Sergei Rudnizky, Peter J. Murray, Emily W. Sørensen, Theo J. R. Koenig, Sushil Pangeni, Raquel Merino-Urteaga, Hemani Chhabra, Laura Caccianini, Iain F. Davidson, Manuel Osorio-Valeriano, Paul W. Hook, Paul Meneses, Jingzhou Hao, Jasmin S. Zarb, Nikos S. Hatzakis, Winston Timp, Lucas Farnung, Seychelle M. Vos, Jan-Michael Peters, Aleksei Aksimentiev, Taekjip Ha

**Author notes:** These authors contributed equally to this work (Sergei Rudnizky, Peter J. Murray, Emily W. Sørensen, Theo J. R. Koenig).

## Abstract

Genomes are organized into chromatin loops through cohesin-mediated extrusion, with CTCF acting as a polar boundary element. As cohesin approaches CTCF at kilobase-per-second speeds, it must rapidly choose whether to stall or bypass. How CTCF encodes this probabilistic decision within a brief encounter window has remained unclear. Here we show that CTCF governs this probabilistic outcome by rapidly sampling a dynamic ensemble of conformations generated by spontaneous rearrangements of its DNA-binding zinc fingers. This ensemble is tuned by DNA sequence, CpG methylation, nearby nucleosomes, and the cohesin regulator PDS5A before cohesin engagement. Upon cohesin binding, PDS5A enhances loop-anchor mechanical stability, reinforcing orientation-dependent boundaries. These findings establish conformational ensemble tuning, rather than static occupancy, as a regulatory principle linking base pair–scale motions to megabase-scale genome organization.

**One sentence summary:** Chromatin boundary function is governed not by CTCF occupancy alone, but by a tunable ensemble of DNA-bound conformations that probabilistically gates cohesin capture.

## Introduction

Eukaryotic genomes are organized into loops, which shape chromatin contacts, defining chromosomal regulatory units (*1*–*4*). In mammals, the eleven zinc finger (ZF) transcription factor CTCF (*5*) plays a central role in regulating these loops by acting as a barrier (*6*–*10*) to cohesin-mediated DNA extrusion (*11, 12*), facilitating the formation of topologically associated domains (TADs) (*1, 3, 9, 10, 13*). Looping factors are broadly conserved across vertebrates (*14*), and their dysregulation has been linked to developmental disorders (*15*), cohesinopathies (*16*), and cancer (*17, 18*).

During loop formation, NIPBL loads cohesin and activates ATPase-dependent cohesin extrusion (*11*). Upon encountering CTCF sites, translocating cohesin complexes either stall or bypass to continue extrusion (*19*). While CTCF functions as an asymmetric roadblock with a higher probability of stalling cohesin when first encountered from its N-terminus (*19, 20*), it remains a semi-permeable barrier even in this orientation (*19*). CTCF boundary permeability is a functionally significant feature that tunes diverse stochastic cellular processes, such as regulation of enhancer-promoter interactions (*17, 21*) and antibody diversification (*22*).

CTCF-cohesin encounters occur in the dynamic chromatin landscape, where TAD boundary strength is modulated by motif sequence variation (*23*), CpG methylation (*17, 18, 24*), nucleosome positioning (*25*), and other cellular processes (*26*–*28*). The cohesin regulator PDS5 (*29, 30*) is essential for maintaining TAD boundaries, with its acute loss weakening insulation (*13, 31, 32*). PDS5-associated cohesin is mutually exclusive with NIPBL-bound cohesin (*29, 30*), suggesting dynamic competition between distinct cohesin states (*33*). How this cohesin stall-or-bypass decision is tuned by local chromatin context and cohesin subunit composition remains unclear (*34*).

Furthermore, cohesin capture by CTCF is constrained to a tight spatiotemporal window, with CTCF having only ∼15-50 ms to interact with a cohesin complex translocating on DNA at 1-3 kb/s (*33, 35*). Within this brief window, which is orders of magnitude shorter than CTCF’s minute-scale residence time on chromatin (*36*), the cohesin complex must interact with binding sites on CTCF’s N-terminal intrinsically disordered region (IDR) (*37*–*39*). The recapitulation of orientation-dependent, partial stalling *in vitro* (*19*) suggests these capture properties are inherent to CTCF, motivating a mechanistic dissection of how a single CTCF-DNA complex is capable of rapid capture, while maintaining a probability of bypass.

Here, we resolve the conformational ensemble of CTCF-DNA complexes at the scale of individual zinc-finger domains and the same millisecond timescales of cohesin encounters. We show that chromatin environment and cohesin-regulating factors tune this ensemble to control cohesin capture and boundary permeability across scales, bridging base-pair-scale dynamics to megabase-scale genome organization.

### An unwrapped CTCF conformation is enriched at cohesin-occupied sites

We reasoned that because CTCF boundary strength can be modulated independently of CTCF occupancy (*31*), it must be encoded in *how* a bound CTCF engages chromatin. By using GpC methylation footprinting, which labels only unprotected DNA, coupled to nanopore sequencing (NanoNOMe) (*40, 41*) (Figs. 1A, S1, Materials and Methods), we sought to directly resolve the binding state of single CTCF molecules and single ZF domains in cells. To do this, we examined CTCF footprints across >20,000 high-confidence motifs (Materials and Methods). Here, we interpret protection from GpC methylation within a ZF triplet as ZF binding. This approach recapitulated the canonical CTCF footprint flanked by well-positioned nucleosomes (*25, 42*–*44*), consistent with the assay faithfully capturing chromatin-bound CTCF-DNA complexes (Fig. S1B). On average, GpCs within central ZFs (ZFs 3-8) were more protected than those at the periphery (ZFs 1-2, 9-11) (Fig. 1B). This observation of more frequently dissociated peripheral ZFs is consistent with previous studies showing that CTCF’s core ZFs have higher affinity for the motif than the peripheral ZFs (*24, 45*–*47*).

**Fig. 1:**
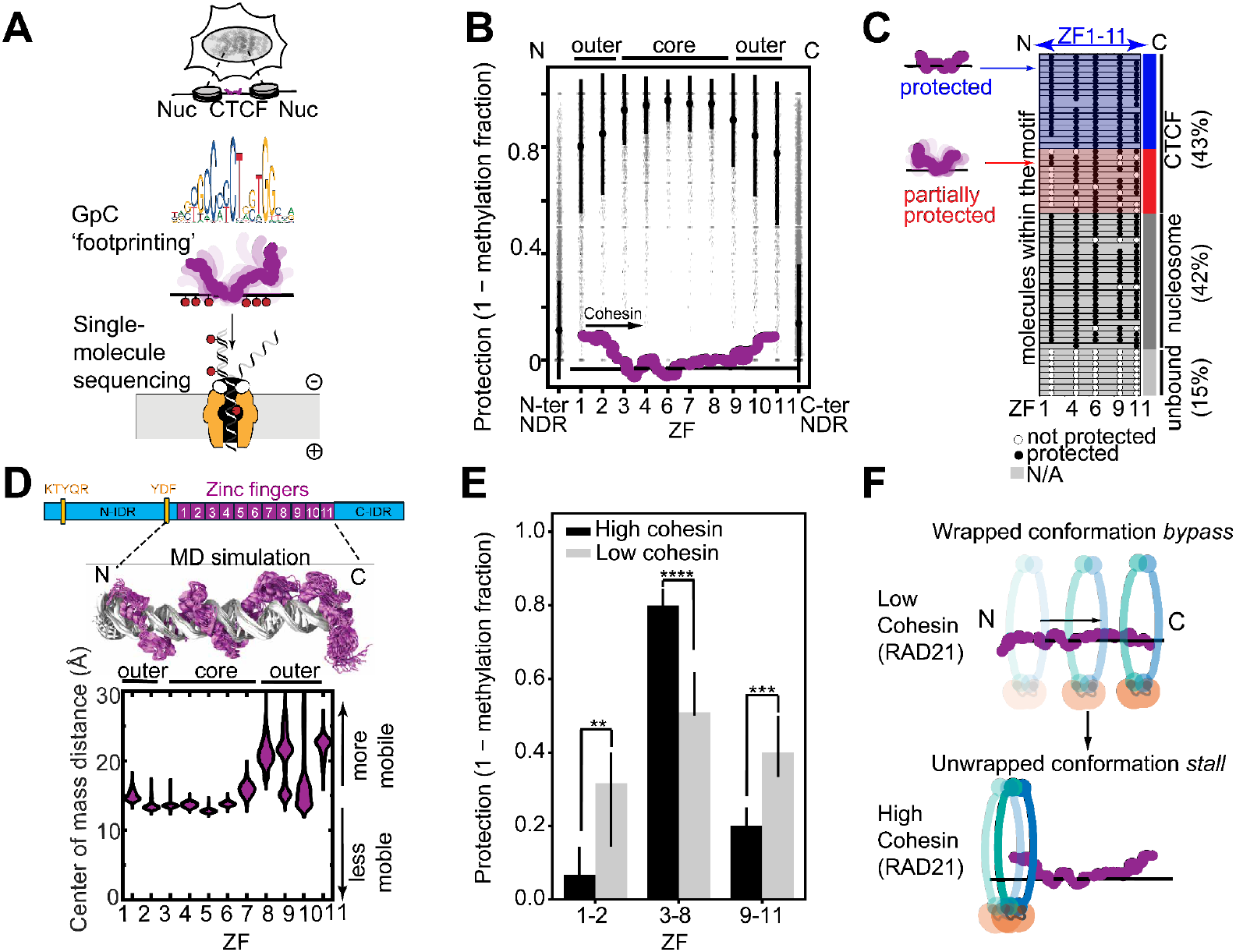
Cohesin preferentially associates with an unwrapped CTCF conformation in cells. **(A)** A schematic representation of the nanopore sequencing GpC footprinting assay in HEK293T cells. **(B)** GpC inaccessibility at CTCF-bound sites grouped by ZF triplet. The mean triplet protections (1 - methylated fraction) for each motif are shown as gray points with mean ± standard deviation across all motifs shown as black dots and bars. Data are given as means ± SD with n referring to the number of included motifs followed by the corresponding number of reads in parentheses: ZF1 = 0.80 ± 0.25, n = 3,643 motifs (72,125 reads); ZF2 = 0.85 ± 0.23, n = 1,020 (16,999); ZF3 = 0.94 ± 0.13, n = 3,914 (77,737); ZF4 = 0.96 ± 0.11, n = 9,147 (184,500); ZF6 = 0.97 ± 0.08, n = 3,132 (65,642); ZF7 = 0.96 ± 0.11, n = 3,916 (85,908); ZF8 = 0.96 ± 0.09, n = 4,396 (93,376); ZF9 = 0.90 ± 0.18, n = 1,905 (36,131); ZF10 = 0.84 ± 0.23, n = 2,811 (53,307); ZF11 = 0.78 ± 0.27, n = 2,837 (53,615); N-ter NDR = 0.11 ± 0.18, n = 22,634 (514,821); C-ter NDR = 0.14 ± 0.18, n = 22,718 (513,016). “N-ter NDR” and “C-ter NDR” refer to regions 20-60 bp away from the CTCF motif in the N-terminal or C-terminal direction, respectively, situated in accessible, nucleosome depleted regions (NDRs) (*44*) (Figs. S1B, S20D). ZF5 is excluded from the analysis due to low GpC occurrence. The “cohesin” arrow emphasizes the CTCF-cohesin encounter orientating that is more efficient for stalling. **(C)** All single-molecule sequencing reads (n = 58 reads) at a representative CTCF binding site (chr16:8792017-8792147). Black circles indicate unmethylated GpCs, white circles indicate methylated GpCs, and grey boxes indicate positions where methylation detection is not possible (e.g., insufficient/uncalled methylation signal in the read). Reads are grouped, from top to bottom, with displayed averages taken across all reads at all motifs (n = 1,061,848 reads), as CTCF-bound (43%), “nucleosomal” (42%) (*48*), or “unbound” (15%) (Materials and Methods). At CTCF-bound reads, protected molecules are estimated to make up about 60% of all reads and partially protected molecules make up the remaining about 40%. See Fig. S1 and Materials and Methods for estimation calculations. **(D)** Above: a to-scale domain map of the CTCF protein depicting its N-terminal IDR, 11 ZF domains, C-terminal IDR, and cohesin-interacting KTYQR and YDF motifs (*39*). Middle: Multiple overlaid snapshots (every 40 ns) of CTCF bound to the endogenous *PDGFRA* insulator sequence sampled by 1 µs molecular dynamics (MD) simulations. A simulation of CTCF bound to its consensus motif and corresponding pLDDT and PAE plots for the AlphaFold 3 (*50*) initial conditions for both motif sequences can be found in Fig. S2. Below: Simulated distribution of distance between the center of mass of each ZF and its 3 nearest DNA bp for each ZF in the endogenous insulator sequence. A second replicate for endogenous insulator and two replicates for the CTCF consensus sequence are shown in Fig. S2. **(E)** GpC inaccessibility within ZF clusters (1-2, 3-8, 9-11) for CTCF sites with the lowest (1st decile “Low RAD21”, grey) and highest (10th decile, “High RAD21”, black) RAD21 ChIP signal (downloaded from the ENCODE portal www.encodeproject.org with the identifier ENCFF241ZVM). Data are shown as median with 95% confidence intervals (CI). Low RAD21 (ZF1-2, ZF3-8, ZF9-11): 0.32 (0.14-0.40 CI) (n = 212), 0.51 (0.50-0.62 CI) (n = 628), 0.40 (0.33-0.50 CI) (n = 422). High RAD21 (ZF1-2, ZF3-8, ZF9-11): 0.07 (0–0.14 CI) (n = 305, p = 0.009), 0.80 (0.73-0.83 CI) (n = 743, p = 1.16x10^−11^), 0.20 (0.13-0.25 CI) (n = 414, p = 0.0005). P-values are calculated with a two-tailed Mann-Whitney test, with significance indicated as **p<0.01, ***p<0.001, ****p<0.0001. **(F)** A cartoon depicting a model where cohesin (red ring) is preferentially associated with an unwrapped CTCF conformation.

Inspecting individual CTCF-bound reads (Materials and Methods) (*48*) allowed us to examine the ZF-binding state of single CTCF molecules in cells. The protein footprints appeared in two forms: fully protected and partially protected (Fig. 1C). Across the motifs, we estimated ∼60% of molecules to be completely protected and ∼40% to be partially protected (Figs. 1C, S1C-D, Materials and Methods). Both populations, as expected for functional CTCF-DNA complexes, were flanked by phased nucleosomes (*49*) (Fig. S1E). Conditional probability analysis of ZF co-binding within single molecules revealed strong correlations between neighboring ZFs, suggesting that ZFs tend to associate and dissociate in stronger core, and weaker peripheral groups (Fig. S1F).

To test whether this preferential dissociation of peripheral relative to core ZFs reflects an intrinsic property of the CTCF–DNA complex, we performed all-atom MD simulations. As model systems, we used AlphaFold 3 (*50*)-predicted structures of CTCF bound to the consensus motif or the endogenous *PDGFRA* insulator, whose disruption has been implicated in the development of gliomas (*17*), as the initial state of the simulations. In agreement with nanopore sequencing results, this simplified system was sufficient to recapitulate weaker association of CTCF’s peripheral ZFs than its core ZFs for both motifs examined (Figs. 1D, S2). As CTCF’s ZFs are typically “snaked” through the major groove (*20, 24, 46, 47*), this behavior supports the existence of two distinct populations: “unwrapped” (peripheral ZFs dissociated) and “wrapped” (all ZFs associated) CTCF molecules.

If the wrapped and unwrapped states are functionally relevant to cohesin stalling, cohesin-enriched CTCF sites in cells should preferentially exhibit one of these conformations. To link these conformational states to cohesin association, we stratified CTCF motifs by chromatin immunoprecipitation sequencing (ChIP-seq) signal for the RAD21 cohesin subunit (*33*) and compared GpC protection patterns across ZF clusters. At the most cohesin-associated CTCF motifs, we observed a distinct ZF binding profile matching the unwrapped conformation that was not present at low cohesin occupancy CTCF sites (Fig. 1E). This strong enrichment of dissociated peripheral ZFs, particularly near the cohesin-interacting CTCF N-terminus (*20, 38, 51*), suggests that the unwrapped conformation may facilitate cohesin association with CTCF (Fig. 1F).

The existence of two conformations (Fig. 1C) and position-dependent fluctuations of CTCF ZFs (Fig. 1D) led us to ask whether the highly mobile ZFs could enable switching between the wrapped conformation and the cohesin-linked, unwrapped conformation.

### CTCF ZFs rapidly scan discrete DNA-bound conformations at the timescale of cohesin extrusion

To experimentally test whether the high ZF mobility observed in MD simulations could lead to larger-scale interconversion between the wrapped and unwrapped conformations, we developed a single-molecule optical tweezers assay to monitor CTCF ZF-DNA contacts and their dynamics at physiological timescales (Fig. 2A). As a model CTCF motif, we continued to focus on the endogenous, oncogenic *PDGFRA* insulator (*17*).

**Fig. 2:**
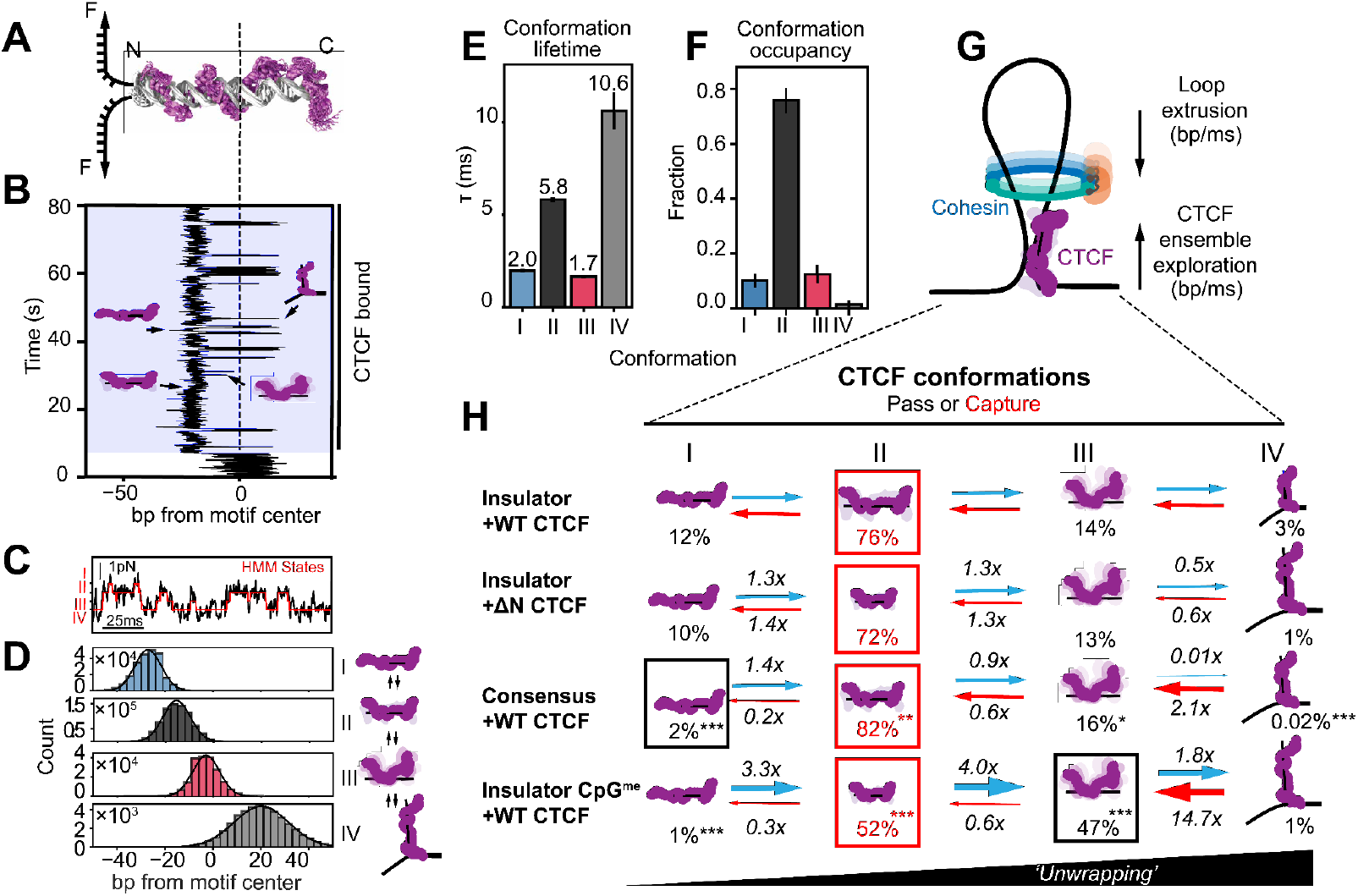
CTCF explores distinct, chromatin-dependent conformations at the timescale of loop extrusion. **(A)** A schematic representation of DNA unzipping while encountering an N-terminally oriented CTCF-bound complex. The depicted CTCF molecule is the same overlaid snapshots of the CTCF MD simulation shown in Fig. 1E. **(B)** A representative trace of DNA fluctuations at the endogenous *PDGFRA* insulator motif (“insulator”). This trace is plotted as the location of the unzipping fork in base pairs from the motif center, with two distinct regimes, unbound (unshaded) and CTCF-bound (shaded). Cartoons are interpretations of CTCF conformation based on the ZF triplets that are bound and unbound at the times indicated by the black arrows. **(C)** A zoom-in of the trace shown in **(B)**, prior to conversion from pN to bp (Materials and Methods), illustrating the mapped HMM states. **(D)** Left: each state’s mapped base pair location constructed from 4 representative replicates of CTCF dynamics bound to the endogenous insulator motif as four Gaussians (conformations I, II, III, and IV, from top to bottom). Gaussian fits (black lines) provide mean ± standard deviation for each state with n representing number of transitions into the state (see Table S1): -27.1 ± 5.5 bp (n = 234,310), - 15.5 ± 5.7 bp (n = 772,942), -3.0 ± 5.7 bp (n = 200,878), and 20.1 ± 11.4 bp (n = 41,865) for conformations I-IV, respectively. Y-axes are scaled individually to adjust for variable state occupancy for visualization purposes. Right: a model depicting interconversion of hypothesized corresponding conformations illustrated as cartoons. **(E-F)** Lifetime **(E)** and conformational occupancy (fraction of time spent) **(F)** in each state measured from the N-terminus of the endogenous insulator motif. Error bars represent the standard deviation between replicates (Table S2). **(G)** A model depicting the importance of CTCF conformational exploration timescales matching that of loop extrusion and its implication in stochastic cohesin capture. **(H)** Summary of CTCF conformational ensemble dynamics probed from the N-terminal orientation for, from top to bottom: insulator with WT CTCF, insulator with ΔN CTCF, consensus with WT CTCF, and CpG-methylated insulator with WT CTCF. Interpretations of conformations are depicted as cartoons with their occupancy percentages written below. Red boxes indicate the conformation with the highest occupancy for each condition, with the black box emphasizing the high abundance of conformation III in the CpG-methylated condition and the low abundance of state I in the consensus motif condition. Transition rates are written over and under arrows as multiples relative to WT CTCF bound to the insulator. All transition rates and occupancies can be found in Figs. S10-13, S15-17, Table S2. P-values are calculated with a two-tailed Mann-Whitney test relative to WT CTCF bound to the insulator and can be found in Table S2, with significance indicated as *p<0.05, **p<0.01, ***p<0.001.

By creating an unzipping fork at the CTCF binding site and holding it at a fixed position, we observed the DNA fork to stochastically open and close (*52, 53*) (Figs. 2B, S3). Upon CTCF binding, these fluctuations were partially suppressed, such that the position of the unzipping fork directly reported how many base pairs were forced into a double-stranded state by the bound protein. As the CTCF-bound motif sampled different degrees of DNA openness (Figs. 2B, S3C), changes in fork position served as a real-time readout of CTCF-DNA structure. Persistence of these fluctuations in the subsequent absence of free CTCF indicated probing of a single bound CTCF molecule, as opposed to dissociation and rebinding (Fig. S3D). Complete unzipping irreversibly disrupted the complex and produced discrete force rips at the expected CTCF motif position, consistent with probing of a single CTCF-DNA complex (Materials and Methods, Figs. S3, S4). Truncation of either the N- or C-terminal IDR (ΔN or ΔC CTCF) did not alter force rip position or magnitude, suggesting that the signal arose from probing ZF-DNA contacts (Fig. S4I-L).

CTCF-bound DNA was best described by four states, as determined by Hidden Markov Modeling (HMM) fitting (Figs. 2C, S5-7). Mapping the observed states to the motif (Figs. 2D, S3-7) revealed, as with the GpC conditional probability analysis (Fig. S1F), distinct conformations driven by ZFs moving in blocks: early (N-terminal), middle (core), and late (C-terminal). These conformations exchanged almost exclusively in a sequential manner (conformation I ↔ conformation II ↔ conformation III ↔ conformation IV) (Fig. S8). In conformation I, all ZFs were bound; in conformation II, only the early ZFs were dissociated; in conformation III, only the late ZFs remained bound (Figs. 2D, S8). Conformation IV mapped to all ZFs being dissociated without complete CTCF unbinding, likely reflecting a loosely-associated state, as proposed for other transcription factors (*52, 54*) (Figs. 2D, S3-8). These dynamics were force-independent in the examined regime (Fig. S9), N-terminal IDR-independent, and remained stable over time, suggesting we measured an intrinsic property of the CTCF-DNA complex and its ZF domains at equilibrium (Figs. 2H, S14, Materials and Methods) (*53*). The sequential and cooperative nature of these ZF association and dissociation events supports a model where ZFs undergo spontaneous, reversible unwrapping and rewrapping around the DNA motif.

Notably, the conformation lifetimes were on the millisecond scale - the same order of magnitude as cohesin’s estimated ∼15-50 ms passage time through the CTCF footprint (Figs. 2E, 2G) (*11, 12, 19, 20*). Given that CTCF stalls cohesin *in vitro* with about 50-90% efficiency, depending on DNA tension (*19*), we hypothesized that these conformations may exist in kinetic competition as cohesin-compatible and cohesin-incompatible forms of CTCF. Observations that the unwrapped conformation II is most prevalent (76% occupancy) (Fig. 2F) and matches the ZF footprint of the cohesin-enriched sites in cells (Fig. 1E) supports a model where conformation II is the productive, “cohesin-capturing” conformation (Fig. 2G). This model predicts that boundary strength can be tuned by shifting the balance between these conformations, so we next asked whether regulatory chromatin cues can bias this ensemble.

### Chromatin context modulates the conformational ensemble of DNA-bound CTCF

As local chromatin features have been shown to strengthen or weaken CTCF-bound boundaries (*18, 23, 55*), we next explored whether they could act through the tuning of the CTCF conformational ensemble. To address this question, we tested the effects of motif sequence, CpG methylation, and nearby nucleosome presence.

Perhaps counterintuitively, the high affinity consensus motif (*24*) was not associated with more ZFs being bound relative to the *PDGFRA* motif. Rather, the unwrapped conformation II increased in both state occupancy (from 76% to 82%) and lifetime (from 5.8 ms to 9.6 ms) (Figs. 2H, S10-13, S15-18, Table S2). Because consensus-like sequences are enriched at constitutive TAD boundaries (*23, 55*), this observation supports the model that the unwrapped conformation is cohesin-compatible.

Because CpG methylation is broadly associated with reduced CTCF binding, but does not always weaken TAD boundaries (*18*), we asked whether methylation can also act through reshaping the conformational ensemble of DNA-bound CTCF. Methylating the regulatory CpG within the ZF4 triplet of this oncogenic insulator motif (*17*) did not abolish CTCF binding (Fig. S19), but caused conformation II occupancy to fall from 76% to 52% and conformation III occupancy to increase from 13% to 47% (Figs. 2H, S10-13, S15-17, Table S2). This conformation redistribution matches the expectation that methylation destabilizes cytosine contacts in the central ZFs (Fig.S19A) promoting their unwrapping and pushing the system towards the highly-unwrapped conformation III (*17, 23, 24*)

Finally, as boundary-associated CTCF sites reside in nucleosome-free regions flanked by well-positioned nucleosomes (*25, 42*–*44*) (Figs. S1B, Figs. S20A-D), we examined CTCF’s conformational ensemble in the presence of a physiologically-positioned, regulatory N-terminal nucleosome (*25*) (Fig. S20C-D). A reconstituted, positioned nucleosome (*56*) (Fig. S20E-M) shifted the landscape towards a more partially-unwrapped profile (increasing C-terminal conformation II from 46% to 69% state occupancy) (Fig. S20L-M).

Together, these results demonstrate that chromatin context reshapes the CTCF conformational ensemble, allowing cells to integrate cues to potentially tune cohesin barrier function independently of protein occupancy changes.

### PDS5A binds CTCF to steer the ensemble toward cohesin capture

Because cohesin complexes can contain one of the mutually exclusive ATPase-activating NIPBL or ATPase-inactivating PDS5 subunits, we next asked whether these factors, which may be enriched at cohesin-capturing CTCF sites in cells, bias this chromatin-tuned conformational ensemble.

Correlating our ZF-resolved binding profiles in cells (Fig. 1A-C) with publicly available ChIP-seq datasets (*57*) revealed that, similar to RAD21 (Figs. 1E, S21), PDS5A enrichment was negatively correlated with peripheral ZF binding and positively correlated with central ZF binding (*24, 46*) (Fig. 3A-C), indicative of preferential association with the partially unwrapped CTCF conformation. In contrast, NIPBL (Figs. 3A-B, S21I) did not exhibit this patterning and its depletion minimally affected RAD21-ZF correlations (Figs. 3B, S21C-D). Consistently, publicly-available mass spectrometry data (*58, 59*) showed that PDS5A was associated with chromatin-bound CTCF to a much higher degree than NIPBL was (Fig. S21A). Together, these observations indicate that, although NIPBL is essential for activating cohesin-mediated loop extrusion (*11*) needed for encounters with CTCF, the cohesin-capturing CTCF conformation is predominately associated with PDS5A in cells.

**Fig. 3:**
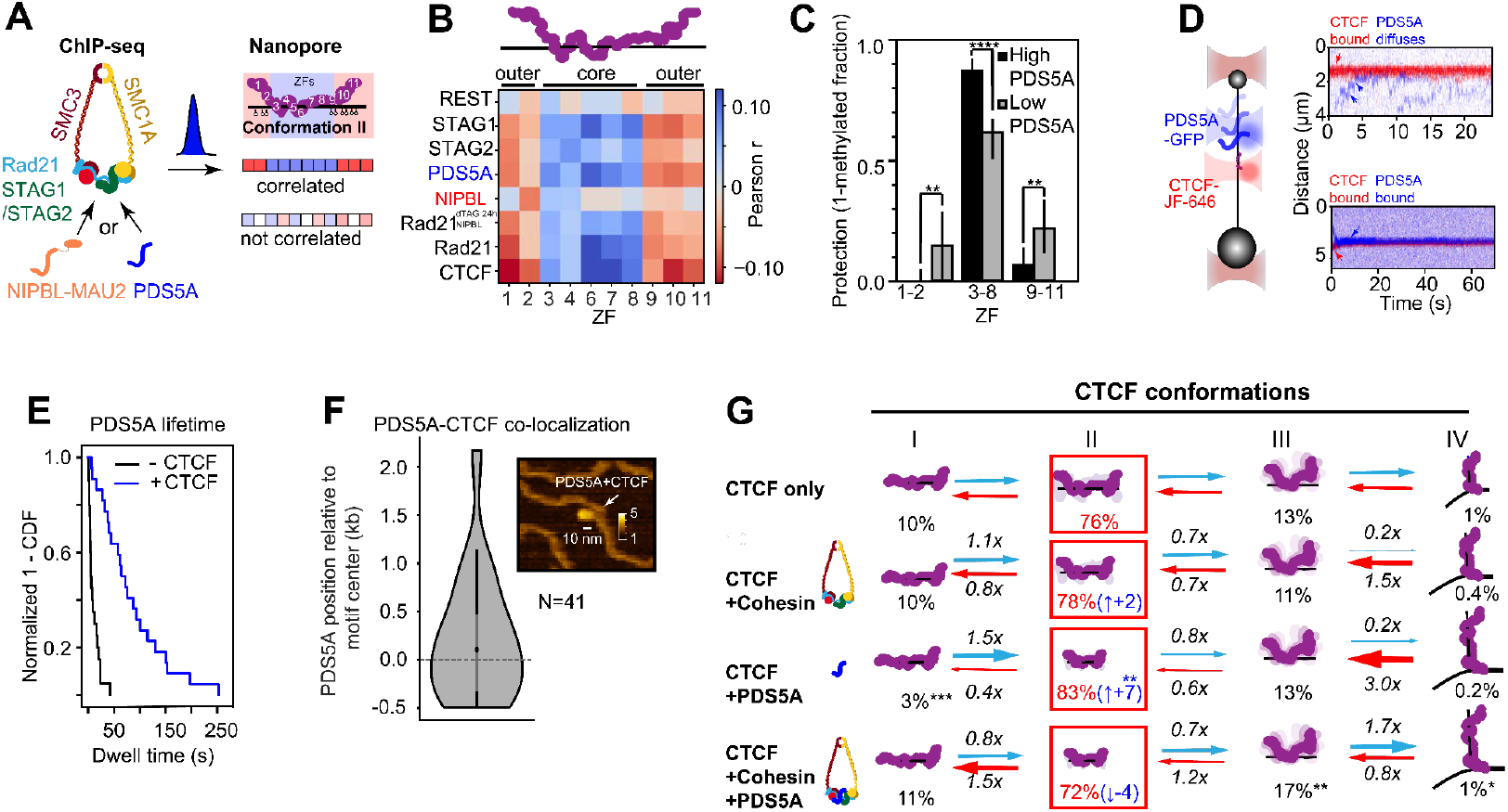
PDS5A engages CTCF to promote the cohesin-capturing conformation. **(A)** Cartoon of cohesin subunits and a schematic representation showing expected results for ChIP-seq and nanopore footprinting correlation analysis if the data are or are not correlated. **(B)** Pearson correlation between publicly available ChIP-seq datasets of cohesin subunits in retinal pigment epithelial (RPE1) cells from Popay et al (*57*) and REST (HEK293), as a negative control (downloaded from the ENCODE portal www.encodeproject.org with the identifier ENCFF981ILL), and GpC inaccessibility for a subset of partially-protected molecules (as shown in Fig. 1A-C) at each ZF. See Fig. S21 for additional proteins, cell types, and significance levels. **(C)** GpC inaccessibility (1 - methylated fraction) within ZF clusters (1-2, 3-8, 9-11) for CTCF sites with the least (1st decile “Low PDS5A”, gray) and most (10th decile, “High PDS5A”, black) PDS5A ChIP signal. Data are shown as median protection with 95% confidence intervals (CI). Low PDS5A (ZF1-2, ZF3-8, ZF9-11): 0.14 (0-0.29 CI) (n = 207), 0.61 (0.50-0.67 CI) (n = 616), 0.21 (0.10-0.33 CI) (n = 363). High PDS5A (ZF1-2, ZF3-8, ZF9-11): 0 (0–0.05 CI) (n = 298, p = 0.0013), 0.87 (0.80-0.92 CI) (n = 687, p < 0.0001), 0.06 (0-0.14 CI) (n = 399, p = 0.0068). P-values are calculated with a two-tailed Mann-Whitney test, with significance indicated as **p<0.01, ****p<0.0001. **(D)** Left: schematic representation of an optical tweezers assay where CTCF-JF646 and PDS5A-GFP binding and sliding are imaged with confocal scanning on long, taut dsDNA. Right: representative kymographs of PDS5A encountering stably bound CTCF binding. The top kymograph shows transient PDS5A interaction (or “bouncing”), while the bottom kymograph shows continuous colocalization for more than 60 seconds (or “docking”). **(E)** 1-CDF of dwell times of PDS5A without CTCF (black; median = 5.4 s, n = 20) and colocalized with CTCF (red; median = 67.8 s, n = 22). **(F)** Binding position of PDS5A, as determined by AFM, on a symmetric DNA substrate containing 2 CTCF consensus motifs (median position = 106 bp from CTCF motif, -160 bp to 297 bp 95% CI, n = 41). Position is given as distance from the nearest CTCF binding site. The inset displays a representative AFM image of PDS5A colocalized with DNA at the expected CTCF binding motif (Materials and Methods). **(G)** Transition rates between four conformational states of CTCF for, from top to bottom, CTCF alone, CTCF and cohesin, CTCF and PDS5A, and CTCF, cohesin, and PDS5A. Interpretations of conformations are depicted as cartoons with their occupancy percentages written below. Red boxes indicate the most occupied state. Transition rates are written over and under arrows as multiples relative to WT CTCF bound to the insulator. All transition rates and occupancies can be found in Figs. S10-13, S15-17, and Table S2. P-values are calculated with a two-tailed Mann-Whitney test relative to N-terminal WT CTCF bound to *PDGFRA* and can be found in Table S2, with significance indicated as *p<0.05, **p<0.01.

This strong enrichment of PDS5A, but not NIPBL, with chromatin-bound CTCF could be explained by two scenarios: PDS5A could arrive after NIPBL-cohesin pauses at CTCF sites, or PDS5A could prebind CTCF before an extruding NIPBL-cohesin complex arrives. To distinguish between these possibilities, we developed a two-color optical-tweezers assay (*60*) to mimic PDS5A-CTCF encounters by tracking fluorescently-labeled CTCF and PDS5A-GFP on a DNA substrate harboring CTCF-binding sites (Figs. 3D, S22A). After appearing to locate its binding site via 1D diffusion (D = 0.62 ± 0.45 kb^2^/s, consistent with a previous report (*19*)) (Fig. S22D), CTCF positionally stabilized (Figs. S22B-C, E-F). PDS5A also exhibited 1D sliding behavior on DNA until encountering CTCF (Figs. 3D, S22G-I). Strikingly, PDS5A did not bypass either diffusive or immobilized CTCF, either “bouncing” off or “docking” onto it (Figs. 3D, S22H-I). Docking onto immobilized CTCF dramatically stabilized PDS5A on DNA (from 5.4 s to 67.8 s median lifetime) (Fig. 3E). Although we were blind to motif orientation, the presence of two discrete behaviors is consistent with the asymmetric barrier behavior of CTCF (*19, 20*). Consistently, complementary atomic force microscopy (AFM) imaging revealed enrichment of PDS5A density at CTCF motif positions (Figs. 3F, S23).

Having demonstrated that PDS5A associates with CTCF independently of cohesin (Fig. 3D-F) and that the unwrapped CTCF conformation is enriched at sites with high PDS5A prevalence (Fig. 3B-C), we next examined how PDS5A modulated CTCF’s conformational ensemble. PDS5A presence shifted the ensemble towards the cohesin-capturing, unwrapped conformation (from 76% to 83% state occupancy) (Figs. 3G, S24) to levels comparable to the consensus sequence (82% state occupancy) (Figs. 2H, 3G, S15, Table S2). This selective enrichment supports a “priming” mechanism in which PDS5A preconditions CTCF for productive cohesin capture but remains to be demonstrated during active extrusion. This effect was not observed in the presence of the cohesin complex (SMC1a, SMC3, RAD21, and STAG1) alone or when both cohesin and PDS5A were present (Fig. 3G), consistent with PDS5A acting upstream of cohesin engagement.

### PDS5A-cohesin stabilizes loop anchors to enforce orientation-dependent loop boundaries

Having observed enrichment of the cohesin-capturing CTCF conformation by PDS5A (Fig. 3G), we next asked, once bound, whether PDS5A directly impacts the permeability of polar CTCF barriers through ensemble modulation. If PDS5A were to tune the IDR-independent conformational exploration of CTCF’s ZFs (Fig. 2H), we hypothesize it must directly engage them. To test this hypothesis, we modeled the structure of the PDS5A-CTCF insulator complex using AlphaFold 3 (*50*) (Figs. 4A, 4C, S25). In addition to recapitulating the known interaction at CTCF’s N-terminal IDR (*38*), the model indeed predicted additional, albeit lower confidence, contacts between PDS5A and CTCF’s N-terminal ZFs (Figs. 4A, 4C, S25).

**Fig. 4:**
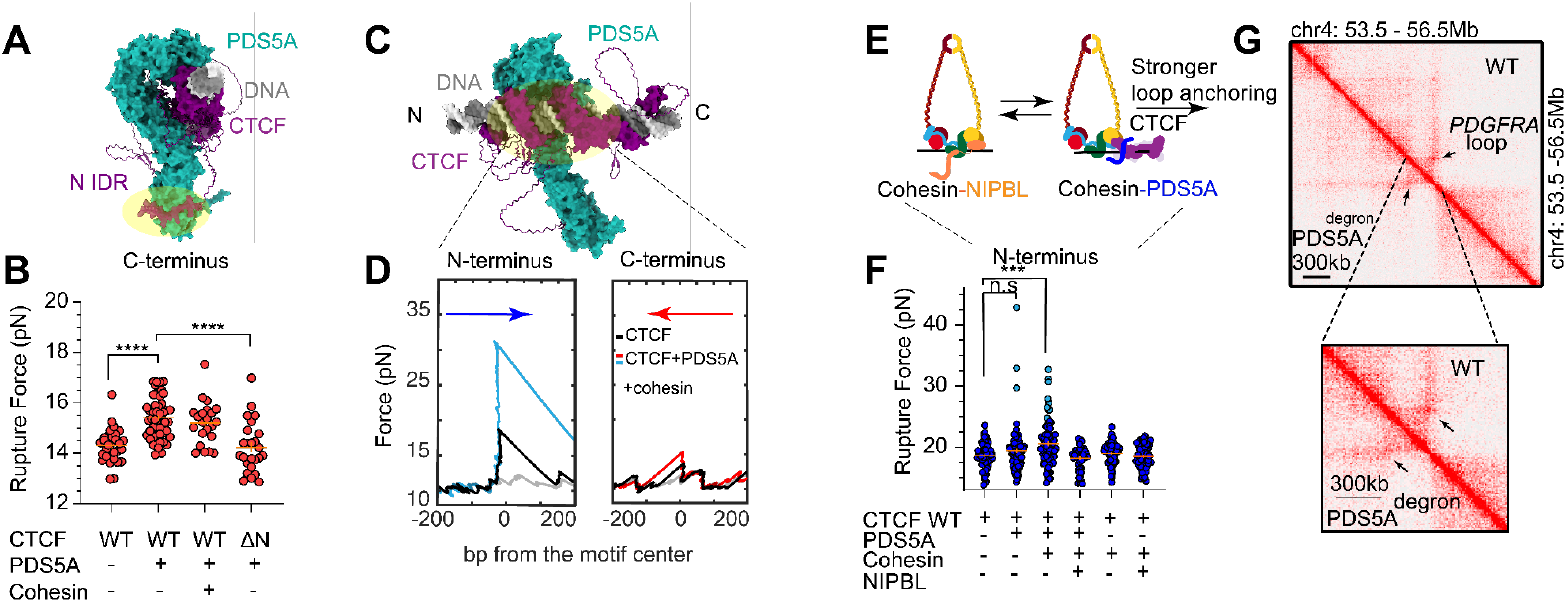
PDS5A-cohesin mechanically stabilizes loop anchors to enforce orientation-dependent insulation. **(A)** AlphaFold 3-predicted structure (*50*) of CTCF (purple) bound to *PDGFRA* insulator dsDNA (grey) and PDS5A (turquoise). A yellow-shaded oval highlights the previously reported interaction between PDS5A and the KTYQR motif in CTCF’s N-terminal IDR (*38*). Corresponding pLDDT and PAE plots can be found in Fig. S25. **(B)** Rupture forces of C-terminal unzipping of WT and ΔN CTCF in the presence and absence of PDS5A and cohesin. Mean rupture forces are 14.3 ± 0.1 pN (n = 31) for WT CTCF, 15.5 ± 0.1 pN (n = 52) for WT CTCF and PDS5A, 15.2 ± 0.2 pN (n = 22) for WT CTCF, PDS5A, and cohesin, 14.2 ± 0.2 pN (n = 25) for ΔN CTCF and PDS5A. Data are mean ± standard error. Using a 1-D, two-state K-S test, significance is indicated as follows: ***p<0.001, ****p<0.0001. **(C)** A rotated view of the AlphaFold 3-predicted structure shown in **(A)**, with the predicted interaction between PDS5A and CTCF’s ZFs highlighted with a yellow oval. **(D)** Representative unzipping traces of CTCF in the presence of PDS5A and cohesin from the N-terminal (left, cyan) and C-terminal (right, red) orientation. For comparison, representative unbound DNA (gray) and CTCF-only (black) unzipping traces are shown. **(E)** A cartoon model of the kinetic competition between the mutually exclusive PDS5A and NIPBL subunits of cohesin. Cohesin-PDS5A, and not cohesin-NIPBL, is depicted bound to CTCF. **(F)** Rupture forces of N-terminal unzipping of WT CTCF in the presence and absence of PDS5A, cohesin, and NIPBL. Mean rupture forces are 18.6 ± 0.3 pN (n = 57) for WT CTCF, 19.4 ± 0.4 pN (n = 96) for WT CTCF and PDS5A, 20.5 ± 0.4 pN (n = 83) for WT CTCF, PDS5A, and cohesin, 18.2 ± 0.3 pN (n = 46) for WT CTCF, PDS5A, NIPBL, and cohesin, 19.0 ± 0.2 pN (n = 76) for WT CTCF and cohesin, 18.5 ± 0.2 pN (n = 66) for WT CTCF, NIPBL, and cohesin. Data are mean ± standard error. Points shown in cyan were predicted to be within a higher rupture force population (Materials and Methods). For WT CTCF and PDS5A, populations had mean rupture forces of 18.9 ± 0.2 pN (n = 93) and 35.2 ± 3.2 pN (n = 3). For WT CTCF, PDS5A, and cohesin populations had mean rupture forces of 19.6 ± 0.3 pN (n = 73) and 27.8 ± 0.9 pN (n = 10). **(G)** A Hi-C map of the *PDGFRA* locus in PLC/PRF/5 cells with and without induced PDS5A FKBP12F36V-dTAG degron. A zoomed-in view emphasizes the loss of the corner dot at the *PDGFRA* insulator. Data were obtained from Yu et al (*32*).

To probe the effect of PDS5A on ZF-DNA contacts directly, we used optical tweezers DNA unzipping to map positions and strength of local protein-DNA interactions (*52, 61*–*65*) (Figs. S4, S26). PDS5A modestly stabilized C-terminal ZF contacts, increasing their mean rupture force (from 14.3 ± 0.1 pN to 15.5 ± 0.1 pN), which was abolished upon N-terminal IDR truncation (mean = 14.2 ± 0.2 pN), consistent with the IDRs established role in PDS5A recruitment to CTCF (Figs. 4B, S26) (*38*). Strikingly, sampling CTCF’s N-terminal ZFs in the presence of PDS5A revealed a distinct population of CTCF-DNA complexes exhibiting an exceptionally high rupture force (mean = 35.2 ± 3.2 pN) (Figs. 4F cyan points, S26B), far above the 15–25 pN range reported for other transcription factors (*52, 54, 64, 66*).

Furthermore, the combination of the cohesin complex and PDS5A increased the abundance of the CTCF molecules with high N-terminal ZF rupture force from 3.1% to 12.0% (Figs. 4D-F, S26). Consistent with this stabilization being PDS5A mediated, the cohesin complex alone was insufficient to observe this N-terminal ZF strengthening (Figs. 4D-F, S26). This PDS5A-dependent stabilization may serve as an anchoring mechanism at loop boundaries, reinforcing their strength, directionality, and possibly insulating capabilities.

In contrast, addition of extrusion-competent NIPBL (Figs. 4E-F, S27) suppressed PDS5A-mediated mechanical stabilization of CTCF N-terminal ZFs, consistent with reported direct competition between the two factors (Figs. 4E, S26) (*31*). As PDS5A and other cohesin subunits, but not NIPBL, are enriched with chromatin-bound, unwrapped CTCF (Figs. 3B-C, S21), this further supports a model in which NIPBL drives extrusion to the boundary, while PDS5A-cohesin stabilizes the post-capture anchor. Indeed, comparison of available Hi-C data before and after PDS5A depletion at the constitutive *PDGFRA* insulator (*17*) showed a reduction in CTCF-mediated insulation (Fig. 4G).

Taken together these data support a unifying, two-step model, in which PDS5A first biases CTCF towards a capture-competent conformation to promote stalling of incoming NIPBL-cohesin and then helps convert the captured complex into a stable, extrusion-inactive loop anchor at the boundary (Fig. 5).

**Fig. 5:**
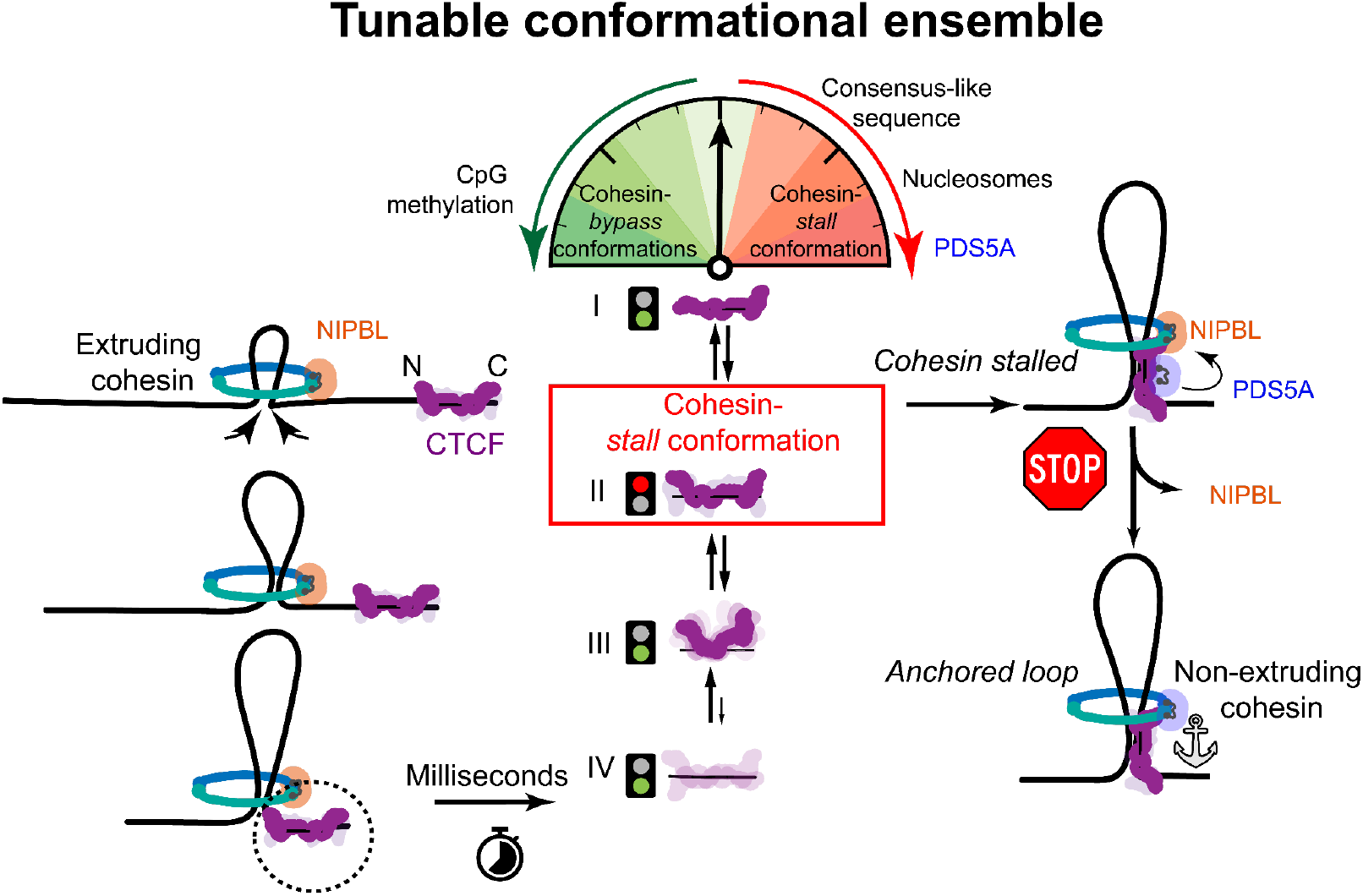
CTCF conformational scanning gates cohesin capture versus read-through to tune boundary permeability. A model depicting a two-step process (*34*) of loop formation and loop stabilization. First, CTCF must stochastically capture rapidly translocating NIPBL-cohesin. To do this, CTCF samples an ensemble of productive and unproductive conformations on the millisecond timescale through sequential unwrapping and rewrapping of blocks of ZFs. This ensemble is tuned by CTCF motif sequence, CpG methylation, and nucleosome positioning. The cohesin subunit PDS5A also shifts this conformational ensemble by pre-binding and priming CTCF in the cohesin-capturing conformation prior to NIPBL-cohesin’s arrival. Once NIPBL-cohesin reaches CTCF, the pre-bound PDS5A exchanges with NIPBL and stabilizes the CTCF-cohesin complex, forming highly stabilized loop boundaries.

## Discussion

As CTCF has only ∼15-50 ms to engage rapidly extruding cohesin at a boundary element, capture-versus-bypass decisions are unlikely to be dictated exclusively by minute-scale CTCF binding and unbinding (*34, 36*). Instead, we propose that boundary permeability is encoded in a fast, DNA-anchored CTCF conformational ensemble that tunes the probability of cohesin capture versus bypass (Fig. 5). Our data indicate that this ensemble is shaped by the local chromatin environment, which modulates cohesin stalling probability, likely by altering the availability or geometry of cohesin-engaging motifs on the CTCF-DNA complex. In this framework, imperfect insulation is integral to boundary function: semi-permeability permits regulated bypass, which is essential for stochastic cellular processes, like antibody diversification, while preserving efficient capture. It also helps explain why essential, strong boundaries, such as those that separate chromosome superdomains, are often comprised of tandem arrays of consecutive, predominantly unidirectionally oriented CTCF motifs. In such arrays, multiple, closely spaced motifs increase the effective probability of cohesin stalling and buffer boundary strength against local chromatin variability. Conceptually, this shifts boundary regulation from an occupancy-centric view to a state-centric one, in which permeability is encoded by ensemble composition rather than binding alone.

PDS5A biases the CTCF ensemble composition toward a capture-competent conformation prior to cohesin engagement, priming an orientation-dependent encounter. We propose that at the boundary, capture at a CTCF site may then serve as a platform for cohesin subunit exchange, promoting the transition from NIPBL-bound to a mechanically robust, PDS5-bound cohesin loop anchor. Defining the timing and molecular determinants of this on-site transition *in vivo* will be crucial to connect cohesin regulatory state to boundary lifetime and permeability. A key prediction of this model is that boundary strength can be tuned via shifts in ensemble occupancy or PDS5A recruitment to alter stalling probability and anchor stability without necessarily changing CTCF occupancy, consistent with a recent preprint (*31*).

Boundary function requires a fine balance between stability and flexibility: loops must be persistent enough to support insulation yet dynamic enough to adapt to changing chromatin context. We propose that rapid, local conformational sampling of DNA-anchored CTCF, tuned by sequence, chromatin state, and cofactors provides a kinetic principle (*67*) for how boundaries are established, maintained, and remodeled without wholesale disassembly. In this view, sequence and chromatin features modulate the ensemble, while cohesin regulators, such as PDS5, convert the ensemble properties into boundary permeability and lifetime. Together, our results demonstrate how base pair-scale conformational dynamics of a DNA-bound factor can be directly translated into genome-scale regulation of chromatin architecture. More broadly, we suggest that ensemble tuning may represent a general strategy by which other sequence-specific factors regulate partner engagement, cooperativity, or responsiveness during transient genomic encounters.

## Acknowledgments

The authors thank Mathias Madalinski for HPLC purification of the JF646 HaloTag ligand; Andrew Drabeck for STAG1 purification and msfGFP-PDS5A cloning; Maciej Zakzec for cloning of N- and C-terminal CTCF deletion constructs; and Anders Hansen and Gina M. Dailey (from the laboratories of Robert Tjian and Xavier Darzacq) for providing a plasmid for mouse CTCF expression. We are grateful to the laboratories of Sua Myong, Bradley Bernstein, Anders Hansen, Frederick Alt, Ariel Kaplan and Carl Wu for critical discussions. We thank Cees Dekker, Hylkje Geertsema, and Jos Zwanikken for their comments during T.J.R.K.’s thesis defense, and Brian Analikwu for advice during his thesis. We also thank Xiaona Tang and colleagues in the Carl Wu laboratory for server access, and Roman Barth for technical advice on in vitro loop-extrusion assays. We acknowledge the laboratory of Jesse Dixon for generating cohesin-subunit ChIP-seq data. We thank David Mohr and the Genetic Resources Core Facility (GRCF) at Johns Hopkins University for performing NextSeq and NovaSeq runs. We thank Yana Li of the Johns Hopkins Eukaryotic Tissue Culture Facility for expressing mouse CTCF. Finally, we thank the Hatzakis, Farnung, Timp, Aksimentiev, Peters, Vos, and Ha laboratories for stimulating discussions throughout the project. J-MP is also an adjunct professor at the Medical University of Vienna and an external member of the Max Planck Institute of Biochemistry, Martinsried. During the preparation of this work the authors used OpenAI’s ChatGPT for assisting with programming and refining writing. After using this tool, the authors reviewed and edited the content as needed and take full responsibility for the content of the published article. This article is subject to HHMI’s Immediate Access to Research policy, which requires that this article be made publicly available as initial and revised preprints deposited on a designated preprint server under a CC BY 4.0 license.

## Funding

European Molecular Biology Organization (EMBO) fellowship (ALTF 825-2021)

Human Frontier Science Program (HFSP) fellowship (LT0049/2022-L)

Beckman Fellowship, UIUC

TACC Frontera (MCB20012)

ACCESS allocation grant (MCA05S028)

National Science Foundation Science and Technology Center for Quantitative Cell Biology (NSF STC-QCB, Grant No. 2243257)

Novo Nordisk foundation challenge center for Optimised Oligo escape (NNF23OC0081287)

Novo Nordisk foundation Center for 4D cellular dynamics (NNF22OC0075851)

Villum foundation experiment (40801)

National Institutes of Health (R35 GM122569, U01 DK127432)

Howard Hughes Medical Institute

Boehringer Ingelheim

The European Union (ERC AdG 101020558 LoopMechRegFun)

The European Union (MSCA 101072505 CohesiNet)

The Vienna Science and Technology Fund (LS19-029)

## Author contributions

S.R., P.J.M., E.W.S. and T.J.R.K. contributed equally to this work. S.R., P.J.M. and T.H. conceived the project. S.R., P.J.M. and T.J.R.K. generated molecular constructs and performed optical tweezers measurements, and S.R. and P.J.M. analyzed rupture-force data. E.W.S. performed analysis of dynamic measurements and quantified AFM data. H.C. and A.A. performed and analyzed MD simulations. I.F.D. expressed, purified and labelled human CTCF and mutants, and J.S.Z. purified mouse CTCF. L.C. expressed and purified cohesin and PDS5A, and M.O.-V. expressed and purified NIPBL. P.M. and J.H. reconstituted nucleosomal constructs. R.M.-U., P.M., P.J.M. and S.R. performed single-molecule TIRF experiments. S.P., P.J.M. and S.R. performed confocal optical tweezers and AFM measurements. P.J.M. performed AlphaFold3 structure predictions. S.R. performed Cas9 accessibility assays in cells. S.R. and P.W.H. performed nanopore sequencing, and S.R. analyzed the nanopore data with input from P.J.M., E.W.S. and P.W.H. S.R. carried out and analyzed CUT&RUN and MNase-seq, and mined ChIP-seq, ChIP-MS, ChIP-SICAP and Hi-C datasets. N.S.H., W.T., L.F., S.M.V., J.-M.P. and A.A. provided discussion, expertise and support for collaborations. S.R., P.J.M. and E.W.S. wrote the manuscript in coordination with T.H. and with input from J.-M.P. and I.F.D. T.H. supervised the project.

## Data, code, and materials availability

All data are available in the main text or the supplementary materials. Custom scripts, figures, and data can be found at Zenodo. Custom scripts are also available on GitHub (https://github.com/pmurra20/CTCF-boundary-permeability-code). Raw data is archived and available upon request. Instructions for synthesizing materials are provided in the supplementary materials and are also available upon reasonable request.

## Supplementary Materials

Materials and Methods

Supplementary Text

Figs. S1 to S28

Tables S1 to S2

References (*1*–*110*)

Data S1

