## Supplementary materials for "Chromatin boundary permeability is controlled by CTCF conformational ensembles"

**The PDF file includes:**

Materials and Methods
Supplementary Text
Figs. S1 to S28
Tables S1 to S2

**Other Supplementary Materials for this manuscript include the following:**

Data S1

### Materials and Methods

Protein preparation:

Human CTCF cloning

The insect cell expression construct for full length human CTCF (pLib 10xHis-CTCF_1-727-Halo-Flag; #Mz92) was previously described (*19*). N-terminally deleted human CTCF (10xHis-CTCF_266-727-Halo-Flag; #Mz93) and C-terminally deleted human CTCF (10xHis-Halo-CTCF_1-579-Flag; #Mz51) were generated by PCR using Phusion Hot Start Flex DNA Polymerase (NEB M0535S) and inserted into pLib (*68*) using Gibson assembly (*69*).

Human CTCF protein expression and purification

Full length and truncated forms of CTCF were expressed and purified as described (*19*) with modifications. Briefly, *Spodoptera frugiperda* Sf9 cells derived from pupal ovarian tissue of the female fall armyworm (Thermo Fisher Scientific B82501) were cultured at 100 rpm and 27 °C in Grace medium (prepared in-house) supplemented with 10% fetal calf serum (Gibco A5256801), 2 mM L-glutamine (Gibco 25030-024), 1x Penicillin-Streptomycin (Sigma-Aldrich P0781) and 0.1% Poloxamer 188 (Gibco 24040-032). Bacmids were generated in *E. coli* DH10EmBacY as described (*70*).

For baculovirus infection of Sf9 cells, $1x10^6$ Sf9 cells were seeded on a 6 well plate in 3 ml Sf-900 SFM media (ThermoFisher Scientific 12658027), transfected with bacmid DNA using Fugene 6 (Promega) and incubated for 96 hours at 27 °C. The baculovirus-containing cell supernatant (V0) was then collected and used to infect 50 ml Sf9 cells at a density of 1 $x10^6$ cells/ml in Grace medium supplemented as described above. Cells were grown at 100 rpm and 27 °C, centrifuged 72 hours after infection and 9 ml cell supernatant (V1) was then used to infect 750 ml Sf9 cells at a density of 1.2 $x10^6$ cells/ml in Grace medium supplemented as above in the presence of 0.1 mM $ZnCl_2$. Around 4 x 750 ml cultures were infected per construct. Cells were centrifuged around 54 hours after infection, washed in PBS, frozen in liquid nitrogen and stored at -80 °C.

Around 20 ml cell pellet per construct was lysed by Dounce homogenisation and resuspended to 100 ml in CTCF buffer (35 mM HEPES pH 7.5, 350 mM NaCl, 5% glycerol) supplemented with 0.1 mM $ZnCl_2$, 0.05% Tween-20, 5 mM imidazole, 1 mM PMSF, EDTA-free cOmplete tablet (1 per 50 ml) (Roche 11873580001), 3 mM betamercaptoethanol, 10 µg/ml aprotinin (Carl Roth A162.3), 2 mM benzamidine, benzonase (prepared in-house) and DNase I (800 units, prepared in-house). The lysates were cleared by centrifugation at 41,000 g for 45 min at 4°C. The soluble fractions were incubated with 2 ml NiNTA agarose (Qiagen 30230) for 90 min at 4°C and washed with CTCF buffer supplemented with 0.1 mM $ZnCl_2$, 0.01 % Tween-20, 35 mM imidazole, 1 mM PMSF and 2 mM benzamidine. Proteins were eluted with CTCF buffer supplemented with 0.1 mM $ZnCl_2$, 1 mM PMSF, 2 mM benzamidine and 300 mM imidazole. The eluates were then combined with 2 ml of FLAG-M2 agarose resin (Sigma A2220) and incubated for 2 hours at 4°C. Beads were washed with Flag buffer 350 (35 mM HEPES pH 7.5, 350 mM NaCl, 5% glycerol) supplemented with 0.1 mM $ZnCl_2$. Beads were resuspended in one bed volume of Flag buffer 350 supplemented with 0.1 mM $ZnCl_2$ and incubated protected from light in the presence of excess JF646-HaloTag Ligand at room temperature with rotation for 20 min. Beads were washed extensively with Flag buffer 350 supplemented with 0.1 mM $ZnCl_2$ and then with Flag buffer 350 in the absence of 0.1 mM $ZnCl_2$. Bound proteins were eluted with Flag buffer 350 supplemented with 0.5 mg/ml 3xFlag peptide. Eluates were concentrated using Vivaspin 20 50 kDa MWCO ultrafiltration units (Sartorius VS2032), frozen in liquid nitrogen and stored at -80°C.

Janelia Fluor 646-HaloTag Ligand preparation for CTCF labeling

A 20 mM stock of Janelia Fluor 646 SE (Tocris 6148) and a 50 mM stock of HaloTag Amine (O2) Ligand (Promega P6711) were prepared in dimethylformamide (DMF) immediately prior to use. Aliquots of these stocks were combined, diluted in DMF and supplemented with diisopropylethylamine (final 250 µl reaction mixture: 6.74 mM Janelia Fluor 646 SE, 2.69 mM HaloTag Amine (O2) Ligand, 37 mM diisopropylethylamine). The reaction mixture was incubated at room temperature for ~ 16 hours protected from light. The reaction mixture was diluted 10-fold in solvent A (5 % acetonitrile, 0.1 % formic acid) and purified in five successive reverse phase HPLC runs using an Ultimate 3000 (ThermoFisher Scientific) equipped with a Kinetex 5µ XB-C18 100A, 250 x 4.6mm column using the following gradient of solvents: A (5 % acetonitrile, 0.1 % formic acid), B (acetonitrile, 0.1 % formic acid); 0 – 100 % B in 30 min at a flow rate of 0.8 ml / min. Peak fractions were pooled, lyophilized, resuspended in DMF, frozen in liquid nitrogen and stored at -80°C.

Human STAG1, SMC1a, SMC3, RAD21 and PDS5A cloning

Human STAG1 and PDS5A were amplified from human cDNA. Genes were inserted into 438B (Addgene plasmid 55219) by Ligation Independent Cloning (LIC). The genes were inserted behind a 6xHis tag followed by a Tobacco etch virus protease (TEV) cleavage site. msfGFP-PDS5A was generated by inserting a 6xHis-10xArg-msfGFP-TEV tag before the PDS5A coding region.

Human SMC1a and RAD21 were amplified from Addgene plasmids 32363 and 57666, respectively. SMC3 was amplified from human cDNA. SMC1a and SMC3 were inserted into vector 438A. RAD21 was cloned into vector 438F where RAD21 was followed by a TEV cleavage site and a maltose binding protein (MBP) tag. The subunits were combined into pBig1a to create a co-expression vector (*68*).

Purified plasmid DNA (0.3–0.8 µg) was electroporated into DH10αEMBacY cells to generate bacmids (Geneva Biotech) (*71*). DNA was extracted by isopropanol precipitation from positive clones and transfected into Sf9 cells grown in ESF921 (Expression Systems) with X-tremeGENE9 transfection reagent (Sigma). The resulting virus (V0) was harvested 48–72 h after transfection. Virus was then amplified by infecting 25 mL of Sf9 or Sf21 cells with 25–150 µL of the harvested V0, and grown at 27°C, 300 rpm. Viruses were harvested 48–72 h after proliferation arrest by centrifugation and stored at 4°C.

For protein expression, 600 ml of Hi5 cells at 1 million cells/mL confluency were infected with 300-500 µL of virus and grown at 27° for 60-72h in ESF921 medium. The cells were collected by centrifugation. STAG1 was resuspended in Lysis Buffer 1 (STAG1: 300 mM KCl, 20 mM Tris-HCl pH 7.5, 30 mM Imidazole pH 8.0, 5 mM beta mercaptoethanol (BME), 10% glycerol (v/v), 0.284 µg $ml^{-1}$ leupeptin, 1.37 µg $ml^{-1}$pepstatin A, 0.17 mg $ml^{-1}$ PMSF, and 0.33 mg $ml^{-1}$ benzamidine). PDS5A, GFP-PDS5A were resuspended in Lysis buffer 2 (500 mM NaCl, 25 mM Tris-HCl pH 7.5, 30 mM Imidazole pH 8.0, 5 mM BME, 10% glycerol (v/v), 0.284 µg $ml^{-1}$ leupeptin, 1.37 µg $ml^{-1}$pepstatin A, 0.17 mg $ml^{-1}$ PMSF, and 0.33 mg $ml^{-1}$ benzamidine) at 4°, snap frozen in liquid nitrogen, and stored at -80°. SMC1a-RAD21-MBP-SMC3 pellets were resuspended in Lysis Buffer 3 (Lysis Buffer: 500 mM NaCl, 20 mM Tris-HCl pH 7.9, 30 mM Imidazole pH 8.0, 5 mM BME, 10% glycerol (v/v), 0.284 µg $ml^{-1}$ leupeptin, 1.37 µg $ml^{-1}$pepstatin A, 0.17 mg $ml^{-1}$ PMSF, and 0.33 mg $ml^{-1}$ benzamidine) and immediately processed for purification.

Human STAG1 and PDS5A/PDS5A-GFP purification

All steps were performed at 4° unless otherwise specified. Frozen cell pellets were thawed in a room temperature water bath and lysed by sonication. Lysates were clarified by centrifugation. The clarified lysate was filtered successively through 5 µm and 0.45 µm syringe filters. Filtered lysate was applied to a 5 mL HisTrap (Cytiva) column equilibrated in Lysis Buffer. The loaded lysate was washed with 10 CV of Lysis Buffer, followed by a 5 CV of High Salt Buffer (STAG1: 1000 mM KCl, 20 mM Tris-HCl pH 7.5, 30 mM Imidazole pH 8.0, 5 mM BME, 10% glycerol (v/v). PDS5A and GFP-PDS5A: 1000 mM NaCl, 25 mM Tris-HCl pH 7.5, 30 mM Imidazole pH 8.0, 5 mM BME, 10% glycerol (v/v)), and 5 CV of Lysis Buffer, followed by 5 CV of Low Salt Buffer (STAG1: 150 mM KCl, 20 mM Tris-HCl pH 7.5, 30 mM Imidazole pH 8.0, 5 mM BME, 10% glycerol (v/v), PDS5A and msfPDS5A 150 mM NaCl, 25 mM Tris-HCl pH 7.5, 30 mM Imidazole pH 8.0, 5 mM BME, 10% glycerol (v/v)). The HisTrap was attached to a HiTrapQ column equilibrated in Low Salt buffer. Protein was eluted from the HisTrap column onto HiTrap Q column over a gradient from 0-100% nickel elution buffer (STAG1: 150 mM KCl, 20 mM Tris-HCl pH 7.5, 500 mM Imidazole pH 8.0, 5 mM BME, 10% glycerol (v/v), PDS5A and msfGFP-PDS5A: 150 mM NaCl, 25 mM Tris-HCl pH 7.5, 300 mM Imidazole pH 8.0, 5 mM BME, 10% glycerol (v/v)). After eluting the protein, the HisTrap column removed. The HiTrapQ column was washed with 5 CV of Low Salt buffer. The HiTrapQ column was developed over a gradient from 0%-100% High Salt Buffer. Peak fractions were analyzed on 10% Tris-Glycine SDS-PAGE. Fractions containing the protein of interest were pooled, mixed with 1.5 mg of His6-TEV protease (except for msfGFP-PDS5A), and dialyzed overnight in 7kDa MWCO SnakeSkin tubing (Fisher) at 4° against 1-2 L of Dialysis Buffer (STAG1: 300 mM KCl, 20 mM Tris-HCl pH 7.5, 30 mM Imidazole pH 8.0, 5 mM BME, 10% glycerol (v/v). PDS5A/GFP-PDS5A: 500 mM NaCl, 25 mM Tris-HCl pH 7.5, 30 mM Imidazole pH 8.0, 5 mM BME, 10% glycerol (v/v)).

Protein was removed and applied to a 5 mL HisTrap column (except for msfGFP-PDS5A, which was directly used for size exclusion chromatography), equilibrated in the appropriate Lysis Buffer. This enabled the capture of the 6xHis-MBP tag, 6xHis-TEV protease, and uncleaved protein. The cleaved protein was collected as flow through and concentrated in a 30K MWCO 15 mL Amicon Millipore ultrafiltration device. The concentrated protein was then applied to a STAG1: Superose 6 increase column or an S200 16/600 HiLoad column (Cytiva) or PDS5A/GFP-PDS5A: Superose 6 10/300 pg increase column (Cytiva) equilibrated in Size Exclusion Chromatography Buffer (STAG1: 300 mM KCl, 20 mM Tris-HCl pH 7.5, 0.5 mM Tris (2-carboxyethyl)phosphine (TCEP), 10% glycerol (v/v), PDS5A/GFP-PDS5A: 500 mM NaCl, 25 mM Tris-HCl pH 7.5, 0.5 mM TCEP, 10% glycerol (v/v)). Peak fractions were analyzed on 10% Tris-Glycine SDS-PAGE. Appropriate fractions were pooled and concentrated in a 30K MWCO Amico Millipore ultrafiltration device. Protein was aliquoted, snap frozen in liquid nitrogen, and stored in -80°.

Human Cohesin (SMC1a, SMC3, RAD21) purification

Hi5 cells infected with cohesin virus were harvested after 60-65h of infection and were immediately processed for protein purification. Cells were collected by centrifugation at 238xg for 30 minutes. Cell pellets were resuspended in Lysis Buffer and lysed by sonication. Lysed cells were clarified by centrifugation. The clarified lysate was filtered successively through 5 μm and 0.45 μm syringe filters. Once filtered, the lysate was applied to a custom amylose column packed with 10-15 mL of Amylose resin (NEB), equilibrated in Lysis Buffer. The loaded lysate was washed with 2 CV of Lysis Buffer, followed by a 2 CV High Salt wash (1000 mM NaCl, 20 mM Tris-HCl pH 7.9, 30 mM Imidazole pH 8.0, 5 mM BME, 10% glycerol (v/v)), and 2 CV of Lysis Buffer. Protein was eluted with 2 CV of Amylose Elution Buffer (500 mM NaCl, 20 mM Tris-HCl pH 7.9, 300 mM Maltose, 5 mM BME, 10% glycerol (v/v)). Peak fractions were analyzed by 6% Tris-Glycine SDS-PAGE and Coomassie Staining. Appropriate fractions were pooled and concentrated in a 15 mL 50K MWCO Amicon Millipore ultrafiltration device. Concentrated protein (1-1.5 mL) was loaded on a Superose 6 10/300 increase column (Cytiva), equilibrated in Size Exclusion Chromatography Buffer (500 mM NaCl, 20 mM Tris-HCl pH 7.9, 0.5 mM TCEP, 10% glycerol (v/v)). Peak fractions were analyzed on 10% Tris-Glycine SDS-PAGE and coomassie staining. Appropriate fractions were pooled and concentrated in a 50K MWCO Amicon Millipore ultrafiltration device. Protein was aliquoted, snap frozen in liquid nitrogen, and stored in -80°.

Mouse CTCF protein expression and purification

The mouse fusion protein 3X-FLAG-Halo-TEV-mCTCF-HisX6 recombinant Bacmid DNA (*72*) was generated using the Bac-to-Bac Baculovirus expression system (ThermoFisher 10359016). Sf9 cells were infected with the baculovirus to express the recombinant fusion protein. The recombinant protein was purified from Sf9 cells, as previously described (*72*). Briefly, the cell culture was centrifuged to isolate the cell pellet. The cell pellet was gently resuspended and washed with PBS solution and centrifuged at 3,900 x g for 10 minutes. The cell pellet was gently resuspended in five packed cell volumes of a mouse CTCF high salt buffer (1 M NaCl, 50 mM HEPES pH 7.9, 0.05% NP-40, 10% glycerol, 10 mM 2-mercaptoethanol) and sonicated. The solution was centrifuged for 30 min at 12,000 rpm. The clarified lysate was supplemented with 10mM imidazole pH 8.0. The sample was transferred and incubated with an equilibrated NI-NTA resin (Qiagen 30410) on a rocker for 90 minutes. The beads were washed several times with four column volumes of mouse CTCF high salt buffer supplemented with 20mM imidazole, followed by washing the beads twice with four column volumes of 50 mM HEPES pH 7.9, 10% glycerol, 0.01% NP-40, and 20 mM imidazole. The protein was eluted with 0.5 M NaCl, 50 mM HEPES pH 7.9, 10% glycerol, 0.01% NP-40, and 0.25 M imidazole. The eluted fractions were analyzed by SDS-PAGE gel followed by SimplyBlue SafeStain (Invitrogen LC6060). Selected elution fractions containing protein were combined and transferred to an equilibrated Anti-FLAG M2 Affinity gel (Millipore Sigma A2220) and incubated on a rocker for 3 hours. The resin was washed with 1 M NaCl, 50 mM HEPES pH 7.9, 0.05% NP-40, and 10% glycerol and then equilibrated and washed with 0.2 M NaCl, 50 mM HEPES pH 7.9, 10% glycerol, and 0.01% NP-40. The protein was eluted with 0.4 mg/mL 3XFLAG peptide (Millipore Sigma F4799) in 0.2 M NaCl, 50 mM HEPES pH 7.9, 10% glycerol, and 0.01% NP-40 buffer. Protein concentrations were determined using an SDS PAGE gel stained with ProtoStain Blue Colloidal Coomassie Stain (National Diagnostics EC-727) and were compared to Bovine Serum Albumin (BSA) standards. All protein purification steps were performed at 4°C and in the presence of protease inhibitors (or cOmplete EDTA-free Protease Inhibitor Cocktail (Roche 11836170001)).

Cloning, expression and purification of the NIPBL and MAU2 complex

*H. sapiens* NIPBL and MAU2 were amplified from cDNA and cloned into the MacroLab 438 vector series for expression in insect cells using ligation-independent cloning (*73*).The NIPBL expression construct contained an N-terminal 3x FLAG tag, followed by a Halo tag, and a HRV 3C protease cleavage site, and a C-terminal TEV protease cleavage site, followed by a 10x His tag. MAU2 was cloned into the 438-A vector with no tag. Both subunits were then combined into the pBigBac1b vector (*68*).Plasmids were sequence verified by long-read sequencing. Bacmid, virus, and protein production were then performed as previously described (*74*). The NIPBL/MAU2 complex was expressed in Hi5 insect cells and purified at 4°C. Cells expressing NIPBL/MAU2 were resuspended in lysis buffer (250 mM NaCl, 20 mM Na HEPES pH 7.4, 10% (v/v) glycerol, 30 mM imidazole, 1 mM TCEP, 0.284 μg $ml^{-1}$ leupeptin, 1.37 μg $ml^{-1}$ pepstatin A, 0.17 mg $ml^{-1}$ PMSF and 0.33 mg $ml^{-1}$ benzamidine) and lysed by sonication. The lysate was centrifuged and cleared by ultra-centrifugation. The supernatant containing NIPBL/MAU2 was filtered using 0.22 μm syringe filters. The filtered supernatant was applied to a HisTrap HP 5 mL column (Cytiva 17524801) previously equilibrated in lysis buffer. The column was subsequently washed with 10 CV of lysis buffer and 5 CV of low salt buffer (150 mM NaCl, 20 mM Na·HEPES pH 7.4 at 25 °C, 10% (v/v) glycerol, 1 mM TCEP). The NIPBL/MAU2 complex was eluted onto a 5 mL HiTrap Q (Cytiva 17115401) with 10 CV low salt buffer supplemented with 500 mM Imidazole. The HisTrap column was removed, and the HiTrap Q column was washed with 10 CV of low salt buffer. The NIPBL/MAU2 complex was eluted with a 20 CV linear gradient with high salt buffer (1 M NaCl, 20 mM Na·HEPES pH 7.4 at 25 °C, 10% (v/v) glycerol, 1 mM TCEP). The elution was fractionated and analyzed using SDS-PAGE. Fractions containing NIPBL/MAU2 were pooled and concentrated using an Amicon 100,000 MWCO centrifugal filter unit (Millipore UFC910008). The concentrated sample was applied to a Superose 6 Increase 10/300 GL (Cytiva 29091596) column equilibrated in gel filtration buffer (250 mM NaCl, 20 mM Na·HEPES pH 7.4,10% (v/v) glycerol, 1 mM TCEP pH 8). NIPBL/MAU2-containing fractions were concentrated with an Amicon 100,000 MWCO centrifugal filter unit, aliquots were frozen in liquid nitrogen and stored at −80 °C. Throughout the manuscript, the NIPBL/MAU2 complex will be referred to as “NIPBL”.

Unzipping construct preparation for optical tweezers experiments

The “Y-shaped” DNA constructs for single-molecule unzipping experiments (DNA handles) were generated as described previously (*52*, *64*, *65*) with minor modifications. Two 2000 bp dsDNA handles were PCR-amplified from lambda DNA (NEB N3011) using Q5 DNA High-Fidelity Polymerase (NEB M0491S) to include restriction sites for 3×digoxigenin and 3×biotin tag ligation (to improve stability under force) (*60*) as well as sites for nicking enzymes. The handles were treated with Nt.BbvCI (NEB R0632L) and Nb.BbvCI (NEB R0631L) restriction enzymes to produce complementary ~29-nt overhangs, which were annealed in equimolar ratios to form a ~4000 bp construct in annealing buffer (1x TAE (40 mM Tris-acetate, 10 mM EDTA), 12.5 mM MgAc). A ~230 bp alignment sequence was PCR-amplified from 601c2-plasmid (*60*) with Q5 polymerase, restriction digested with DraIII-HF (NEB R3510S), and ligated to the construct with T4 DNA ligase (NEB M0202M), which was then size selected with gel-excision from E-gels (Thermo Scientific G402021, G402022, G401004) using a Qiaquick Gel Extraction Kit (Qiagen 28704). The digoxigenin- and biotin-modified oligos were ligated to overhangs generated by BglI (NEB R0143S) and underwent DraIII-HF restriction digestion, respectively. Variable sequences (500-1000 bp), including the CTCF consensus sequence, the *PDGFRA* insulator, and the *PDGFRA* insulator fused with a Widom-601 nucleosome positioning sequence, were restriction digested with BbsI-HF (NEB R3539L) to ligate a short DNA hairpin with T4 DNA ligase, and DraIII-HF to create a complementary overhang for purified DNA handles. All PCR, restriction digestion, and ligation reactions were subsequently purified with a Qiaquick PCR Purification Kit (Qiagen 28106) or CleanNGS beads (Bulldog Bio CNGS050). Variable regions contained a single CTCF binding site and were otherwise computationally designed to minimize off-target binding. The CTCF binding sites used were the *PDGFRA* motif (+/- Widom 601 sequence), the consensus motif, or an alternative consensus motif with slight variation in the periphery used for mouse CTCF experiments.

For optical tweezers experiments, the final construct was assembled by mixing Rapid Ligation Buffer (Promega C6711), prepared DNA handles (1 nM final), variable region (20-30 nM final), 1 µL ~2.08 µm, anti-digoxigenin-modified, polystyrene beads (Spherotech DIGP-20-2), 1 µL concentrated T4 DNA ligase (NEB M0202L), and nuclease free water to a final volume of 10 µL. This mixture was incubated at RT for 10 min and diluted to a working concentration by adding 500 µL of experimental buffer.

<u>CpG Methylation of *PDGFRA* constructs</u>

For methylation experiments, 400 ng of the fully-prepared *PDGFRA* unzipping variable region was treated with 1 µL of M.Sssi (NEB M0226S) and 13 uM S-adenosyl-methionine (SAM) (NEB B9003S) following the manufacturer's protocol. The treatment was validated by restriction digestion with HinP1I (NEB R0124S), which is ineffective at digesting methylated DNA. By running an E-gel of the reaction product against an unmethylated positive control, we determined that the vast majority of the sample was protected from digestion and therefore methylated.

<u>Lambda DNA for confocal C-trap and TIRF measurements</u>

Lambda DNA (NEB N3011S) containing 12-nucleotide 5′ overhangs were filled using Klenow fragment lacking exonuclease activity (NEB M0212S). The polymerization reaction included 16 nM lambda DNA, 33 µM each of dATP (Qiagen 201912), dGTP (Qiagen 201912), biotinylated-dUTP (ThermoFisher R0081), and biotinylated-dCTP (ThermoFisher 19518018), and 3 µL of Klenow in a 50 µL final volume with NEB buffer 2. The mixture was incubated at 37 °C for 1 hour and heat-inactivated at 75 °C for 20 minutes. To remove excess nucleotides and proteins, drop dialysis was performed using 0.025 µm VSWP filters (Millipore Sigma VSWP02500) suspended over 20–25 mL of dialysis buffer (10 mM Tris-HCl, pH 8.0; 0.1 mM EDTA) in a petri dish. A 50–150 µL drop of DNA was gently placed on the membrane using a large-orifice pipette tip, ensuring the membrane did not dip into the buffer. Dialysis proceeded at room temperature for at least 1.5 hours, after which the DNA was carefully retrieved and stored at 4 °C for long-term use.

<u>HS-AFM sample preparation</u>

A nearly symmetric, ~8 kb (7,979 bp; including 3 ssDNA bases from RE digestion on both ends) DNA substrate containing consensus CTCF motifs near each end (centered 514 or 511 bp from the end) was ordered within a pBluescript II KS (+) plasmid vector from GenScript Biotech. The plasmid was amplified through transformation into JM109 competent cells (Promega L2005), plating on LB agar plates with ampicillin (Teknova L1004), inoculation in LB media (ThermoFisher 12795027) containing 100 ug/mL ampicillin (Sigma-Aldrich A9518-100G), and purification with the E.Z.N.A.® Plasmid Mini Kit II (Omega Bio-Tek D6945-02). The substrate was then removed from the backbone with DraIII restriction enzyme digestion, as described above, and subsequently size selected with gel-excision purification, as described above. A ~1 kb DNA (1,006 bp) segment of the 8 kb DNA substrate with a central CTCF consensus motif (510 bp and 495 bp from motif center on each side) was prepared using PCR, as described above, with the 8 kb DNA as a template.

Nucleosome reconstitution

The variable region containing a Widom-601 (*56*) positioning sequence was prepared following the same protocol as other variable regions. Once amplified, restriction digested, and hairpin-ligated, ten titrations of different Octamer:DNA ratios were prepared (0.50x, 0.75x, 1.00x, 1.25x, 1.50x, 1.75x 2.00x, 2.50x, 3.00x, and 4.00x), with 1 pmol of DNA each and an increasing pmol of human recombinant octamer (EpiCypher 16-0001). Each titration was brought to 40 µL volume using a high salt buffer (2M NaCl, 10mM Tris-Cl pH 7.5, 1mM EDTA, 3.5 mM BME, 0.02% NP40) (*75*). Reaction solutions were transferred to mini dialysis tubes (ThermoFisher 69560) for dialysis in 600 mL of high salt buffer. Buffer exchange was achieved using a peristaltic pump that slowly transferred low salt buffer (50mM NaCl, 10mM Tris-Cl pH 7.5, 1mM EDTA, 3.5 mM BME, 0.02% NP40) into the dialysis beaker for 18 hours at 4°C. Nucleosomes were harvested and their presence was confirmed using 6% DNA retardation gel (ThermoFisher EC63655) by loading 10 µL of sample with 2 µL 6X purple loading dye with no SDS (NEB B7024), at 110 Volts, using 0.5X TBE, for 1 hour. The gel was stained with SYBR Safe (ThermoFisher S33102) and imaged.

Cell culture

HEK293T (ATCC® CRL-3216) cells were maintained at 37 °C with 5% CO2 in Dulbecco's Modified Eagle Medium (DMEM) (Corning 10-013-CM) supplemented with 10% fetal bovine serum (FBS) (Clontech Labs 631106), 100 units/mL Antibiotic-Antimycotic (100X) (Thermo Fisher Scientific 15240112) (referred to as DMEM complete). Routine mycoplasma testing was performed monthly.

Optical tweezers conditions and sample capture for unzipping experiments

Imaging was performed using a commercial optical tweezers setup, the C-Trap® (LUMICKS). Calibration of the setup was carried out by analyzing the thermal fluctuations of trapped beads, sampled at 78,125 Hz, with the trapping laser was set to 100% laser power, with overall power at 30% and trap 1 split power at 70%, achieving ~0.2-0.35 pN/nm stiffness for both traps. Before measurements, the laminar flow-based microfluidics system was passivated by flowing bovine serum albumin (BSA; 0.1% wt/vol in phosphate-buffered saline (PBS)) for 30 minutes, followed by a 30-minute PBS flush. Tether formation was performed *in situ* (inside the flow cell) by trapping an anti-digoxigenin bead (bound by DNA handles ligated to CTCF motif-containing construct) in one trap, trapping a ~4.34 µm, streptavidin-coated polystyrene bead (Spherotech SVP-40-5) in the second trap, and bringing the two beads into close proximity to allow binding of the 3×biotin tag in the DNA handles to the streptavidin on the large bead.

Unzipping DNA with optical tweezers

In unzipping experiments, after forming a DNA tether, the distance between the traps was linearly increased at 0.6 µm/s (unless otherwise noted) until the terminal hairpin of the tethered construct was stretched to ~25-30 pN. For CTCF unzipping, the tethered beads were incubated and unzipped in a channel containing 3-10 nM WT, ΔN, or ΔC human CTCF or WT mouse CTCF in 1x CL100 buffer (*19*) (35 mM Tris pH 7.5, 100 mM KCl, 5 mM MgCl2, 5% Glycerol, 0.005% Tween-20, 0.1 mg/mL BSA, 1 mM TCEP) or CL100* buffer (*19*) (35 mM Tris pH 7.5,

50 mM KCl, 50 mM NaCl, 5 mM $MgCl_2$, 5% Glycerol, 0.003% Tween-20, 0.1 mg/mL BSA). No statistically significant differences were observed in rupture forces for the two buffers (K-S test $p > 0.05$).

For unzipping experiments involving cohesin-STAG1, cohesin and STAG1 were pre-incubated at an equimolar ratio for 10 minutes on ice at final concentrations of 50 nM in 1x CL100* buffer. When unzipping CTCF-PDS5A-cohesin-STAG1 or CTCF-NIPBL-cohesin-STAG1, the 50 nM preformed cohesin-STAG1 complex was then additionally pre-incubated with PDS5A and/or NIPBL at an equimolar ratio for 10 minutes on ice at final concentrations of 45 nM in 1x CL100* buffer.

When unzipping CTCF in the presence of cohesin complexes or subunits, the tethered beads were incubated and unzipped in a channel containing 10-20 nM of each component at equimolar ratios in 1x CL100* buffer. When NIPBL was used, the flowed-in buffer and protein mixture was preheated to 37C, constantly replenished, and spiked with 5 mM ATP.

Unzipping specificity control
To quantify the specificity of CTCF on- vs off-targets under unzipping experimental conditions, we cyclically unzipped and rezipped DNA handles ligated to the consensus CTCF construct in a microfluidics channel containing 5 nM CTCF in 1x CL100 buffer and counted the number of peaks observed at the known, expected location and other off-target locations. Percentages were calculated based on the number of unzipping cycles containing, or lacking, each classification of rupture force peak. A total of 159 cycles across 19 replicates were collected from 6 different DNA tethers.

Dynamic measurements using DNA fluctuations
To measure the dynamics within the CTCF-DNA complex, DNA was partially unzipped until it approached the location of the CTCF binding site, which was known from sequence design and prior unzipping experiments. Fluctuation experiments were possible at reproducible, simulatable, sequence-dependent peaks within the naked DNA unzipping signature (Fig. S3), which spontaneously thermally fluctuated between ‘open’ and ‘closed’ states (*52*, *53*).The location and magnitude of these peaks result from the underlying sequence of approximately 20-40 bp (*53*). The applied force was finely adjusted to achieve an approximately equivalent probability of closed versus open configurations and thus position the fork at a nearly identical location between replicates. The degree of fork openness was titrated and verified to not affect CTCF dynamics in Fig. S9.

To measure CTCF dynamics, the construct was moved to a channel containing 3-10 nM WT, ΔN, or ΔC CTCF in 1x CL100. Once protein binding stabilizes the closed state, dynamics were measured for 1-5 minutes. When measuring CTCF dynamics in the presence of cohesin complexes or subunits, the construct was moved to a channel containing 10-20 nM of each component at equimolar ratios in 1x CL100* buffer. When NIPBL was used, the flowed-in buffer and protein mixture was preheated to 37C, constantly replenished, and spiked with 5 mM ATP. Cohesin complexes were prepared as described above for unzipping experiments.

Testing force-dependence of DNA fluctuation dynamic measurements

To determine force was affecting the fraction of time spent in each state, we performed a control experiment to one conducted in a previous work (*53*). This past work observed no force-dependency on ZF association and dissociation in the regime measured using the same DNA fluctuation scheme.

In the experiment, rather than striving for an approximately even fraction of time spent in the open and closed DNA states prior to CTCF binding, we slightly varied the location of the unzipping fork, which altered the average force felt by the pre-bound complex, and thus the initial degree of openness of the consensus motif DNA. We performed 14 replicates ranging from 22% to 85% unbound open DNA fraction (Fig. S9, and Supplementary Methods). The same HMM analysis was then performed on the resulting data to examine the fraction of time spent in each state as a function of the unbound force.

<u>Confocal C-trap measurements</u>

Tracking of CTCF and PDS5A was performed on a LUMICKS C-Trap® optical-tweezers instrument equipped with laminar-flow microfluidics and confocal fluorescence imaging. The continuous-wave tweezers were operated in a dual-trap regime. A five-channel laminar-flow cell was used with the following contents: (i) streptavidin-coated beads, (ii) biotinylated λ-DNA in CL100*, (iii) imaging buffer (CL100* supplemented with an oxygen scavenging system containing 0.8% w/v dextrose, 2mM Trolox (Santa Cruz Biotechnology 53188-07-1), 1mg/ml glucose oxidase (Sigma-Aldrich G2133-50KU), and 500U/ml catalase (Millipore Sigma 9001-05-2)), (iv) CTCF–JF646 (10 nM), and (v) PDS5A–GFP (10 nM), all at room temperature. Two ~4.34 µm streptavidin-coated polystyrene beads (Spherotech SVP-40-5) were trapped, calibrated, and used to capture λ-DNA, which was then stretched to 5 pN and held at constant force. A worm-like chain model was used to verify dsDNA (contour length 16.49 µm for 48.5 kb, persistence length 46 nm, stretch modulus 1000 pN). DNA was then exposed to CTCF. For CTCF-JF646 imaging, confocal settings were: excitation at 488nm (blue) or/and 638 nm (red) pixel size 100 nm, and pixel dwell time was varied between 0.05-0.3 ms. For PDS5A measurements, DNA with pre-bound CTCF was transferred to the PDS5A–GFP channel for two-color imaging.

<u>High-speed atomic force microscopy (HS-AFM):</u>
A commercial Sample-Scanning High-Speed Atomic Force Microscope (SS-NEX Ando model) from RIBM (Research Institute of Biomolecule Metrology Co., Ltd.) was used for experiments involving CTCF and cohesin complexes. Tapping mode was employed to minimize interference with the deposited sample, and all deposited samples were captured in solution. Ultra-Short Cantilevers (NanoAndMore USC-F1.2-k0.15-10), specifically designed for high-speed AFM with a resonance frequency of 1200 MHz, a spring constant of 0.15 N/m, and a length of 7 µm, were purchased from NanoAndMore and utilized in these experiments. A wide scanner was employed with scan speeds ranging from 0.05 to 1 frame per second, with the resolution set to 200 × 200 pixels.

<u>HS-AFM conditions and sample deposition</u>
Sample was prepared for AFM deposition by incubating 0.2 nM DNA substrate (~8 kb or ~1 kb) with combinations of the following proteins, depending on the experiment performed, in SMC buffer (50 mM HEPES pH 7.5, 100 mM NaCl, 10 mM MgCl2) and incubating at 37C for 5 minutes. Protein concentrations are as follows: CTCF (1 nM final), cohesin-STAG1 (1.5 nM

final; pre-incubated as described above), PDS5A (1 nM final), NIPBL (1 nM final), and ATP (5 mM final).

Freshly cleaved mica surfaces (NanoAndMore 50-S-15-15) were coated with 3 uL of polylysine (P-lysine) (Millipore Sigma 25988-63-0) at a concentration of 0.05 mg/mL. After 3 minutes, the surfaces were washed twice with 80 uL water. 3 uL of sample was then added and incubated for 5-10 minutes. Sample was washed once with and imaged in water.

Single-molecule total internal reflection microscopy (smTIRF) imaging setup and preparation

smTIRF was performed using a prism-based Nikon Eclipse Ti microscope. A Shanghai Dream Laser Technology (543nm) solid-state laser was used to illuminate Sytox Orange dyes and fluorescent emission was collected using a water immersion 60X/1.27 NA objective. Laser signal was blocked using a custom filter. An electron-multiplying charge-coupled device (Andor iXon 897) was used to capture images. Movies were recorded using the home-built smCamera 2.0 software.

Lambda DNA TIRF loop extrusion

TIRF microscope slides were prepared and passivated as defined above and previously described (*76*) with methoxy polyethylene glycol (PEG) (Laysan Bio MPEG-SVA-2000-5g) and 2% biotin-PEG (Laysan Bio KIT-BIO-PEG-SVA-5K-mPEG-SVA-5K), but with chambers containing one inlet and two outlets in a configuration that enabled switching between parallel and perpendicular flow. Sample immobilization was achieved by successively flowing, 100 µL of 1x DNA buffer (20 mM Tris pH 7.5, 150 mM NaCl, 0.25 mg/mL BSA) at 1 mL/min, 40 µL 0.2 mg/mL neutrAvidin (ThermoFisher 31000) in 1x T50 buffer (10 mM TrisHCl pH 8.0, 50mM NaCl) at 1 mL/min (1 minute incubation), and 100 µL 1x DNA buffer at 1 mL/min in parallel and perpendicular configurations. 50 µL of 33 pM biotinylated λ-DNA (prepared as described above) was then added into the channel in 1x DNA buffer at 40 µL/min using parallel flow. The chamber was washed once with 40 µL 1x DNA buffer and once with 40 µL 37C 1x IB (40 mM Tris pH 7.5, 30 mM KCl, 2.5 mM MgCl2, 0.5 mg/mL BSA, 1 mM DTT, 250 nM Sytox Orange (ThermoFisher S11368), supplemented with an oxygen scavenging system described above). Immobilized λ-DNA was imaged using perpendicular flow prior to the addition of protein mix. 50 µL of 2 nM pre-incubated Cohesin-STAG1, 2 nM NIPBL, 5 mM ATP in 1x IB at 37C was added at 40 µL/min with perpendicular flow. The sample was then incubated for 2 minutes without flow, and the same field of view was imaged with and without 40 µL/min perpendicular flow using 1x IB at 37C.

CTCF-CUT&RUN

CUT&RUN experiments were performed following the EpiCypher CUT&RUN Protocol (*77*) with some modifications. 0.5-1 million HEK293T cells were washed twice in CUT&RUN wash buffer (20 mM HEPES, pH 7.5, 150 mM NaCl, 0.5 mM spermidine, 1x cOmplete EDTA-free Protease Inhibitor (Roche 11836170001)) and centrifuged at 600 × g at 4°C for 3 minutes. The cells were resuspended in 150 µl of wash buffer and conjugated to 10 µl of Concanavalin A-coated magnetic beads (Bangs Labs BP531). After 10 minutes at room temperature, the supernatant was removed, and the cells were permeabilized on ice with 50 µl of antibody reaction buffer (CUT&RUN wash buffer + 0.01% digitonin + 2 mM EDTA). Five minutes later, 1 µl of CTCF CUT&RUN-grade CTCF antibody (Cell Signaling D31H2) was added, and the mixture was incubated at 4°C overnight.

The material was washed twice with 200 µl of permeabilization buffer (CUT&RUN wash buffer + 0.01% digitonin) and resuspended in 50 µl of the same buffer, after which 2.5 µl of pAG-MNase (EpiCypher 15-1116) was added and incubated at room temperature for 10 minutes. After washing twice with 200 µl of permeabilization buffer, the samples were resuspended in 50 µl of permeabilization buffer, and 1 µl of 100 mM $CaCl_2$ was added on ice to activate the pAG-MNase. pAG-MNase digestion was performed at 4°C for 2 hours. The reaction was stopped by adding 33 µl of 2× CUT&RUN Stop buffer (340 mM NaCl, 20 mM EDTA, 4 mM EGTA, 10 µg/ml RNase A (NEB T3018L), 50 µg/ml glycogen (Milipore-Sigma 361507) and 1 µl of E. coli spike-in DNA (0.5 ng/µl) (EpiCypher 18-1401) to each tube. CTCF-bound DNA was released and collected after incubation at 37°C for 10 minutes.

The DNA was purified using the Monarch Spin PCR & DNA Cleanup Kit (NEB T1130L), eluted in 12 µl of elution buffer, and used for library preparation with the CUTANA™ CUT&RUN Library Prep Kit (EpiCypher 14-1001) according to the manufacturer's instructions. To avoid adapter carryover, gel extraction was performed using the Qiagen Gel Purification Kit (Qiagen 28704). Sequencing was performed sequenced at the Johns Hopkins Genomics Core Facility on an Illumina NextSeq500 or NovaSeq platform (50-bp paired-end). The sequencing data were analyzed using nf-core/cutandrun (*78*).

<u>MNase-seq</u>
0.5 million HEK293T cells were crosslinked with 1% formaldehyde for 5 minutes, quenched with 130 mM glycine, collected, pelleted for 3 minutes at 600 x g, and resuspended in 100 µL of CUT&RUN wash buffer. Pellets were spun down for 3 minutes at 600 g with a total of two washes.
Next, cells were permeabilized with CUT&RUN wash buffer supplemented with 0.01% (v/v) digitonin and incubated on ice for 5 minutes, after which $CaCl_2$ was added to a final concentration of 6 mM. In total, 1.8 U of MNase (NEB M0247S) was added, and the pellets were incubated for 15 minutes at 37°C. The reaction was stopped by mixing it with 100 µL of CUT&RUN Stop buffer. The material was pelleted for 10 minutes at 14,000 g at 4°C, and the supernatant containing nucleosomal DNA was purified using a MiniElute PCR purification kit (Qiagen 28006).

For library preparation, MNase-digested DNA was purified using AMPure XP beads (Beckman Coulter A63881). A total of 500 ng of purified DNA was used for library preparation with the NEBNext Ultra Library Preparation Kit (NEB E7645L), with 9 PCR cycles. Libraries were analyzed and quantified using an E-Gel and Qubit Fluorometer and sequenced at the Johns Hopkins Genomics Core Facility on the Illumina NovaSeq SP100 (50 bp paired-end).

MNase-seq data were mapped to the hg38 genome and processed using an integrated analysis pipeline for MNase-seq, nf-core/mnase (*78*). Nucleosome positions were called using DANPOS2 (*79*) as part of the nf-core/mnase-seq pipeline.

<u>Chromatin GpC methylation, DNA extraction, library prep, and Nanopore sequencing</u>
HEK293T cells were grown as described previously and collected by trypsinization. GpC methylation was performed as described (*40*), with modifications. Nuclei were extracted by incubating on ice for 4 min in lysis buffer prepared 1× RSB (100 mM Tris-Cl pH 7.4, 100 mM NaCl, 30 mM $MgCl_2$) and 0.25% NP-40. Intact nuclei were pelleted (5 min, 1,000 g, 4 °C),

resuspended in 1× RSB, and pelleted again (5 min, 800 g, 4 °C). For labeling, ~$5\times10^5$ nuclei were combined in 500 µL containing 300 mM sucrose, 1× M.CviPI reaction buffer, 96 µM S-adenosyl-methionine (SAM) (NEB B9003S), and 200 U M.CviPI (NEB M0227L). Reactions were incubated at 37 °C, 1,000 rpm for 7.5 min; SAM was replenished to 96 µM at 7.5 min and incubation continued for 7.5 min. Reactions were stopped with 515 µL pre-warmed stop solution (20 mM Tris-Cl pH 7.9, 600 mM NaCl, 1% SDS, 10 mM EDTA). DNA was purified using the Monarch HMW DNA Extraction Kit (NEB T3050L) following the manufacturer's instructions. Purified gDNA was sheared to ~15 kb using a Covaris g-TUBE (Covaris 520079), quantified by Qubit, and taken forward to library prep. Libraries for Oxford Nanopore Technologies (ONT) sequencing were prepared with the LSK110 Ligation Sequencing Kit (ONT, SQK-LSK110) and loaded onto a R9.4.1 pore PromethION Flow Cell (ONT, FLO-PRO002) (~0.5-1 µg sheared gDNA per flow cell) and run on PromethION 24 sequencer (serial PCA100172) for 72 hours. Data were collected using MinKNOW version 23.04.5. Current signals were converted to DNA bases using Guppy v6.5.7 (ONT) in high-accuracy mode (450 bps), yielding 148.37 Gb pass (of 187.07 Gb estimated, 16.14 M reads; N50 ≈ 14.9 kb; total data written 1.47 TB).

Reads were aligned to hg38 with minimap2 and processed with samtools:

```
minimap2 -ax map-ont hg38.fa reads.fastq | samtools view -b - | samtools sort -o sample.sorted.bam && samtools index sample.sorted.bam
```

GpC methylation was called with nanopolish (*80*): nanopolish call-methylation -t 48 -q cpggpc -r reads.fastq -b sample.sorted.bam -g hg38.fa > sample.modcalls.tsv

<u>Motif-centered mapping of ZF single-molecule accessibility</u>

We defined high-confidence CTCF sites by intersecting CUT&RUN peaks (SEACR, stringent mode) with ENCODE CTCF ChIP-seq peaks (hg38 reference). Summits were called independently for each CUT&RUN replicate; sites present in ≥2 replicates were retained (bedtools multiinter). Consensus summits were expanded ±25 bp and overlapped with ENCODE HEK293 CTCF narrowPeak calls (ENCFF314ZAL).To locate exact motif instances, the JASPAR CTCF PWM (MA0139.1) was used to scan summit-anchored sequences with FIMO (*81*) ($p < 1\times10^{-4}$). FIMO coordinates (1-based, inclusive) were converted to strand-aware BED entries at motif centers (respecting the N-to-C orientation of the CTCF motif). Motif calls closer than 36 bp center-to-center were removed.

Single-molecule accessibility was derived from nanopolish per-read GpC calls. Nanopolish calls were intersected with the ±1 kb windows surrounding the motif center (bedtools intersect). To avoid positional ambiguity, we retained singleton k-mers (exactly one GpC per k-mer) and assigned per-read states from the nanopolish log-likelihood ratio (LLR): M (methylated) if LLR ≥ 1, U (unmethylated) if LLR ≤ −1, otherwise N (undetermined). The modified base within each k-mer was realigned to the reference by locating the modified position in the k-mer string and projecting it to genome coordinates. The resulting table contained GpC signals within 22,460 CTCF motifs used for the analysis.

For the protection metaplot (Fig. S1B), we averaged the unmethylated (U) fraction within ±1000 bp of the motif center across the 22,460 motifs without excluding reads. For the ZF accessibility plot (Fig. 1B), the strand-aware offset from the motif center was partitioned into 3-bp bins corresponding approximately to ZF1–ZF11 (per the crystal structures, assuming each ZF interacts with a DNA triplet) (*46*). To remove nucleosome reads covering the CTCF motif, motif×read pairs were excluded if all GpCs in a flanking nucleosome-depleted region (NDR) window (±20±60) were unmethylated (U). We defined a read as "CTCF-bound" if at least one core ZF bin contained an unmethylated GpC (i.e., protected/inaccessible). Using these reads, we computed the per-motif fraction inaccessible for each ZF.

For the conditional-probability analysis (Fig. S1F), nucleosome-excluded reads were subjected to a molecule-level filter requiring ≥2 distinct core ZFs bound (U) per read and reads with all measured ZFs bound were removed to focus on the dynamic subset. We then calculated conditional probability P (ZFj=1| ZFi=1). ZF5 was excluded due to low coverage.

For the correlation analysis, cohesin subunits ChIP-seq signals (downloaded from 4D (*82*) portal (*83*), currently publicly available in Gene Expression Omnibus (https://www.ncbi.nlm.nih.gov/geo/) with the accession number GSE277721 (*57*)) and ENCODE for REST (dataset: ENCFF981ILL) were extracted from normalized bigWig files as the median signal in a ±200 bp window around each motif center. We computed per-motif, per-bin CTCF-bound ZF "binding probabilities" (fraction of reads with U; numerator = reads with any U in the bin; denominator = measured reads in the bin) restricted to dynamic molecules (≥1 bound and ≥1 unbound core ZF across ZF1–ZF11). Binding probabilities were correlated with per-motif ChIP signals for the following ChIP-seq GSE277721 datasets (REST, STAG2, PDS5A, NIPBL, STAG1, RAD21-NIPBL degron 24h, RAD21, CTCF). For each factor and ZF bin (ZF1–ZF11, excluding ZF5) we computed Pearson's r across motifs with raw p-values and controlled the false discovery rate using Benjamini-Hochberg across all factor×ZF tests. ZF5 was excluded from analysis due to low GpC coverage.

Unzipping data processing and analysis

Experimental data were digitized at 2,500 Hz, baseline-corrected, and converted into force and extension vectors using calibration parameters. To precisely adjust and subtract any residual extension offset in individual experiments, force-extension curves up to 10 pN were fit to an extensible worm-like-chain (eWLC) model of double-stranded DNA, converting to DNA base pairs (*52*, *63–65*), aligning the traces as described previously (*65*).The 'bp from the motif center' in force-position curves was calculated by subtracting the known position of the ZF6 interaction with the consensus sequence. (*24*, *46*)

To calculate the mean breaking force in a specific region, we generated an "interaction vector" for each unzipping trace, where an interaction is defined as an increase in force at constant position, followed by a force drop. The interaction with the highest breaking force inside the boundaries of each region defines the region's breaking force, which is then averaged over the ensemble. The significance of differences in breaking forces was assessed using the K-S test, without assuming normality. Applying the same criteria to data obtained without CTCF resulted in no binding events detected. The higher rupture force population was identified using a

Gaussian mixture model and BIC to determine whether a condition was more likely to contain 1 or 2 populations and predict which points fell in each population. 2 populations were more likely for N-terminal WT CTCF+PDS5A (BIC = 525.1 for 1 Gaussian or 455.9 for 2) and WT CTCF+PDS5A+cohesin conditions (BIC = 457.8 for 1 Gaussian or 456.9 for 2).

Force-weighted (FW) dwell time histograms calculated as previously described (*62*, *65*, *84*). Briefly, the number of data points in 1 bp bins was counted and divided by the sampling rate (2500 Hz), multiplied by the detected rupture force, after subtracting the average background signal of naked DNA.

Confocal C-Trap measurement analysis

Data were analyzed using custom python scripts using the Pylake API released by LUMICKS. Kymo tracking feature was used to track the individual traces to obtain the position and time information which was then subsequently used to calculate MSD then diffusion coefficient. Details of which are included in the script submitted in the GitHub.

AFM image processing and analysis

HS-AFM images were viewed and analyzed using the software built by Prof. Toshio Ando's laboratory-built software, Kodec 4.4.7.39, with available source. Tilt and other image correction details are available in the literature (*85*). The contrast was adjusted to enhance the structural features of the images for visualization.

The bp position of proteins colocalized on the substrate DNA was determined via a semi-automated approach. Firstly, the colocalized proteins were annotated using the circle tool in ImageJ. Hereafter, DNA was annotated by a machine-learning model (manuscript in preparation) and lightly dilated to bridge sub-pixel gaps before skeletonization to a 1-px centerline; edge-touching DNA components were excluded. For each protein ROI, the center was snapped to the nearest skeleton pixel and nearby DNA components within a fixed radius were considered as candidates. The along-contour shortest path from the snap point to the nearest DNA end was then computed on the skeleton graph and skipping circular components lacking endpoints. The candidate yielding the shortest path was selected, and path lengths were converted from pixels to nanometers (using the image-specific pixel size) and then to base pairs using a specified nm/bp calibration for the instrument (manuscript in preparation).

AlphaFold 3 modeling

Predicted AlphaFold 3 structures (*50*) of CTCF bound to the double-stranded consensus motif, G-rich strand of the consensus motif, C-rich strand of the consensus motif, double stranded *PDGFRA* motif and G-rich strand of the *PDGFRA* motif in the presence of 11 Zn ions were generated on December 5$^{th}$ 2024. These predicted models were used as starting configurations of the MD simulations. AlphaFold 3 model confidence was assessed using pLDDT and PAE metrics, as reported in Figs. S2K-N and S25. pLDDT (predicted Local Distance Difference Test) plots were generated by averaging atom pLDDT values for each residue providing a per-residue confidence score reflecting how well each residue's position was predicted. PAE (Predicted aligned error) reports the expected positional error between residue pairs after alignment, indicating the reliability of hypothesized protein-protein interactions.

A predicted structure of PDS5A bound to *PDGFRA*-bound CTCF was generated using AlphaFold 3 (*50*) on April 25th, 2025. Amino acid sequences for CTCF (UniProtKB: P49711) and PDS5A (UniProtKB: Q29RF7) were obtained from UniProt. 11 zinc ions were included in the submitted job. Contact interactions were predicted using PyMOL by identifying inter-chain atom-atom pairs located within 3.5 Å.

MD simulations

All MD simulations were performed using NAMD2. (*86*), 2 fs integration timestep and 2-2-6 multiple time stepping. Bsc1, ffSB14 and ZAFF were used for DNA (*87*), proteins (*88*) and ZF (*89*) respectively, TIP3P for water (*90*), with CUFIX corrections to model interactions between ions and nucleic acid/proteins (*91*). The SETTLE algorithm (*92*) was employed to enforce rigidity of covalently bonded hydrogen atoms within water molecules. Additionally, for non-water molecules, the RATTLE algorithm (*93*) was utilized to constrain the motion of hydrogen atoms involved in covalent bonds. The long-range electrostatic interactions were computed using the particle mesh Ewald (PME) scheme over a 1-Å-spaced grid (*94*). VDW and short-range electrostatic forces were evaluated using the 10–12 Å smooth cut-off scheme. Non-bonded interactions were implemented using "1-4 scaling" with a scaling factor of 0.833. NPT (constant number of particles *N,* pressure *P* and temperature *T)* simulations (*95*) were performed using a Nosé-Hoover Langevin piston with a period of 400 fs, decay of 200 fs at a target pressure of 1 atm and Langevin thermostat (*96*) set at 298 K with a 0.5 $ps^{-1}$ damping. Simulations performed in the NVT (constant number of particles *N,* volume *V* and temperature *T)* ensemble employed the Langevin thermostat. Energy minimization was carried out using conjugate gradients (*97*).

Predicted AlphaFold 3 structures were converted to a PDB format using VMD (*98*) and used as starting configurations. The intrinsically disordered domains of CTCF (residues 1-265 and 585-727) were truncated, and only the zinc finger-DNA complex was simulated. The resulting construct avoids confounding, non-specific interactions from the disordered tails while preserving Zn finger interactions, geometry and topology. The protein was protonated at pH 7 using H++ (*99*). The PDBs were converted to Amber format, solvated and ionized to produce a neutral system with 0.15 M KCl using the tLEaP package of AmberTools22 (*100*). Atomic coordinates were saved every 18 ps. VMD (*98*) and MDAnalysis (*101*) were used for visualization and analysis. Each system was minimized for 20,000 steps and equilibrated in the NPT ensemble for 4.6 ns. During minimization and NPT equilibration, harmonic restraints with a spring constant of 5 kcal/ (mol $Å^2$) were applied to non-hydrogen atoms of nucleic acid and protein. The production simulations (2 replicas of each system), each spanning 1 µs, were performed within an NVT ensemble, devoid of any restraints.

Hydrogen-bond analysis: DNA-protein hydrogen bonds were identified using a donor - acceptor distance cutoff of 3.5 Å and an angle within 30° of linearity (donor-hydrogen-acceptor).

Dynamic trace preprocessing and formatting

The dynamic traces were prepared for HMM state determination by first cropping the sections containing CTCF binding, followed by linear correction to account for experimental drift. Finally, intercepts were aligned to remove baseline variability, ensuring consistent state identification and improved HMM parameter estimation (Fig. S28). The number of base pairs unzipped was calculated by subtracting the force-dependent extension of the handles and then dividing by twice the extension of a single ssDNA base at the given tension as described previously (*52*, *53*).

Hidden-state model selection and cross-condition consensus

All analyses were performed in Python 3.9 using the package hmmlearn (for HMM).

*Model fitting:* Force-time trajectories from individual molecules were first concatenated after removing invalid values. These one-dimensional force series were then modeled with Gaussian-emission HMMs, in which each hidden state is associated with a Gaussian distribution of forces and transitions between states follow a Markov chain. For each candidate number of hidden states (from 1 up to 5), a separate HMM was trained by expectation-maximization (EM). The algorithm iteratively adjusted transition probabilities and emission parameters until the improvement in model fit was smaller than a set tolerance ($10^{-2}$) or a maximum of 300 iterations was reached. For every fitted model, we recorded the overall log-likelihood (how well the model explained the observed data), as well as information-theoretic model selection criteria.

*Model selection criteria:* To decide how many states best described the data, we compared models using two standard information criteria: the Akaike Information Criterion (AIC) and the Bayesian Information Criterion (BIC). Both criteria balance model fit against model complexity, penalizing models with more parameters unless they provide a substantially better fit. In addition, we examined the incremental log-likelihood gain when adding an extra state. This allowed us to identify when increasing the number of states yielded large improvements (as for three to four states) versus when improvements were marginal (as for four to five states).

*Per-condition evaluation and consensus determination:* For each experimental condition, models with up to five states were fitted, and the number of states minimizing the BIC was identified. To obtain a global consensus across conditions, BIC curves were aligned by state number and averaged, and the consensus state number was defined as the one that minimized this mean BIC. In addition, we examined the mean BIC improvement between successive state numbers. Following Kass and Raftery, we defined an “elbow” at the point where the average BIC improvement dropped below 10 units, indicating that adding more states no longer provided meaningful explanatory power (Fig. S6).

*Application to our dataset:* Applying this approach, eight of nine experimental conditions reached their minimum BIC at four states, including the consensus sequence, *PDGFRA* N-terminal CTCF WT, *PDGFRA* N-terminal CTCF WT with PDS5A, methylated PDGFRA N-terminal CTCF WT, nucleosome-associated C-terminal CTCF WT, *PDGFRA* C-terminal CTCF WT, and *PDGFRA* N-terminal CTCF ΔN. The mean BIC across all conditions was likewise minimized at four states, establishing this as the consensus model size. Only one condition - *PDGFRA* N-terminal CTCF WT + Cohesin - showed a further decrease in BIC when a fifth state was added. The log-likelihood gain from four to five states was modest (~155 units), but still sufficient to exceed the BIC penalty, yielding a ΔBIC of ~136, but the improvement was negligible compared to the large increase observed when moving from three to four states (~83,000 units). We therefore decided to use four states, as that best captures the data across conditions (Fig. S6).

*Final model fitting:* Once the consensus number of states was determined, we refit the final HMMs for each condition on the full dataset using hmmlearn. Models were initialized with diagonal covariances, quantile-spaced means, and a sticky transition prior (70% self-transition probability). A sticky transition matrix makes the HMM conservative - it only changes state when the data strongly support it. This produces more stable state assignments and avoids overinterpreting noise as new states. Start probabilities were set uniformly, and automatic reinitialization was disabled. Models were trained for up to 1000 EM iterations or until convergence (tolerance $10^{-3}$).

Transition rate and energy landscape calculations, and visualization

The free energy landscape (Fig. S17) is constructed by determining the well heights ($G_i$) and barrier heights ($G_{ij}^{\ddagger}$) from transition probabilities (Fig. S10) and state lifetimes (Table S2) assuming thermal equilibrium (*102–104*). These calculations rely on equilibrium relationships and transition state theory to estimate the relative free energies of states and activation barriers. To compute the free energy of each state, the equilibrium constant between states $i$ and $j$ is determined from the forward and reverse rate constants, defined in equation 1.

$$K_{ij} = \frac{k_{ij}}{k_{ji}}$$

(equation 1)

Where $k_{ij}$ represents the rate constant for the transition from state $i$ to state $j$, and $k_{ji}$ represents the rate constant for the reverse transition. The rate constants are derived from the transition probabilities ($P_{ij}$) and state lifetimes ($\tau_i$) where the total transition rate out of a state is given by equation 2. Here, the transition probabilities exclude self-transitions / the probability of not transitioning to another state. Errors on the transition rate are calculated from error propagation of $P_{ij}$ and state lifetimes $\tau_i$. Where the error on $P_{ij}$ is calculated as the standard deviation between replicates and the error on $\tau_i$ is estimated as the error from fitting an exponential CDF.

$$k_{ij} = \frac{P_{ij}}{\tau_i}$$

(equation 2)

Using the equilibrium constant, the free energy difference between states $i$ and $j$ is computed as given in equation 3.

$$\Delta G_{ij} = -k_B T ln\,(K_{ij})$$

(equation 3)

Where $k_B$ is the Boltzmann constant and $T$ is the absolute temperature. The free energy of each state is then calculated iteratively relative to a reference state, typically setting $G_1 = 0$, such that the relation stands as in equation 4.

$$G_j = G_i + \Delta G_{ij}$$

(equation 4)

Barrier heights are computed relative to the well energies by considering the activation free energy required for a transition between states. The forward barrier height is determined from the rate constant as given in equation 5.

$$G_{ij}^{\ddagger} = G_i - k_B T ln\left(\frac{k_{ij}}{\nu_{ij}}\right)$$

(equation 5)

Where $\nu_{ij}$ represents the attempt frequency, an intrinsic property of the system characterizing how often transitions are attempted. Although $\nu_{ij}$ would vary between conditions, as $\nu_{ij}$is unknown, we used the same value for all transitions and conditions, approximated by the Eyring prefactor (*105*) ($\nu = k_B\, T/h$). Similarly, the reverse barrier height is computed as given in equation 6.

$$G_{ji}^{\ddagger} = G_j - k_B T ln\left(\frac{k_{ji}}{\nu_{ji}}\right)$$

(equation 6)

The energy landscape is visualized by plotting the state energies and transition barriers along a continuous reaction coordinate. All energies are expressed relative to the reference state I, which is set to zero. The energy levels ($G_i$) are positioned at discrete state indices, while the barrier heights ($G_{ji}^{\ddagger}$) are placed at intermediate positions between adjacent states. To ensure a smooth representation of the energy landscape, the energy values are interpolated using piecewise cubic Hermite interpolation (PCHIP). This method preserves the shape of the landscape without introducing artificial oscillations. To facilitate this interpolation, additional artificial data points are introduced at the edges of the landscape with elevated energy values, ensuring a visually consistent upward curvature at the boundaries. The interpolated curve is then evaluated at finely spaced reaction coordinates, generating a continuous trajectory that represents the transition between states.

<u>Trace categorization for PDS5A and cohesin traces by HMM model selection</u>

Due to the nature of the optical tweezers experiment, we could not verify which additional proteins were present on CTCF-bound-DNA upon addition of PDS5A or cohesin or PDS5A and cohesin. To categorize how well individual traces correspond to different protein compositions, we constructed HMMs for each experimental condition and compared their ability to explain the observed data. As a common starting point, we used a reference HMM trained on pure CTCF traces, which defined the emission distributions for four conformational states of the DNA-protein complex. The emissions (Gaussian distributions of the measured force signal) were kept fixed across all subsequent analyses to ensure that the state identities remained consistent across conditions. Meaning that state I always represented the highest-force, fully bound state, while state IV represented unbound DNA-like fluctuations.

For each condition beyond CTCF, we adapted the kinetics of the reference model using all available traces for that condition. Each trace was decoded using the Viterbi algorithm, producing a most-likely state sequence. From these sequences, we tabulated the number of times each state was visited first, and the number of observed transitions between every pair of states.

These counts were converted into probabilities by normalizing over the total dwell time per state, with a small smoothing factor added to avoid zeros. In this way, we obtained condition-specific estimates of the transition matrix and starting probabilities, while keeping the emission distributions fixed. The result was a new HMM for each condition that retained the same state definitions as the CTCF reference but differed in its dynamical parameters. For stability, these kinetics-adapted models could optionally be further refined by a short round of expectation-maximization, which adjusted only the start and transition probabilities.

Once these models were built, every trace was evaluated by computing its log-likelihood under each of the models deemed biologically plausible for that dataset. For a given observed trajectory $X = \{x_1, \ldots, x_T\}$, the HMM likelihood is defined in equation 9.

$$p\,(X|M) = \sum_{z_1,\ldots,z_T} \pi_{z_1} b_{z_i}\,(x_1) \prod_{t=2}^{T} a_{z_{t-1},z_t} b_{z_t}\,(x_t)$$

equation 9

Where $\pi$ are the starting probabilities, $a_{ij}$ the transition probabilities, and $b_j\,(x)$ the Gaussian emission densities for state j. Because the emissions are identical across models, differences in likelihood reflect only how well the observed temporal patterns of switching agree with the condition-specific kinetics. Each trace was assigned to the model that maximized the log-likelihood. In addition, we calculated the difference in log-likelihood between the selected model and the pure CTCF model ($\Delta LL$). This provided a measure of how strongly the trace deviated from CTCF-like behavior and therefore how confidently it could be attributed to the presence of PDS5A, cohesin, or both. Through this procedure, every trace was categorized into the regime that best explained its dynamics, while ensuring that state definitions were preserved across all conditions. This strategy allowed us not only to separate traces into CTCF-only versus factor-bound regimes, but also to quantify how strongly each factor altered the kinetics of DNA-protein interactions. In the Results, we use this framework to show that traces exhibiting factor-specific kinetic signatures can be robustly distinguished from those consistent with CTCF alone.

### Supplementary Text

Fitting of lifetimes of CTCF-DNA binding using MLE

The lifetime distributions of states of CTCF binding to DNA that were best represented by more than a single exponential function were analyzed using unbinned MLE. A three-binding-state model, represented by a triple-exponential distribution (equation S1), was determined to best describe the binding behavior of the binding behavior of the high force state (state 2) of CTCF when assuming at 2-state HMM analysis (Fig. S5). The fitting parameters were optimized iteratively until the likelihood function converged, ensuring the best-fit parameters and model.

$$P_{t_i} = \frac{W_1}{\tau_1} \times exp\left(-\frac{t_i}{\tau_1}\right) + \frac{W_2}{\tau_2} \times exp\left(-\frac{t_i}{\tau_2}\right) + \frac{W_3}{\tau_3} \times exp\left(-\frac{t_i}{\tau_3}\right)$$

(equation S1)

Temporal limitations of the measurement and analysis

The use of optical tweezers is associated with some level of noise due to the Brownian motion of the beads. To estimate the timescale of Brownian motion for a 2 µm bead (the bead we move in our unzipping experiments) in an optical trap with a stiffness of 0.2 pN/nm at room temperature, we calculate the characteristic relaxation time (τ). This calculation is seen in equation S2 and S3, where $\gamma$ is the drag coefficient (viscous resistance), k is the trap stiffness, $\eta$ is the dynamic viscosity and R is the radius of the bead.

$$\tau = \gamma / k$$

(equation S2)

$$\gamma = 6\pi\eta R$$

(equation S3)

This timescale reflects the bead's ability to relax back toward equilibrium after being displaced by thermal fluctuations and ends up being approximately 94 $\mu$s. Force variations below this temporal resolution might therefore not reflect changes in the CTCF-DNA complex but rather the instrument. We therefore conservatively downsample our data to a 500 $\mu$s to decrease effects from Brownian motion. Consequently, there might exist more than four states, especially given the fact that there are eleven ZFs. However, at this temporal resolution we find that there are four statistically backed states (when CTCF is bound). One could imagine that additional states have transitions faster than this temporal resolution at lower force variation than the experimental noise.

Force dependency of CTCF-DNA complex

We propose a theoretical model in which the CTCF protein experiences the same force as the DNA sequence, as measured by the optical tweezers. Under this assumption, the energy landscape of the four observed CTCF states is tilted by the applied force. For this analysis, we focus exclusively on the first three states (state I, state II, and state III), as they are associated with double-stranded DNA (dsDNA) binding. We neglect state IV ($P_{4,consensus}$= 0.02%), in the model.

First, the free energy difference between adjacent states at zero force must be determined. When a force $F$ is applied, the energy landscape is tilted by introducing a linear term proportional to $F$ and the distance $x$ between the states (Bell's model[1056]:

$$\Delta G_{I\to II}\ (0) = \Delta G_{I\to II}\ (F) + F\Delta x_{I\to II}\ , \Delta G_{I\to III}\ (0) = \Delta G_{II\to III}\ (F) + F\Delta x_{II\to III}$$

(equation S4)

Here $F\Delta x_i$ represents the work done by the applied force on the system to transition into state $i$ and $\Delta G_i\ (0)$ is the free energy of state $i$ at zero force. To calculate the energy difference between adjacent states at a given force $F$, we use the Boltzmann relation, where $P$ denotes the occupancy of a particular state:

$$\Delta G_{1\to 2}\ (F) = -k_B T ln\ (P_2\ (F)/P_1\ (F))\ , \Delta G_{2\to 3}\ (F) = -k_B T ln\ (P_3\ (F)/P_2\ (F))$$

(equation S5)

To calculate the theoretical force-dependent occupancies, we employed a formula derived from the Boltzmann distribution, which states that the probability of the system being in a particular state $i$ at equilibrium is given by:

$$P_i = \frac{e^{-\Delta G_i/k_B T}}{\sum_j\ e^{-\Delta G_j/k_B T}}$$

(equation S6)

Where $P_i$ is the probability of state $i$, $\Delta G_i$ is the free energy of state $i$ (relative to a reference state, chosen as state I). For a three-state system, state I is the reference $(\Delta G_I = 0)$, state II has free energy difference $\Delta G_{I\to II}$ and state III has free energy difference $\Delta G_{I\to II} + \Delta G_{II\to III}$. Substituting equation S4 into the Boltzmann formula (equation S6) and simplifying:

$$P_1\ (F = 0) = \frac{1}{1 + e^{-\Delta G_{1\to 2}\ (0)/k_B T} + e^{-\ (\Delta G_{1\to 2}\ (0) + \Delta G_{2\to 3}\ (0))/k_B T}}$$

$$P_2\ (F = 0) = \frac{e^{-\Delta G_{1\to 2}\ (0)/k_B T}}{1 + e^{-\Delta G_{1\to 2}\ (0)/k_B T} + e^{-\ (\Delta G_{1\to 2}\ (0) + \Delta G_{2\to 3}\ (0))/k_B T}}$$

$$P_3\ (F = 0) = \frac{e^{-\ (\Delta G_{1\to 2}\ (0) + \Delta G_{2\to 3}\ (0))/k_B T}}{1 + e^{-\Delta G_{1\to 2}\ (0)/k_B T} + e^{-\ (\Delta G_{1\to 2}\ (0) + \Delta G_{2\to 3}\ (0))/k_B T}}$$

(equation S7)

To estimate parameters for the model, from the HMM fit of the consensus sequence data (n = 9), we extracted the average occupancy (fraction of time spent, Fig. S15, Table S2) of each state. Assuming the protein feels the same force as the pre-bound unzipping fork, we also extracted the average force (post-linear correction as explained in main methods) before binding ($F = 12.09\ pN$). Here, the occupancies of state 1-3 were normalized to add up to 100%, as we do not consider state 4 (see equation S8):

$$P_1 = \frac{P_1}{P_1 + P_2 + P_3}, P_2 = \frac{P_2}{P_1 + P_2 + P_3}, P_3 = \frac{P_3}{P_1 + P_2 + P_3}$$

(equation S8)

To create experimental curves (Fig. S9), we used the experimentally measured average occupancies (at $F = 12.09\ pN$) as initial conditions to calculate the theoretical occupancies of each state in a range of potential forces.

To experimentally investigate the effect of applied force on the occupancies of the states, we used a similar approach to Khamis et al., NAR, 2021 (*53*). By changing the initial, unbound ratio of open and closed DNA, we changed the average force felt by the pre-bound unzipping fork. We then measured state occupancies under these different applied forces. We tested unbound, open DNA (Fig. S5) percentages ranging from 22% to 85%. For each replicate, we plotted the occupancies of each CTCF state as a function of the average, pre-bound applied force. To calculate the errors for the occupancies, we split the bound portion of each trace into ten chunks (n = ~19,000 data points / chunk or ~9.5s / chunk), determined the state occupancies within each chunk, and calculated the standard deviation between them. The errors for applied force were calculated as the standard error of the pre-bound force (n = ~30,000 data points / trace or ~15s / trace).

We then compared the fit of the data to the theoretical, force-dependent model to a force-independent model, where the occupancy of each state is constant in the range of applied forces. The constant force was set to the same average force used as the initial condition in the theoretical model. To investigate which of the two models, force-dependent or force-independent, better fit the data, we calculated AIC and BIC values, which took into account the error on both the occupancy and the applied force. By summing the log-likelihood contributions for all three states, the model comparison treats the system as a whole, considering how well it explains the data for each state simultaneously. Both criteria favored the force-independent model over the theoretical, force-dependent model.

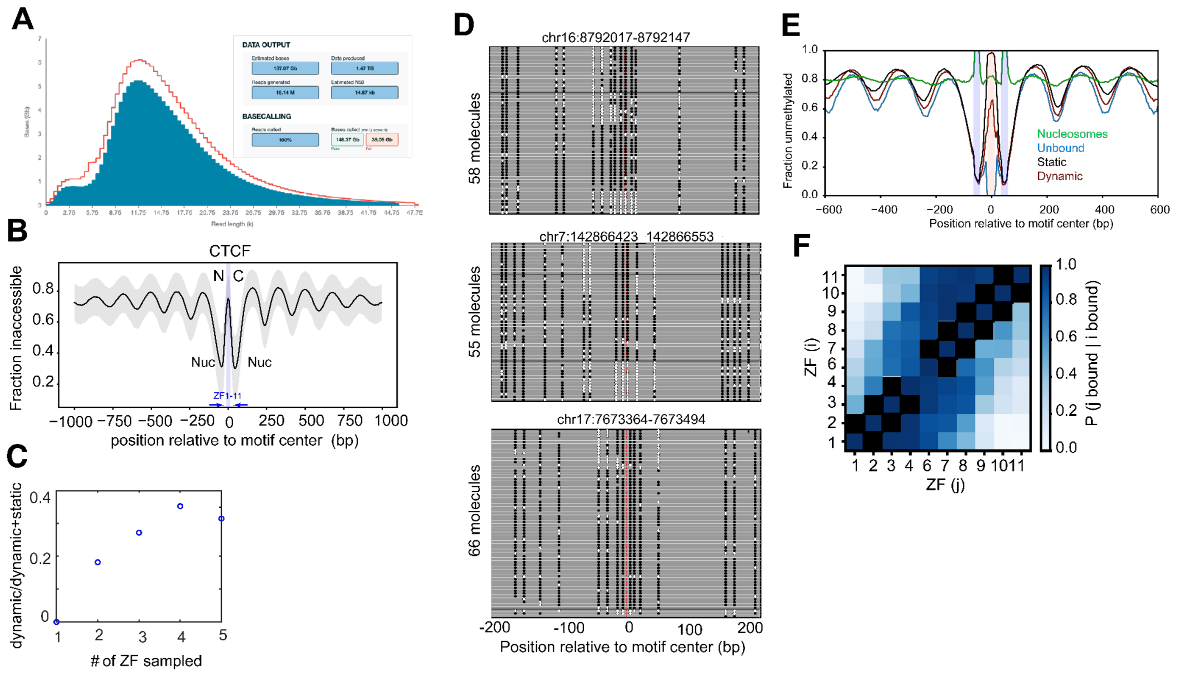


**Fig. S1. CTCF and local chromatin structure probed by GpC methyltransferase (MTase) footprinting followed by nanopore sequencing. (A)** Summary of the PromethION run; read-length distribution (N50) and run statistics are shown. Red-predicted total bases prior to basecalling. Blue- corresponds to the actual basecalled read lengths, **(B)** Mean fraction of inaccessible bases within ±1 kb of CTCF motifs, revealing well-positioned nucleosome arrays. The shaded gray area depicts the standard deviation. Blue rectangles denote the locations bound by CTCF ZFs. **(C)** Fraction of dynamic CTCF molecules among CTCF-bound reads plotted as a function of the number of ZF sampled in our dataset. **(D)** Representative loci overlapping CTCF motifs that illustrate distinct accessibility patterns. Black circles indicate unmethylated GpCs, white circles indicate methylated GpCs, and grey boxes indicate positions where methylation is not possible (no GpC) or uncalled. **(E)** Reads were stratified by motif-region accessibility (see Materials and Methods) into nucleosomal (green), unbound (blue), static (black), and dynamic (red) classes and plotted as mean aggregate profiles across a 600-bp window centered on the motif. Unbound reads lack the ‘array-like’ pattern, consistent with absence of CTCF binding. Static and dynamic reads exhibit a nucleosome-free region at the motif with flanking nucleosomes shifted away relative to unbound DNA. Inset: fraction of dynamic molecules as a function of ZFs probed. **(F)** Conditional probability for a subset of partially accessible molecules of ZF j being bound (unmethylated GpC within the triplet) given that ZF i is bound (Materials and Methods). N (motifs): ZF1=2,229, ZF2=559, ZF3=2125, ZF4=6050, ZF6=2544, ZF7=2768, ZF8=2966, ZF9=984, ZF10=1457, ZF11=1265.

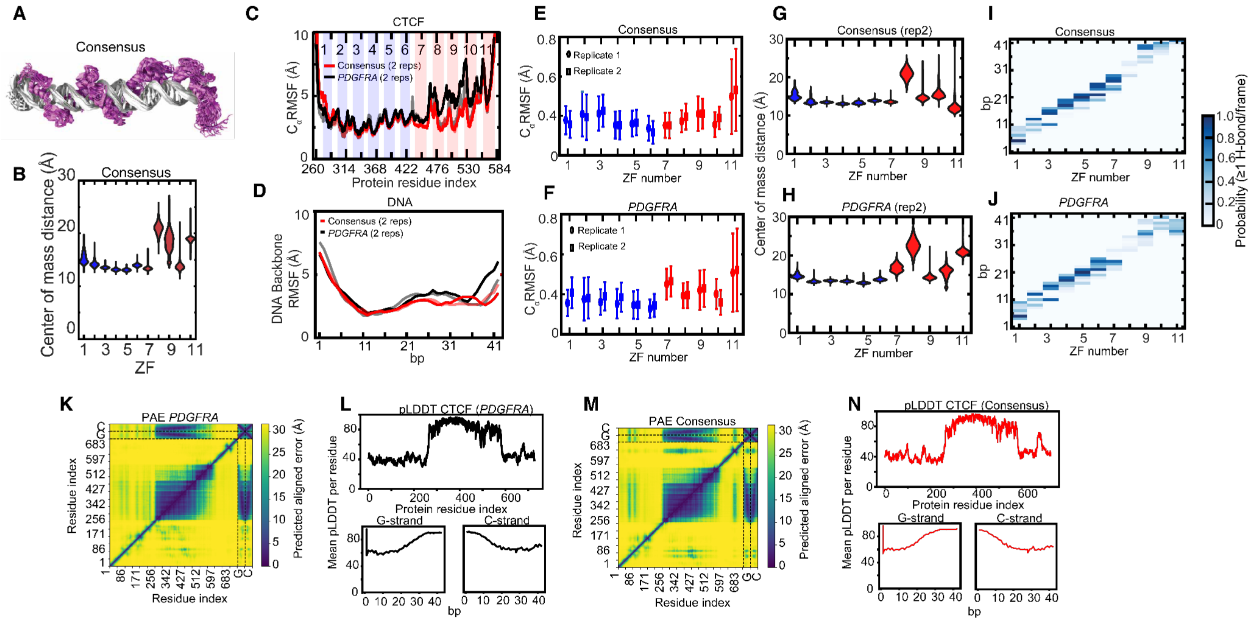


**Fig. S2. MD simulations of IDR-truncated CTCF bound to the consensus or *PDGFRA* sequence. (A)** Multiple overlaid snapshots (every 40 ns) of CTCF bound to its consensus motif sampled by 1 μs molecular dynamics (MD) simulations. **(B)** Simulated distribution of distance between the center of mass of each ZF and its 3 nearest DNA bp for each ZF in the consensus sequence. N-terminal ZFs (1-6) are shown in blue and C-terminal ZFs (7-11) are shown in red. A second replicate is shown in Fig. S2G. **(C)** Root mean square fluctuation (RMSF) of the CTCF C$\alpha$ coordinates with respect to its initial structural model, averaged over the microsecond MD trajectory of the respective system. Prior to each RMSF calculation, the protein coordinates were aligned with the initial structural model. Data are shown for the two replicate simulations of the consensus (red) and *PDGFRA* (black) DNA substrate systems. **(D)** Per base pair backbone RMSF of the DNA substrate with respect to its initial structural model, averaged over the microsecond MD trajectory of the respective system. Data are shown for the two replicate simulations of the consensus (red) and *PDGFRA* (black) DNA substrate systems. **(E-F)** C$\alpha$ RMSF of each ZF, computed by aligning each ZF individually to its initial structural model. Data are shown for the two replicate simulations (ovals and squares) of, **(E)** consensus or **(F)** *PDGFRA* DNA substrate system. N-terminal ZFs (1-6) are shown in blue, while C-terminal ZFs (7-11) are shown in red. **(G-H)** Center-of-mass distance between each ZF and its three nearest DNA base pairs in the **(G)** consensus or **(H)** *PDGFRA* DNA construct, sampled by the respective MD simulations. N-terminal ZFs (1-6) are shown in blue and C-terminal ZFs are shown in red. The first replicate for *PDGFRA* and consensus are shown in Fig. 1D and Fig. S2B, respectively. **(I-J)** Probability of observing one or more hydrogen bonds between a ZF and a DNA bp during MD simulations of the **(I)** consensus or **(J)** *PDGFRA* system. The statistical information in (**I-J**) combines data from the two replicate simulations. **(K)** Predicted aligned error (PAE) plot of the AlphaFold 3 model of CTCF bound to the *PDGFRA* motif, used as the starting point for the MD simulation. Dashed lines separate the different chains of the model (CTCF, *PDGFRA* "G-strand", *PDGFRA* "C-strand"; where the strands are named based on the prominent nucleotide base in the consensus sequence on that strand). **(L)** Plots of the per residue mean predicted local distance difference test (pLDDT) value for each chain of the AlphaFold 3 model of CTCF bound to the *PDGFRA* motif: CTCF (top), *PDGFRA* G-strand (bottom left), *PDGFRA* C-strand (bottom right). **(M)** PAE plot of the AlphaFold 3 model of CTCF bound to the consensus motif, used as the starting point for the MD simulation. Dashed lines separate the different chains of the model (CTCF, consensus “G-strand”, consensus “C-strand”). **(N)** Plots of the per residue mean pLDDT value for each chain of the AlphaFold 3 model of CTCF bound to the consensus motif: CTCF (top), consensus G-strand (bottom left), consensus C-strand (bottom right).

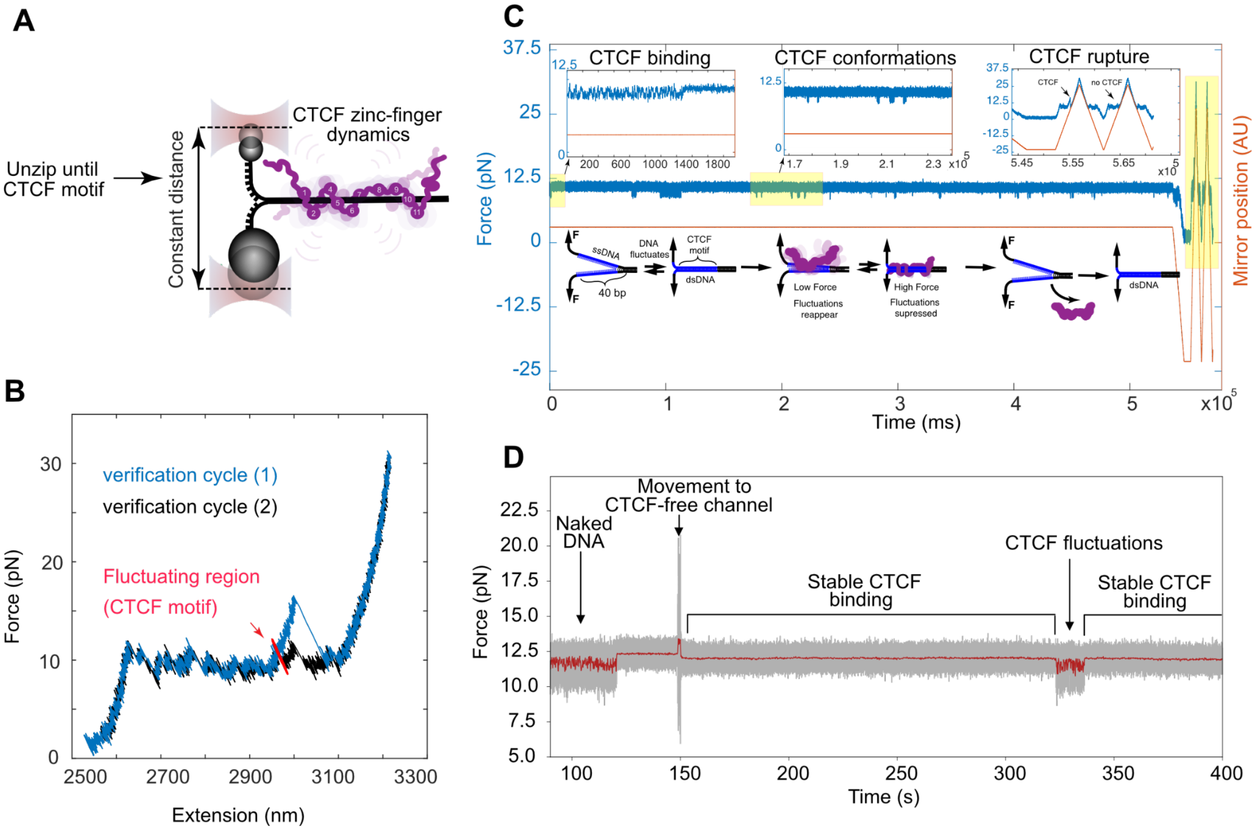


**Fig. S3. Dynamic CTCF measurements in a CTCF-free channel. (A)** A cartoon of CTCF dynamic measurements where the DNA handles are unzipped up to the motif and held at constant position to track thermal breathing of the DNA fork. **(B-C)** A representative dynamic CTCF measurement and subsequent unzipping validation shown as a **(B)** force-extension curve and **(C)** force over time. Additional information on the unzipping scheme can be found in Fig. S4. Upon CTCF binding, DNA transitions from two-state breathing to a higher-force state followed by subsequent appearance of CTCF dynamics (as shown in the left and middle insets of **(C)**). The constant-position dynamics appear as fluctuations in force and DNA extension in the force-extension plot in **(B)** (red). At the conclusion of the dynamic measurement, the DNA was unzipped twice to validate CTCF binding (as seen in the right inset of **(C)**). In the first unzipping cycle (blue in **(B)**) a CTCF rupture peak was observed. In the following unzipping cycle (black in **(B)**) CTCF had dissociated. In **(C)**, a cartoon depicts the experimental scheme. **(D)** A representative trace shows stabilization of force at a higher force upon binding of CTCF to DNA. The stark destabilization in force at 150s is an artifact of movement of the beads to a CTCF-free channel, followed by returning to the previous stably bound CTCF signature. The slight decrease in force pre- and post-movement is often observed and likely due to flow cell differences. Around 330 s we see momentary fluctuations shortly followed by stable CTCF binding again around 340 s.

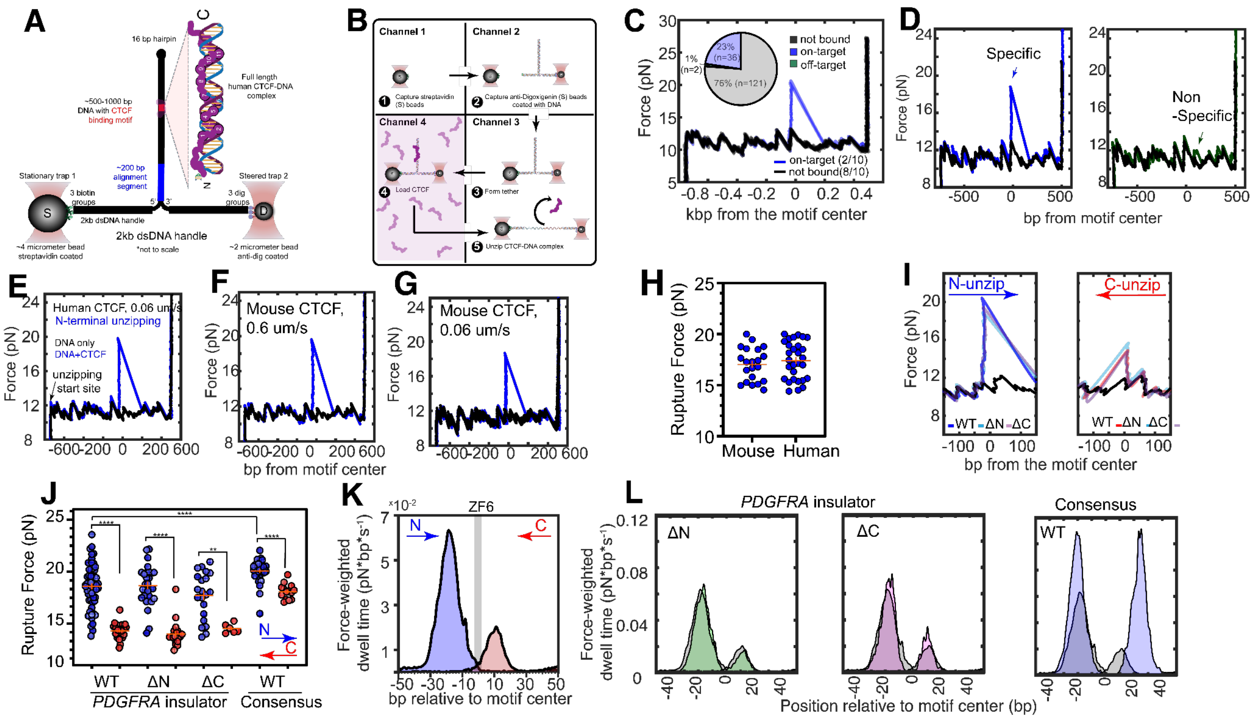

**Fig. S4. Unzipping of CTCF constructs with optical tweezers. (A)** Shown is the unzipping construct attached to functionalized "DNA handles" and modified polystyrene beads trapped in highly focused lasers (optical traps). The ends of the handles contain 3 terminal biotin or 3 terminal digoxigenin, which are able to bind the streptavidin-coated or anti-digoxigenin-coated beads. The unzipped portion of the handles contain an alignment region (blue) for computational correction (Materials and Methods), a CTCF motif (red), a terminal hairpin (black dot), and additional DNA computationally designed to exclude CTCF binding sites (black). Sizes of components are noted. An inset of the red CTCF binding site shows a cartoon of bound, human CTCF. **(B)** A schematic of a typical unzipping scheme within a LUMICKS C-Trap® flow cell. First, a streptavidin-coated bead is captured in an optical trap in channel 1. Next, an anti-digoxigenin-coated bead, pre-incubated with the unzipping construct, is captured in the second trap in channel 2. In a buffer-only channel 3, single DNA tethers are formed by bringing the beads into proximity to each other. The construct is then incubated in channel 4 containing a protein (s) of interest and brought back to the buffer-only channel 3 for unzipping. **(C)** A representative force-extension plot of a DNA construct containing the consensus motif being repetitively unzipped and rezipped from the N-terminal orientation. All cycles are overlaid, 8 of which were unbound cycles (black) and 2 of which were specifically-bound by CTCF (blue). The inset pie chart shows counts of cycles that were unbound (gray), specifically-bound (blue), or nonspecifically bound (green) by CTCF in 19 independent cyclical unzipping traces containing a total of 159 cycles. **(D)** Representative traces illustrating the interpretation of specific (left, blue) vs nonspecific (right, green) CTCF binding. The molecule classified as off-target has a lower force rip and is about 80 bp away from the CTCF consensus motif. **(E-G)** Representative unzipping traces of DNA bound by **(E)** human or **(F)** mouse CTCF unzipped at 0.6 µm/s, where the rupture forces and unzipping profiles look comparable to slower unzipping at 0.06 µm/s (as shown in **(G)** for mouse CTCF). Representative unzipping of unbound DNA is shown in black at the noted speeds. **(H)** Unzipping rupture forces of mouse and human CTCF unzipped at 0.6 µm/s. The mean rupture forces ± standard error are mouse: n = 21, F = 17.02 ±

0.35 pN; human: n = 29, F = 17.41 ± 0.34 pN. **(I)** Representative unzipping traces at 0.6 μm/s from the N- or C-terminal orientation of WT, ΔN, and ΔC CTCF on *PDGFRA* or consensus DNA sequences, as noted. See keys in both panels for the color scheme. **(J)** Single-molecule rupture forces for CTCF-DNA interactions on the *PDGFRA* or consensus motif for WT, ΔN, and ΔC CTCF from the N-terminal (blue) and C-terminal (red) orientations. The mean rupture forces for the *PDGFRA* motif were 18.6 ± 0.3 pN (n = 57), 18.8 ± 0.4 (n = 23), 17.7 ± 0.5 (n = 21) for WT, ΔN, and ΔC, respectively, from the N-terminal orientation and 14.3 ± 0.1 (n = 31), 14.0 ± 0.3 (n = 19), 14.5 ± 0.2 (n = 6) for WT, ΔN, and ΔC, respectively, from the C-terminal orientation. The mean rupture forces for WT CTCF on the consensus motif were 20.1 ± 0.2 pN (n = 34), 18.1 ± 0.2 (n = 14) for the N- and C-terminal orientation, respectively. Data are mean ± standard error. Using a 1-D, two-state Kolmogorov-Smirnov (K-S) test, significance is indicated as follows: **p<0.01, ****p<0.0001. **(K)** Force-weighted dwell time map of CTCF bound to the *PDGFRA* motif as a function of bp position. Interactions probed from the C-terminal orientation are shown in red, while interactions probed from the N-terminal orientation are shown in blue. The locations of ZFs are overlaid as gray bars, assuming each ZF binds a single DNA triplet. **(L)** Force-weighted dwell time maps of ΔN (green) and ΔC (purple) CTCF bound to the *PDGFRA* insulator and WT CTCF bound to the consensus sequence (blue). They are all overlaid over the WT CTCF-*PDGFRA* (gray) force-weighted dwell time map previously shown in **(K)**.

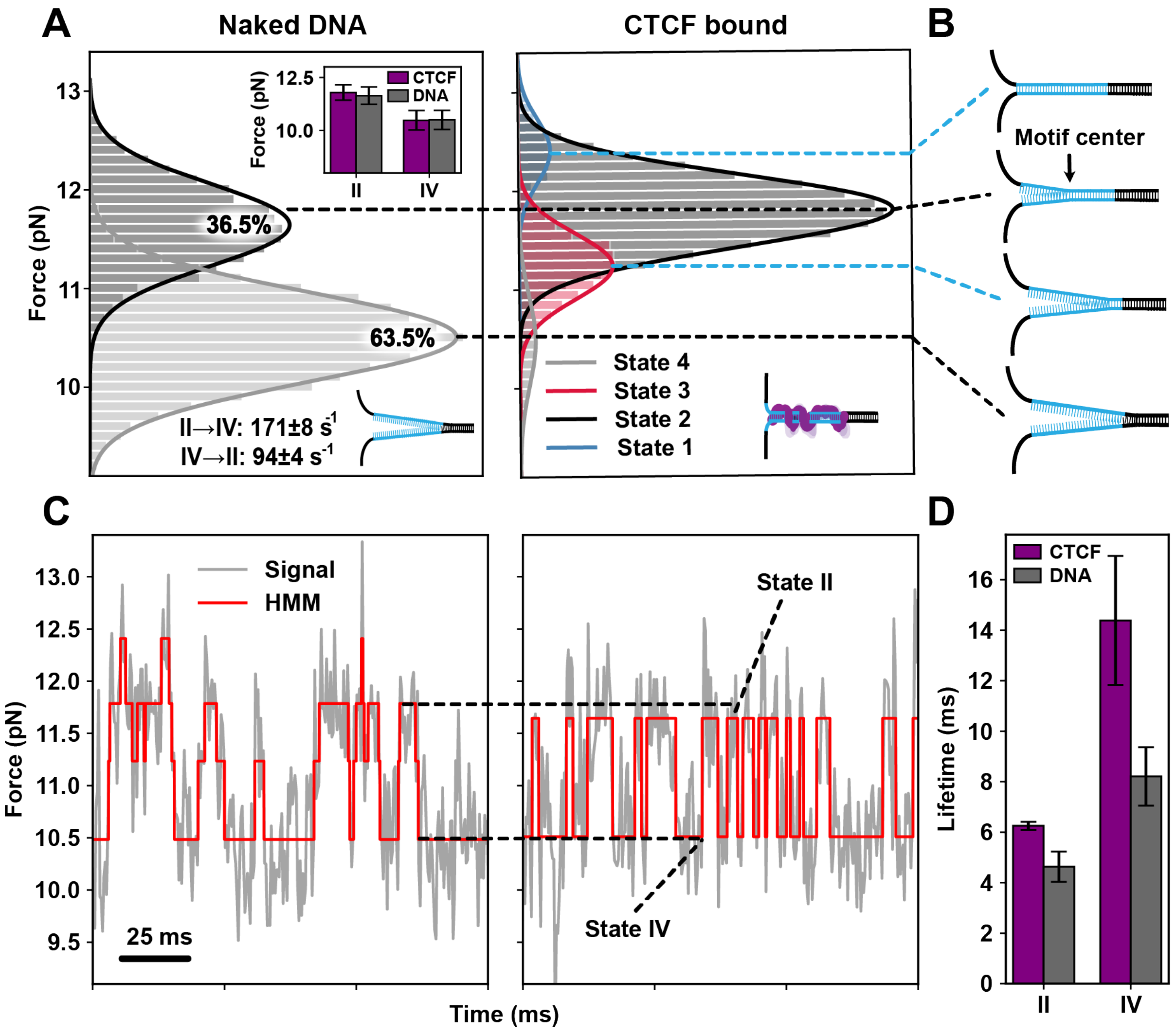


**Fig. S5. Comparison of DNA fluctuation states in naked vs. CTCF-bound DNA. (A)** Force distributions for two states of naked DNA (left) and four states of CTCF-bound DNA (right). The inset bar plot shows the mean force for two aligned states (state II and state IV) in CTCF-bound DNA with naked DNA, with error bars representing the standard deviation from Gaussian fitting. **(B)** Alignment of states with the level of dsDNA unzipping at the motif site: state IV represents a fully open motif, state III a partially closed motif, state II reflects closing at the motif center, and state I indicates a fully closed motif. **(C)** Zoomed-in view of force fluctuations in naked DNA (left) and CTCF-bound DNA (right), with HMM states highlighted in red. Dotted lines indicate the alignment of mean force for state II and IV between naked and CTCF-bound DNA. **(D)** Lifetimes of states II and IV for naked DNA vs. CTCF-bound DNA, with error bars showing the standard deviation of the fit parameter (see Materials and Methods). These plots show one representative uncorrected trace (referring to Materials and Methods, Dynamic trace preprocessing and formatting) for *PDGFRA* N-terminal CTCF WT to ensure accurate comparison between before and after binding of CTCF.

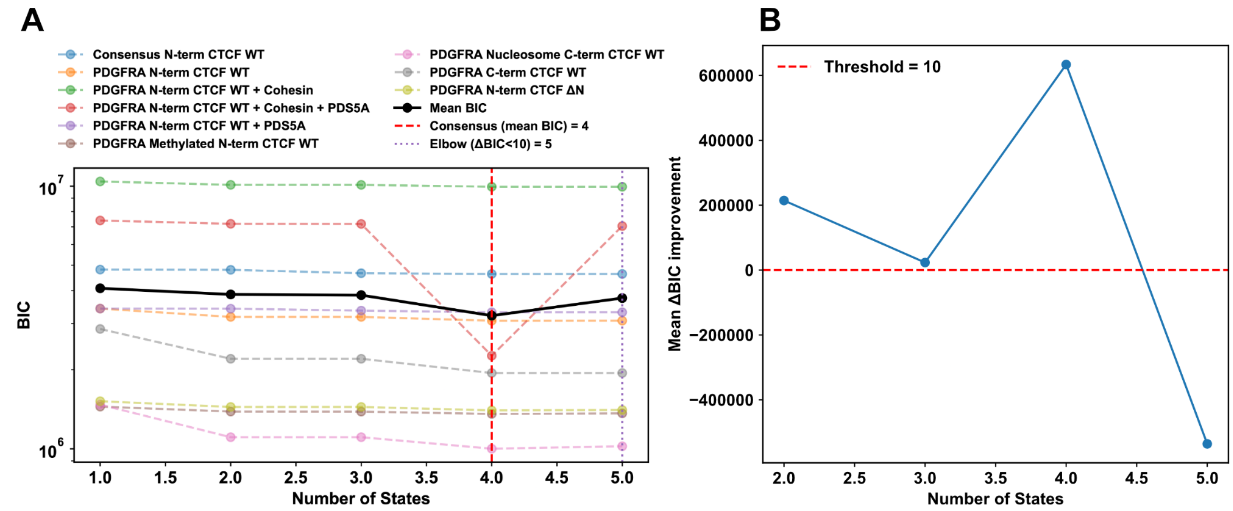


**Fig. S6. Determination of the optimal number of hidden states across conditions. (A)** Bayesian Information Criterion (BIC) values as a function of the number of states for each experimental condition (colored dashed lines). Lower values indicate a better trade-off between fit and parsimony. The black line shows the mean BIC across all conditions, which was minimized at four states (red dashed line). The dotted purple line indicates the "elbow" criterion, where the average BIC improvement fell below 10 units. **(B)** Mean BIC improvement across conditions when increasing the number of states. The largest gain was observed from three to four states, while improvements beyond four states fell below the ΔBIC = 10 threshold (red dashed line), indicating diminishing returns. Together, these analyses identify four states as the consensus model across conditions, with cohesin-bound DNA showing a marginal preference for a fifth state.

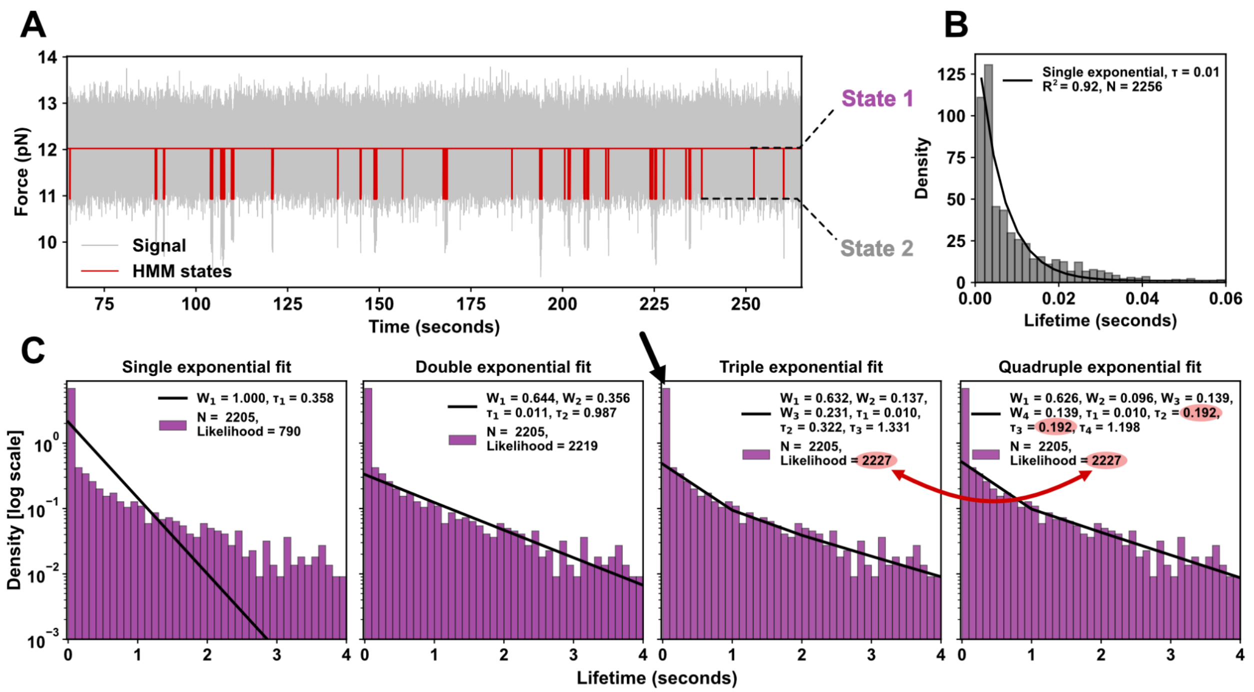


**Fig. S7. Fitting of lifetimes of CTCF-DNA binding states in a 2-state system. (A)** A representative dynamic trace of WT CTCF binding to PDGFRA is shown, with a 2-state HMM fit overlaid in red. State 2 corresponds to the ssDNA-binding conformation identified in the 4-state model described in the main text (Fig. 2). **(B)** Distribution of lifetimes for state 2 (as identified in **(A)**) fitted with a single exponential function. The fit captures the primary lifetime behavior well. **(C)** State 1 lifetime distributions fitted with single, double, triple, and quadruple exponential models using unbinned maximum likelihood estimation (MLE) (see Supplementary Methods). The triple-exponential model maximizes the likelihood, while moving to a quadruple-exponential fit shows no improvement, as two lifetimes converge to identical values, indicating redundancy. This supports the conclusion that there are three distinct binding behaviors in state 1, consistent with the 4-state model proposed in the main text. The black arrow highlights a tall first bin in the histogram, which is not adequately captured by the fit. This suggests the temporal resolution used in the experiment may miss fast transitions, hinting at the potential for additional faster states. However, increasing the temporal resolution could introduce artifacts due to instrumental noise and the Brownian motion of the beads, potentially creating artificial states rather than accurately capturing protein dynamics. We therefore conservatively consider a 4-state model.

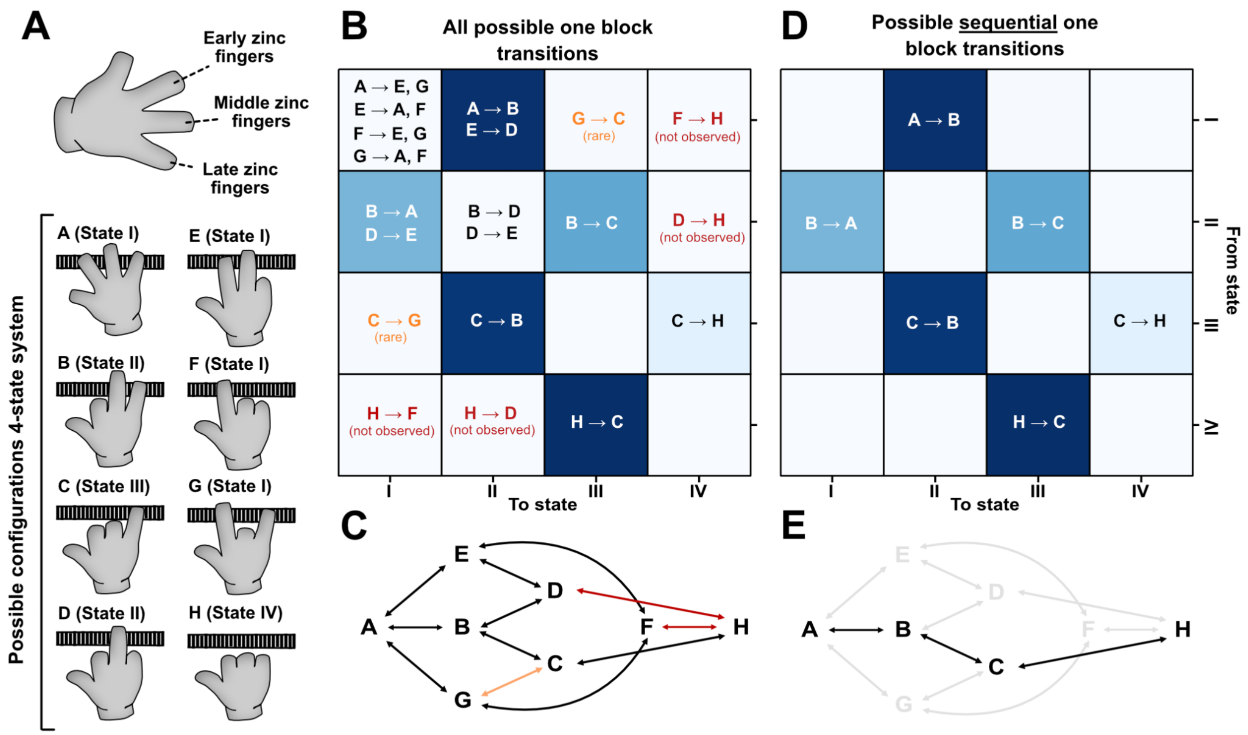


**Fig. S8. Theoretical and empirically observed conformational landscape of CTCF in a four-state system. (A)** Schematic representation of the possible configurations of CTCF assuming a four-state system based on the arrangement of its zinc fingers. Each state (A, B, C, D, E, F, G, and H) represents a unique configuration characterized by the positioning of early, middle, and late zinc fingers. **(B)** Matrix of all possible one-block (represented by one finger in **(A)**) transitions between configurations in the four-state system. Each cell represents a transition from one configuration (row) to another (column). Based on WT CTCF N-terminal *PDGFRA* dynamics measurements, blue cells indicate likely transitions, with darker shades representing more commonly observed transitions. Rarely observed transitions are marked in orange, while red text indicates transitions that have not been observed empirically. **(C)** Network diagram showing all theoretically possible transitions between states. Arrows indicate directionality, illustrating how one configuration could transition to another within the network. **(D)** Matrix of possible sequential one-block transitions. This matrix restricts transitions to sequential changes only, as hypothesized based on structural or functional constraints. Observed transitions are highlighted in blue, representing the most probable pathways under a sequential transition model. Unobserved transitions are left blank, suggesting restrictions in the protein's conformational flexibility or stability in certain configurations. **(E)** Network of empirically observed transitions, assuming sequential transition pathways only. Black arrows highlight observed transitions within the system, while greyed-out arrows indicate non-sequential transitions.

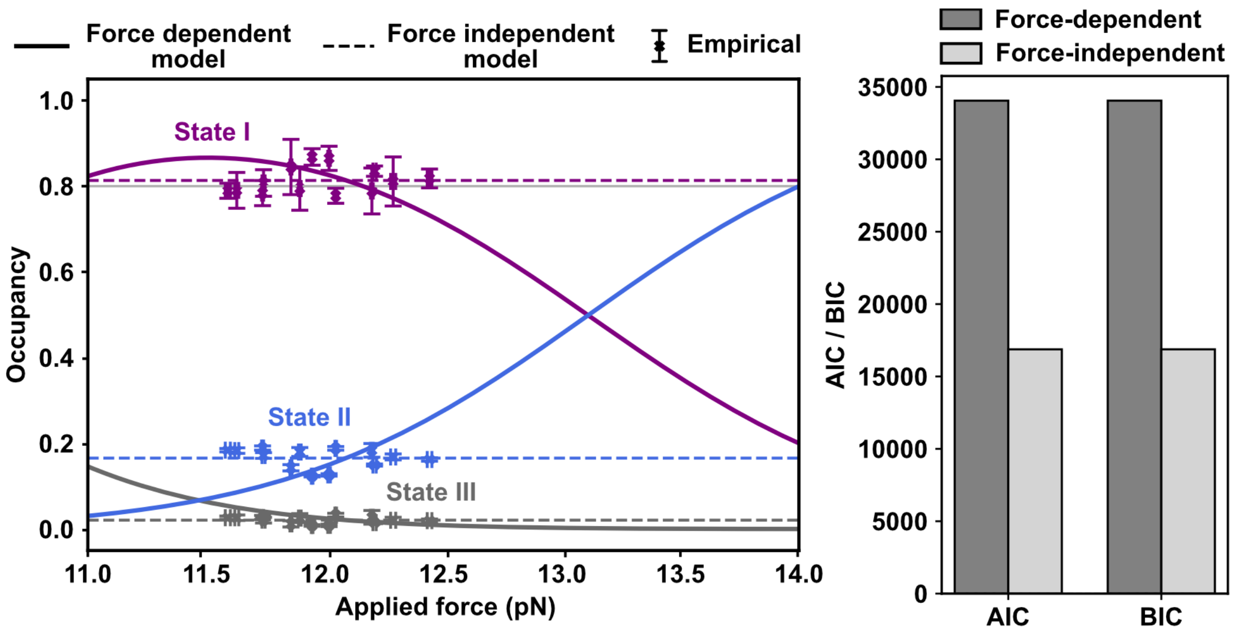


**Fig. S9. Comparison of force-dependent and force-independent models for CTCF occupancy.** Left: The theoretical and empirical occupancies of CTCF conformational states under varying force regimes. The force-dependent model (solid lines) predicts the occupancies of states I, II, and III as functions of the applied force, using Bell's model (*106*) that incorporates free energy tilting due to the applied force. The force-independent model (dashed lines) assumes constant occupancies across all force values. Empirical data points (markers) represent the observed occupancies of states I (gray), II (purple), and III (blue) at specific weighted forces, with x-axis error bars indicating the standard error of the mean of the force and y-axis error bars indicating the standard deviation between the occupancies of 10 segments of the trace. Theoretical force-independent occupancies (horizontal dashed lines) are derived from the zero-force limit of the energy landscape (see Supplementary Methods). Right: The goodness of fit of the two models was evaluated by AIC and BIC metrics, using the sum of the log-likelihoods of the three states.

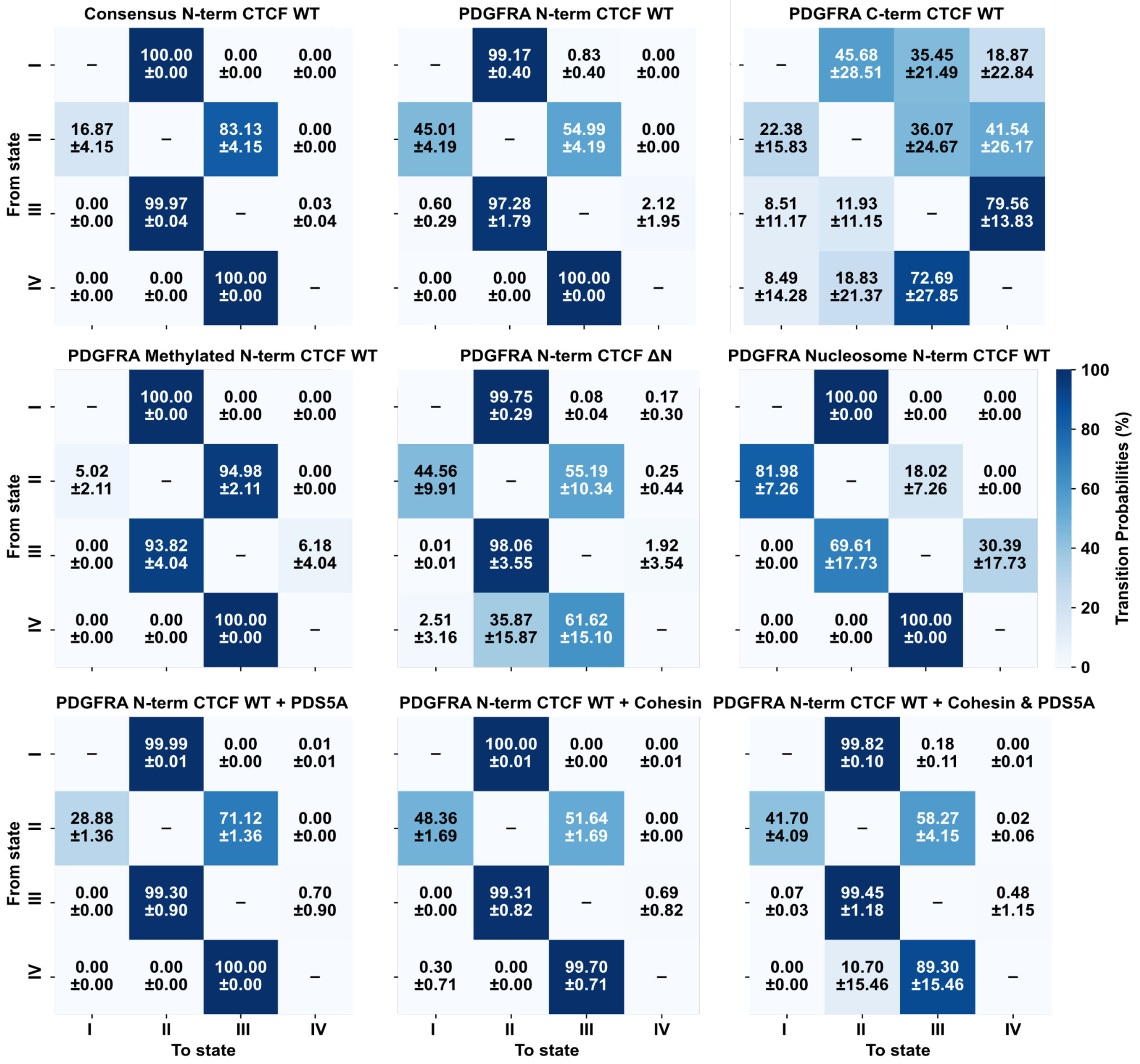


**Fig. S10. Transition probabilities excluding self-transitions** for all experimental conditions and state transitions.

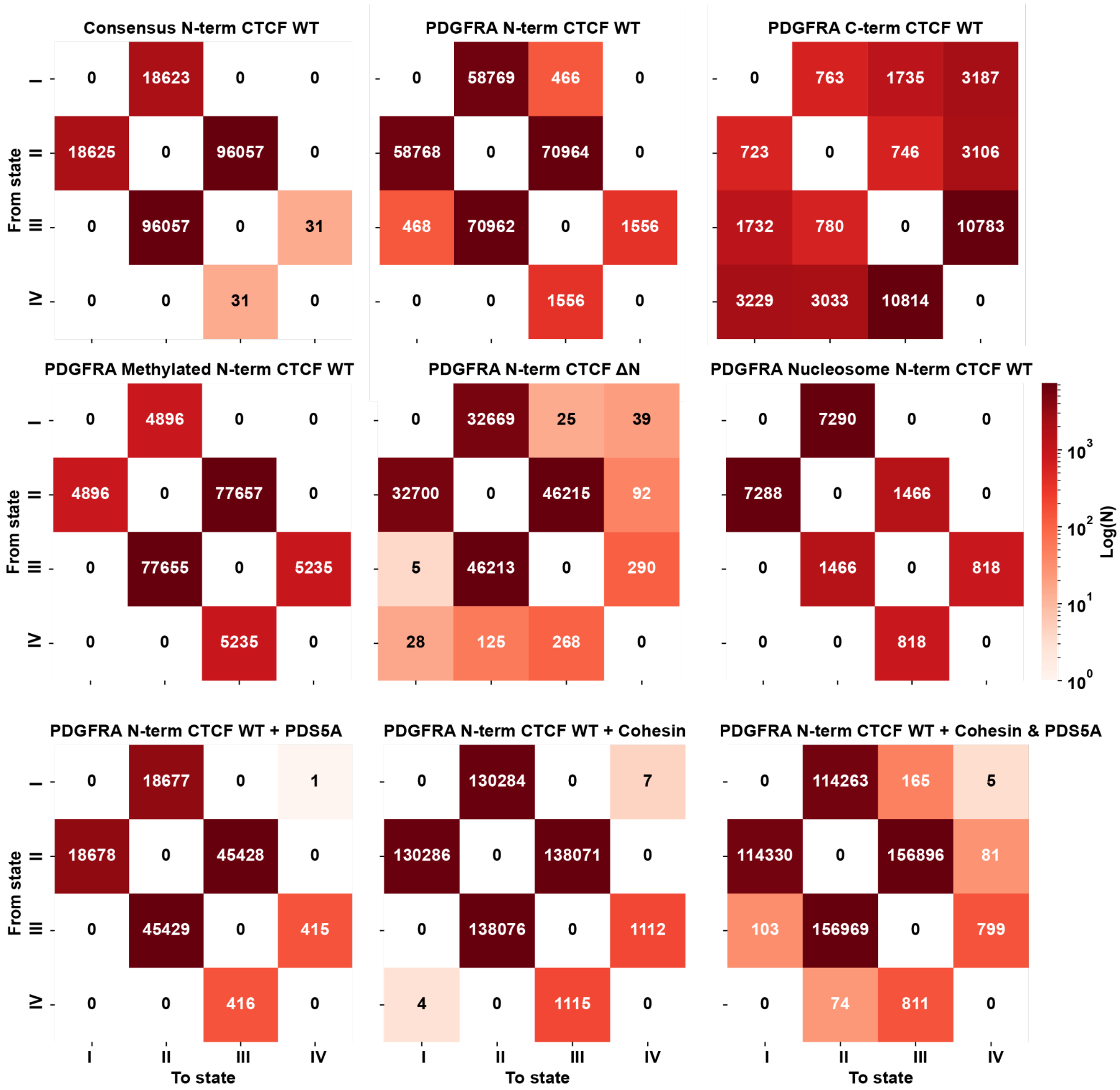


**Fig. S11. Transition counts excluding self-transitions** for all experimental conditions and state transitions.

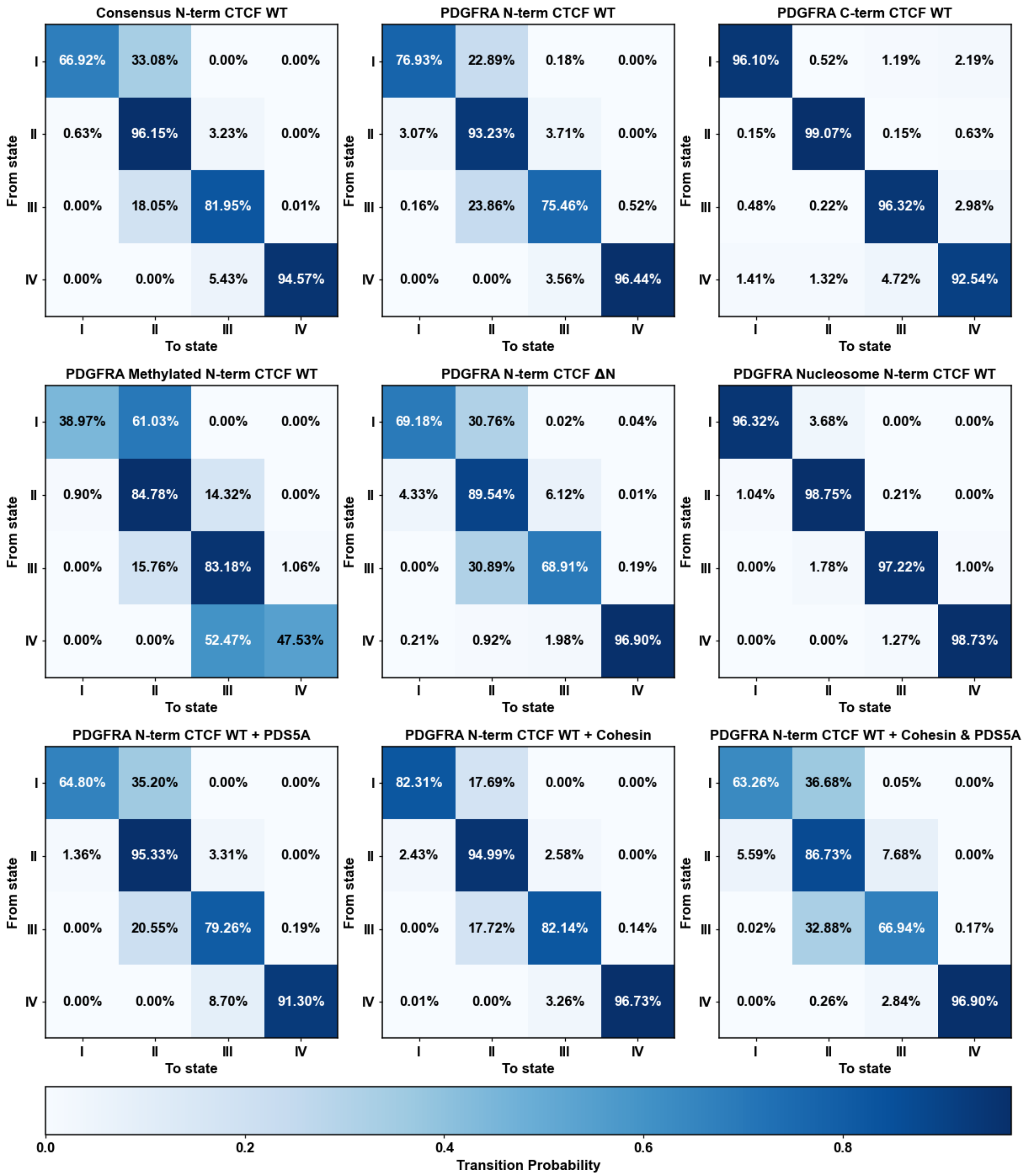


**Fig. S12. Transition probabilities including self-transitions / not transitioning** for all experimental conditions and state transitions.

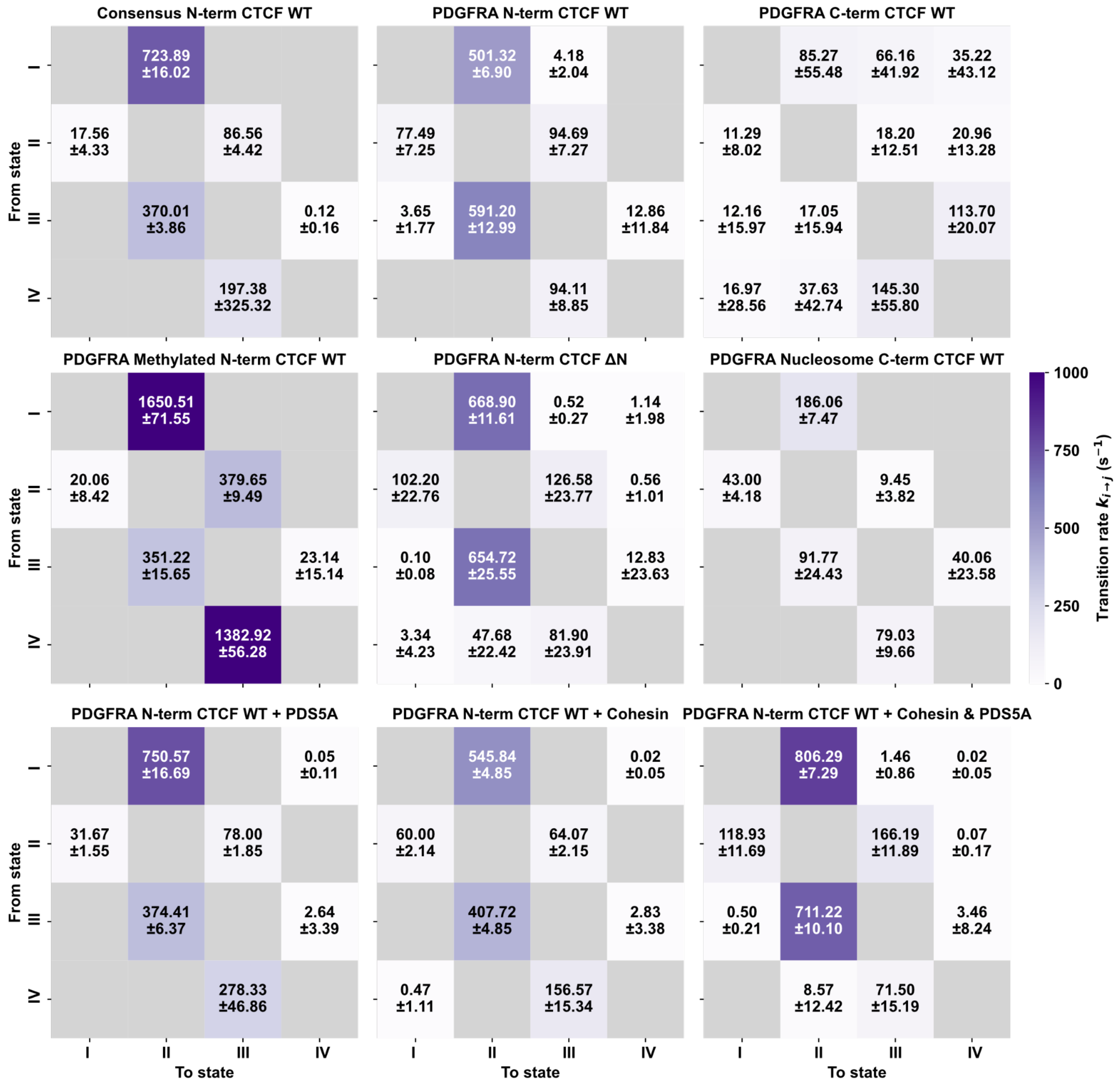


**Fig. S13. Transition rates between all states,** for all experimental conditions and state transitions. Transition rates are calculated as described in main methods.

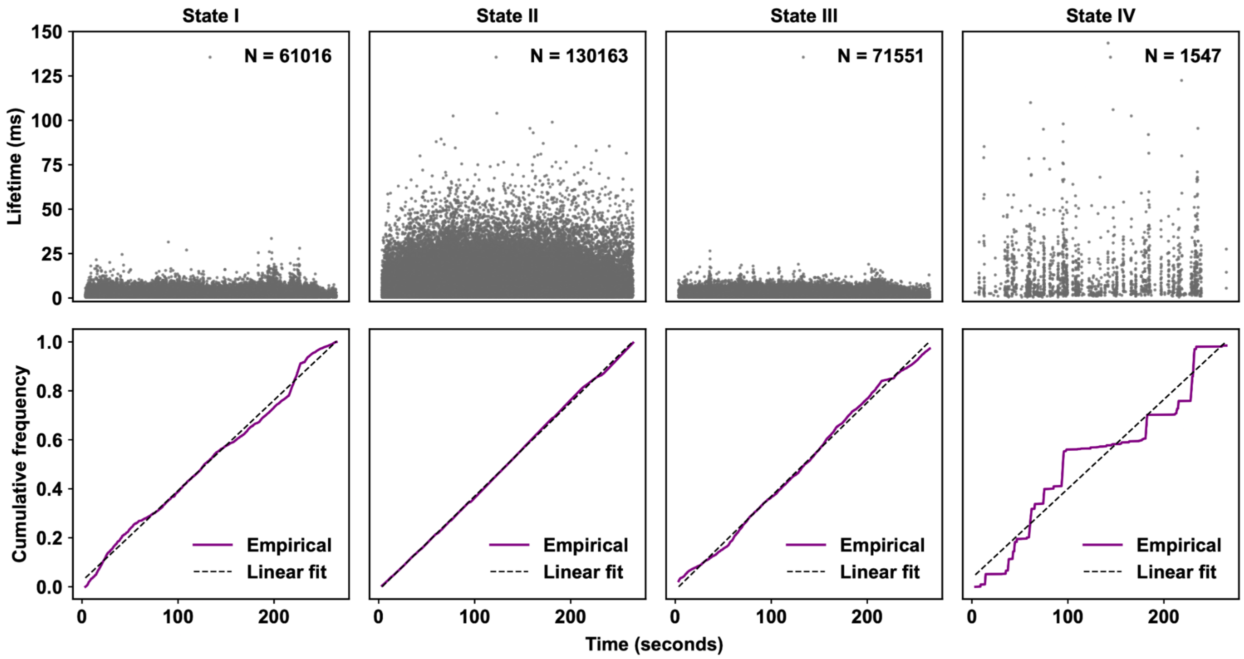


**Fig. S14. Time dependency of lifetimes and frequency of binding states of CTCF.** Lifetime of every binding state shows no correlation with binding duration of CTCF to *PDGFRA* motif (top). Similarly, the frequency of sampling each binding state shows no correlation with binding duration of CTCF to *PDGFRA* (bottom), shown by a linear correlation between frequency of state and duration since binding. These trends were seen across binding motifs and probing directions.

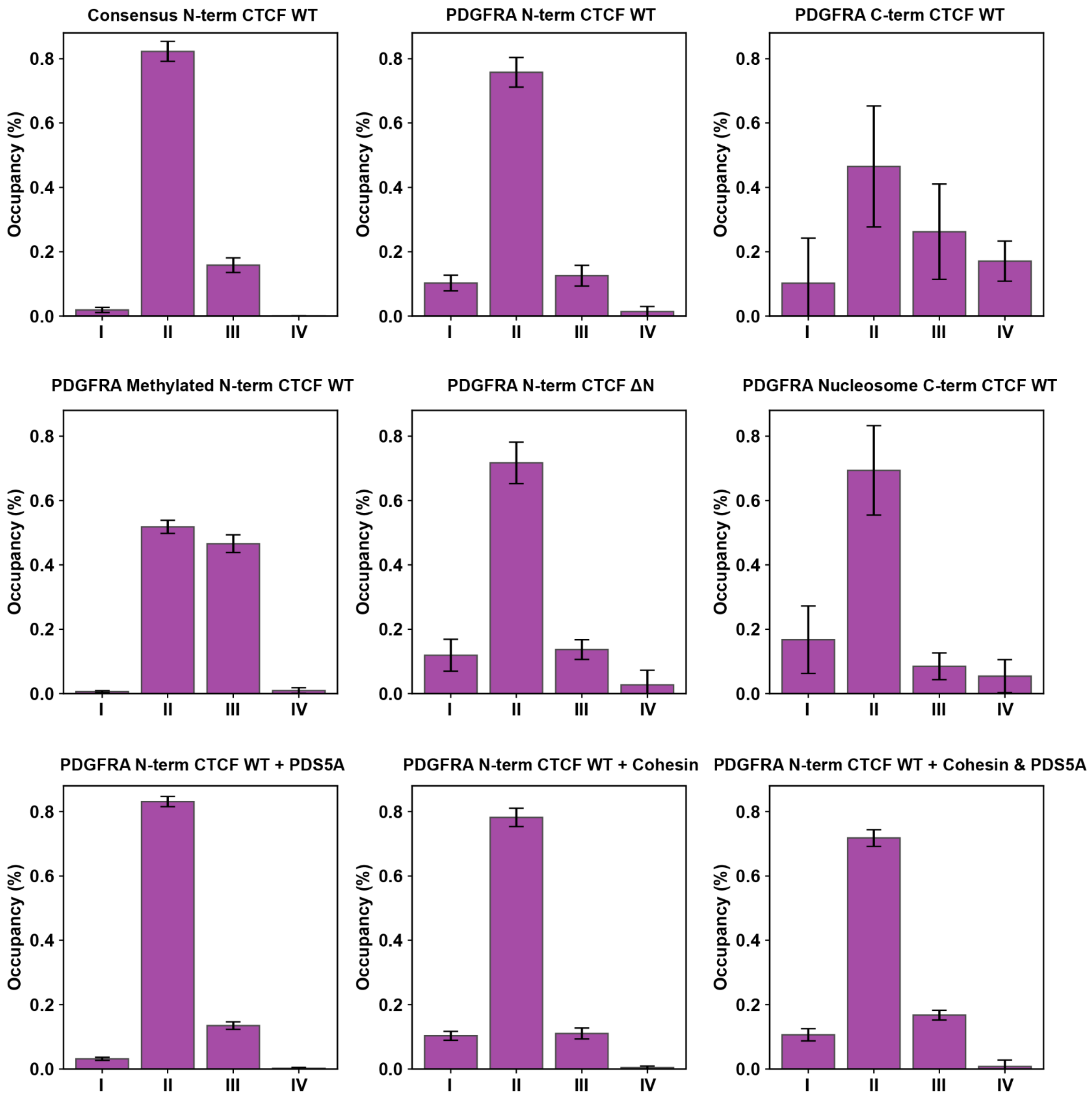


**Fig. S15. State occupancies** for all experimental conditions, determined from dynamic measurements on optical tweezers as described in the main text and Methods and Materials.

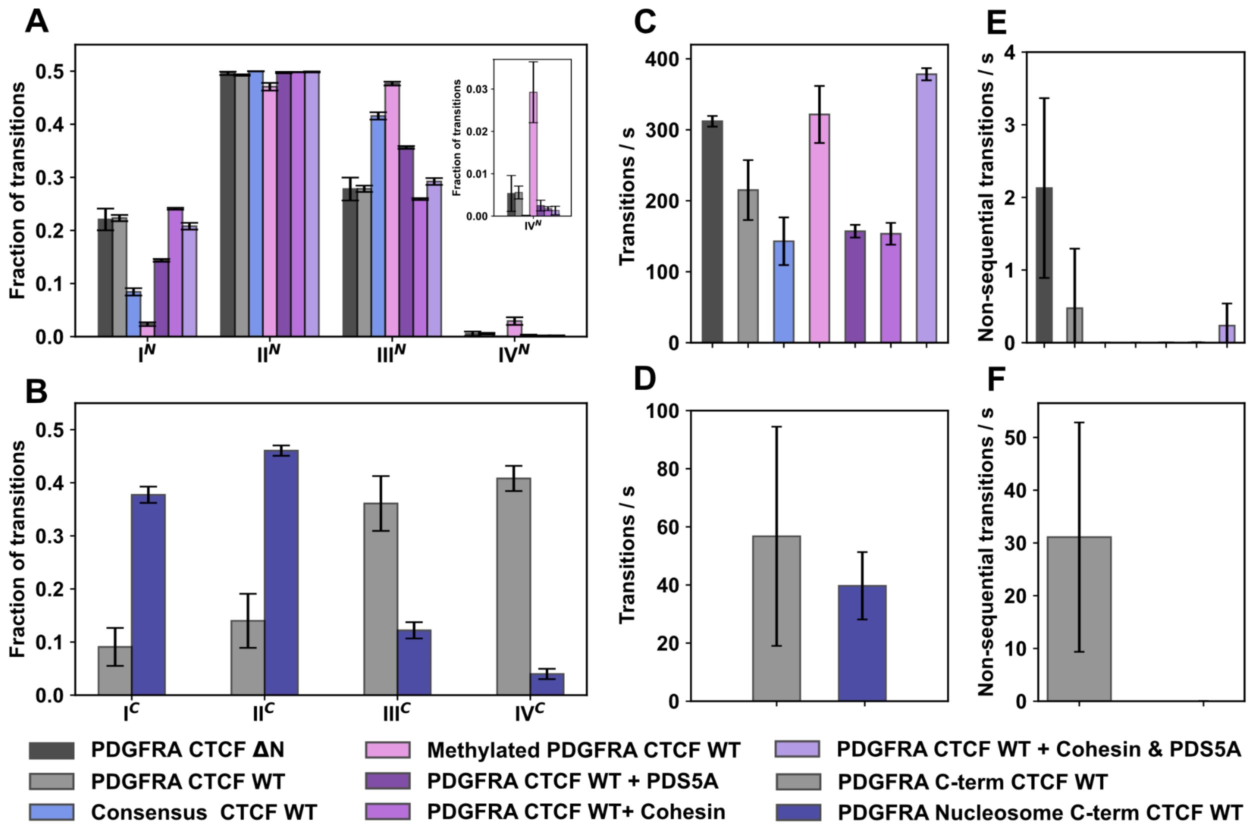


**Fig. S16. Transition statistics between states for all experimental conditions.** Number of transitions to states for **(A)** N-terminus conditions and **(B)** C-terminus conditions. Number of transitions per second for **(C)** N-terminus conditions and **(D)** C-terminus conditions. Number of non-sequential transitions (transitions to non-neighboring states) per second for **(E)** N-terminus conditions and **(F)** C-terminus conditions, errors are large due to very low statistics on these rare non-sequential transitions. Errors on bar plots represent the standard deviation between replicates, where the number of replicates are indicated in Table S1.

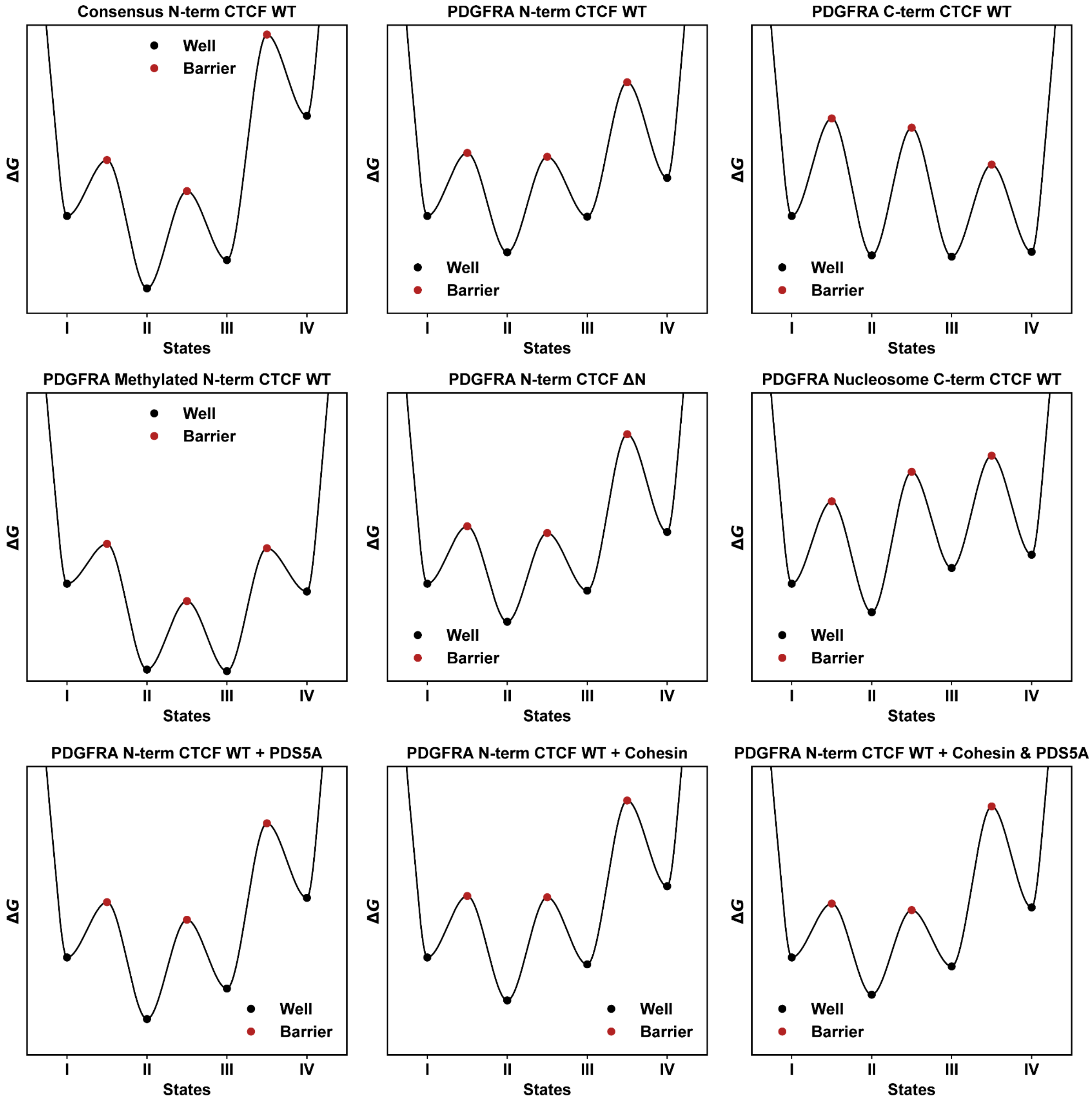


**Fig. S17. Energy landscapes** for all experimental conditions (Materials and Methods), derived from transition rates determined by dynamic optical tweezer measurements.

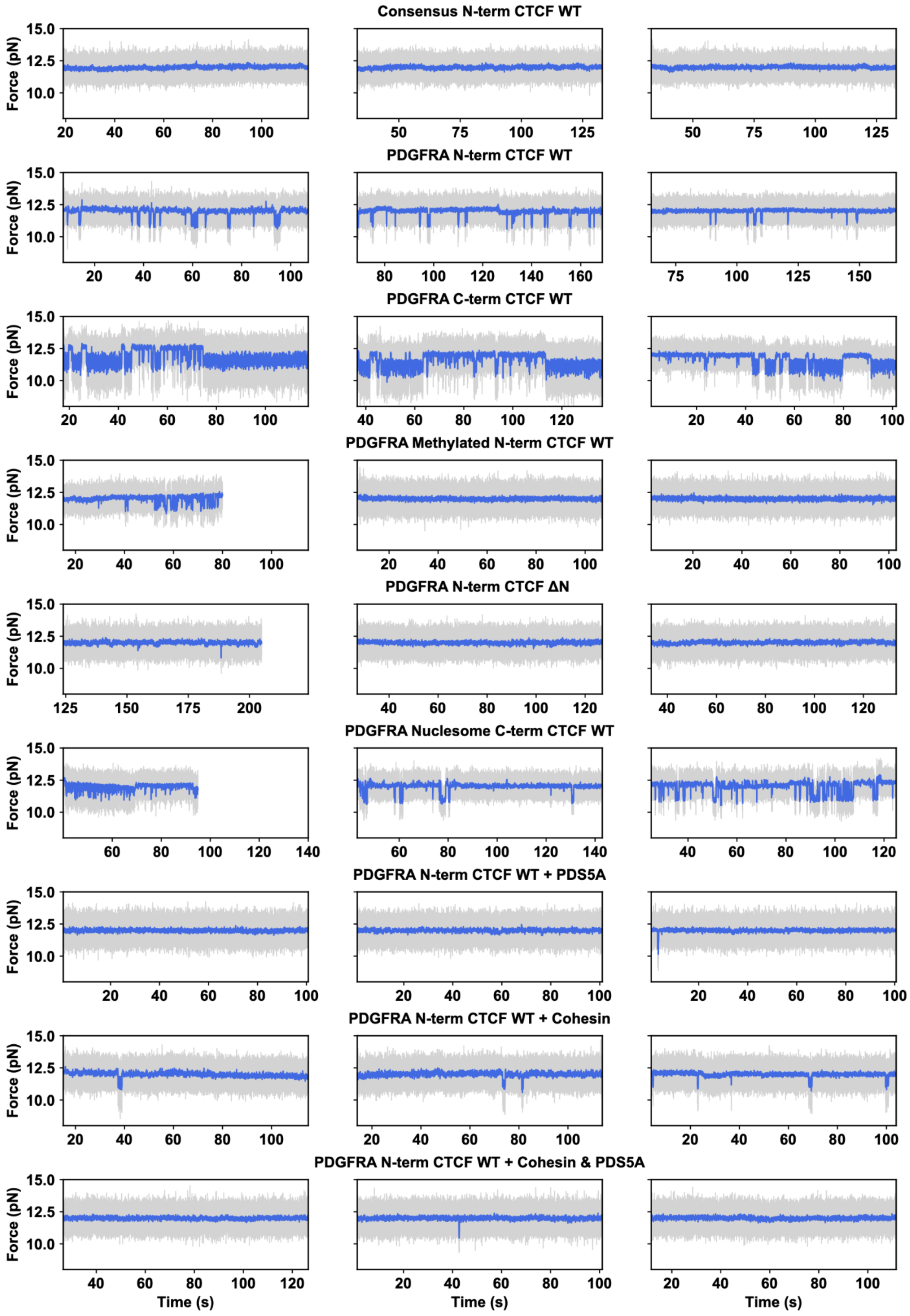


**Fig. S18. Three representative traces for all experimental conditions,** sorted by condition per row. Grey is 500 us temporal resolution and blue is downsampled 50x.

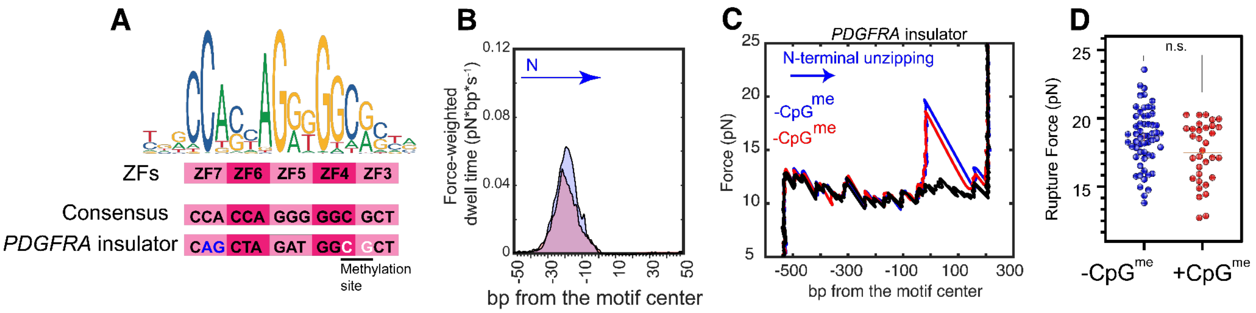


**Fig. S19. Chromatin features affect CTCF conformational dynamics. (A)** A position-weight matrix for the human CTCF binding motif and corresponding binding sites of the core ZFs (*24*, *107*). By taking the most abundant bp at each position, a consensus sequence was determined. Below it is the endogenous *PDGFRA* sequence, with deviation from consensus written in blue and the CpG methylation site written in white. **(B)** Force-weighted dwell time maps, from the N-terminal orientation, of CTCF bound to the *PDGFRA* motif in the presence (red) or absence (blue) of a regulatory CpG methylation. **(C)** Representative unzipping traces of CTCF bound to the *PDGFRA* motif in the presence (red) or absence (blue) of a regulatory CpG methylation within ZF4 on the *PDGFRA* motif from the N-terminal orientation. A representative unbound trace is shown in black. **(D)** Rupture forces, from the N-terminal orientation, of CTCF bound to the *PDGFRA* motif in the presence (red) or absence (blue) of a regulatory CpG methylation. (-methylation): n = 57, F = 18.59 ± 0.28 pN; (+methylation): n = 33, F = 17.48 ± 0.39 pN. No statistically significant difference was identified using a K-S test with a cutoff of $p < 0.05$.

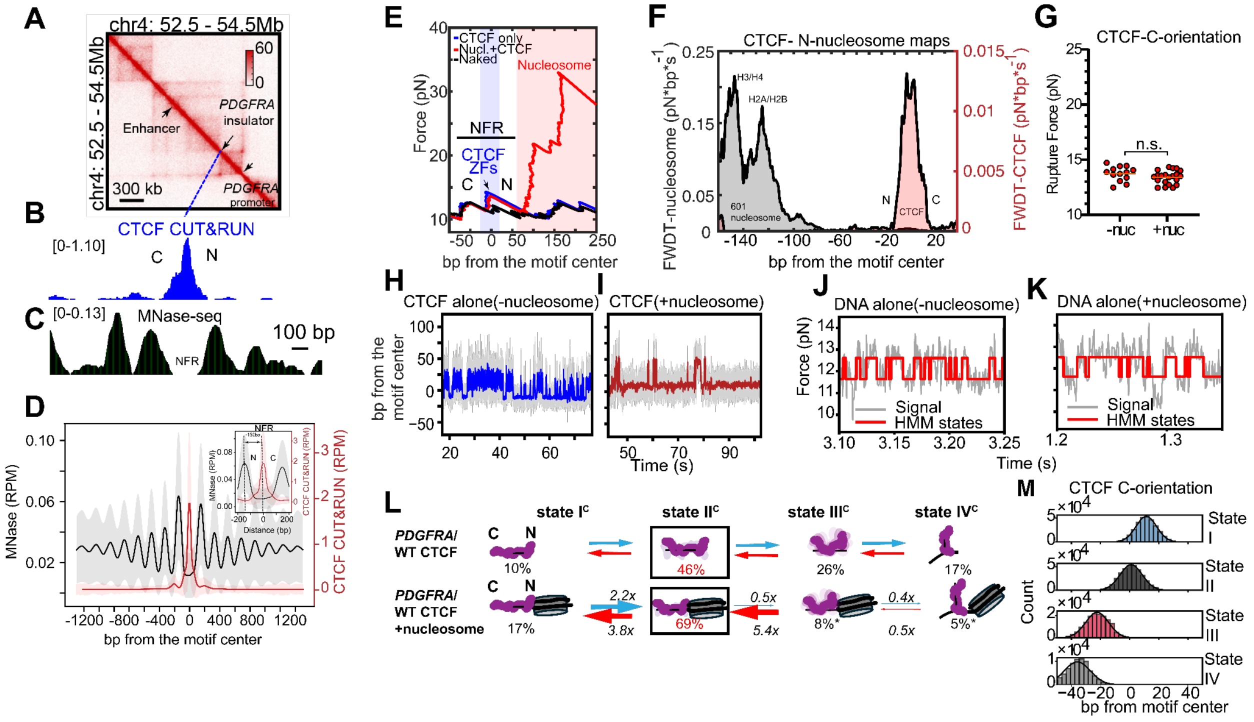

**Fig. S20: CTCF and local chromatin structure probed by GpC methyltransferase (MTase) footprinting followed by nanopore sequencing. (A)** A Hi-C map of the *PDGFRA* locus in HEK293T cells. Hi-C data were obtained from Marin-Gonzalez et al (*108*). The locations of the insulator, corresponding promoter, and corresponding enhancer are labeled and highlighted with arrows. **(B)** CTCF CUT&RUN data of the *PDGFRA* locus in HEK293T cells. **(C)** MNase-seq data of the *PDGFRA* locus in HEK293T cells. **(D)** The average MNase and CTCF CUT&RUN signal across identified, high-confidence CTCF sites. The sites represented are the same as in Fig. 1C, 3B. **(E)** A representative unzipping trace of CTCF on the *PDGFRA* motif probed from the C-terminal orientation with (red) and without (blue) an N-terminally-orientated, reconstituted, Widom 601 nucleosome. **(F)** Force-weighted dwell time map of a Widom 601 reconstituted nucleosome positioned on the N-terminal side of the *PDGFRA* motif (black) and CTCF bound to the *PDGFRA* motif, both unzipped from the C-terminal orientation. **(G)** Rupture forces from the C-terminal orientation of CTCF bound to *PDGFRA* in the presence or absence of an N-terminal nucleosome. (-nucleosome): $n = 13$, $F = 13.73 \pm 0.19$ pN; (+nucleosome): $n = 21$, $F = 13.4 \pm 0.13$ pN. No statistically significant difference was identified using a K-S test with a cutoff of $p < 0.05$. **(H)** Representative CTCF-*PDGFRA* fluctuations probed from the C-terminal orientation with (right, red) or **(I)** without (left, blue) an N-terminal nucleosome. **(J-K)** Zoomed-in view of force fluctuations from the C-terminal orientation in **(J)** naked *PDGFRA* DNA and **(K)** *PDGFRA* DNA with an N-terminal nucleosome, with HMM states highlighted in red. **(L)** Transition rates of CTCF state dynamics on the *PDGFRA* motif probed from the C-terminal orientation with (bottom) or without (top) an N-terminally-orientated nucleosome. Black boxes indicate the highest occupancy state, with a percentage written in red. Other occupancy percentages are written in black. All transition rates and occupancies can be found in Fig. S15, Fig. S16, and Table S2. P-values are calculated with a two-tailed Mann-Whitney test relative to C-terminal WT CTCF bound to *PDGFRA* and can be found in Table S2, with significance indicated as $^*p<0.05$. **(M)** Forces, partitioned into states by the HMM model, collected at all time points from 4 representative replicates of C-terminal CTCF-*PDGFRA* dynamics were converted to bp from the CTCF motif center. Gaussian fits (black lines) provide mean ± standard deviation for each state: 11.43 ± 7.13 (n = 318,378), 1.05 ± 7.65 (n = 345,360), -22.35 ± 7.86 (n = 150,091) and -35.97 ± 8.97 (n = 73,096) for states I-IV, respectively.

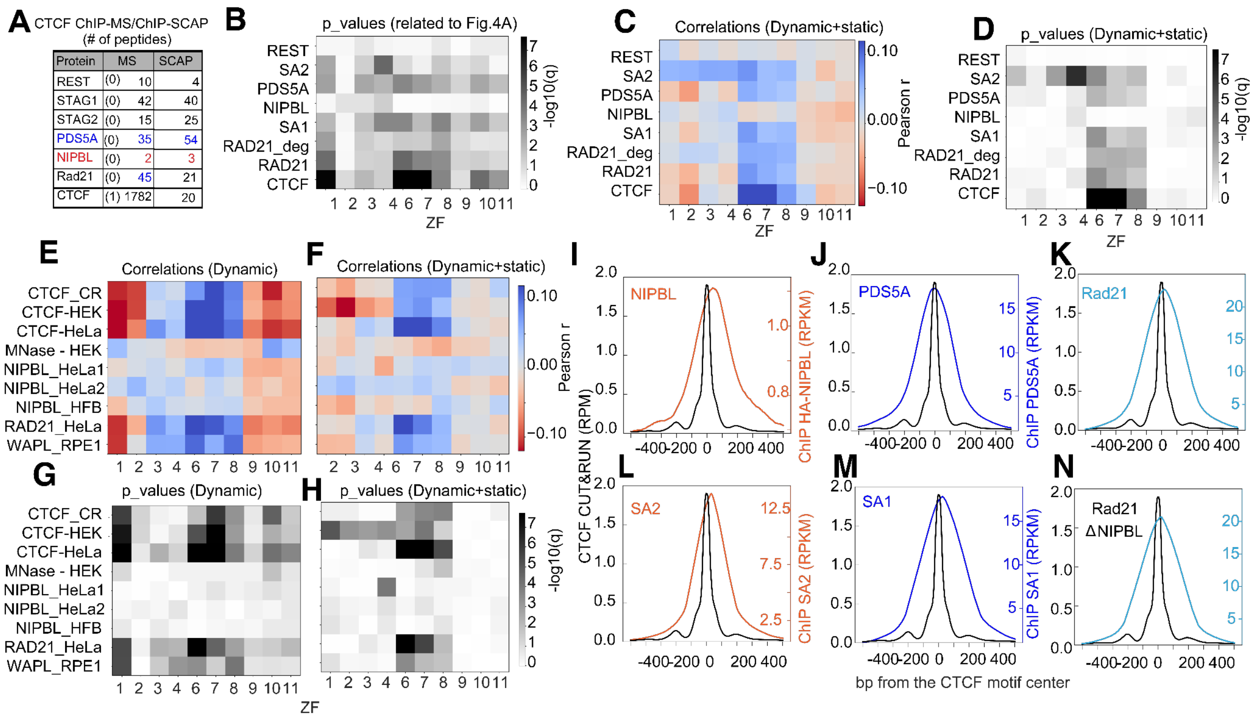


**Fig. S21. CTCF ZF-resolved correlations with cohesin subunits and chromatin features. (A)** Table of CTCF ChIP-MS and ChIP-SCAP values of cohesin subunits in mouse embryonic stem cells obtained from Ortabozkoyun et al (*58*) and from Dehingia (*59*) et al. **(B)** Benjamini-Hochberg (BH)-adjusted p-values for main Fig. 3B. **(C)** Pearson correlation between publicly available ChIP-seq datasets (*57*, *109*) for cohesin subunits and GpC inaccessibility, calculated for the subset of inaccessible and partially accessible molecules (as in Fig. 3B) at each ZF. **(D)** BH-adjusted p-values for **(C)**. **(E-H)** Pearson correlations and corresponding p-values for additional factors/assays/cell types/antibodies, including CUT&RUN for CTCF in HEK293T (original data), ChIP-seq for CTCF in HEK293 (ENCODE), CTCF-GFP in HeLa cells (Banigan et al) (*109*), RAD21 in HeLa (Banigan et al) (*109*), WAPL in RPE-1 (Popay et al) (*57*), MNase-seq in HEK293T (original data), NIPBL-Bethyl-antibody, NIPBL-HA-antibody in HeLa (Banigan et al (*109*), NIPBL in healthy primary human fibroblasts (HFB) (Garcia et al) (*110*). **(I-N)** ChIP-seq metaplots centered on sampled CTCF motifs showing enrichment of **(I)** NIPBL, **(J)** PDS5A, **(K)** RAD21, **(L)** STAG2, **(M)** STAG1, and **(N)** RAD21 following 24 h NIPBL degron induction (*57*). A CTCF CUT&RUN metaplot is overlaid in each panel as a reference. Accession numbers can be found in Materials and Methods.

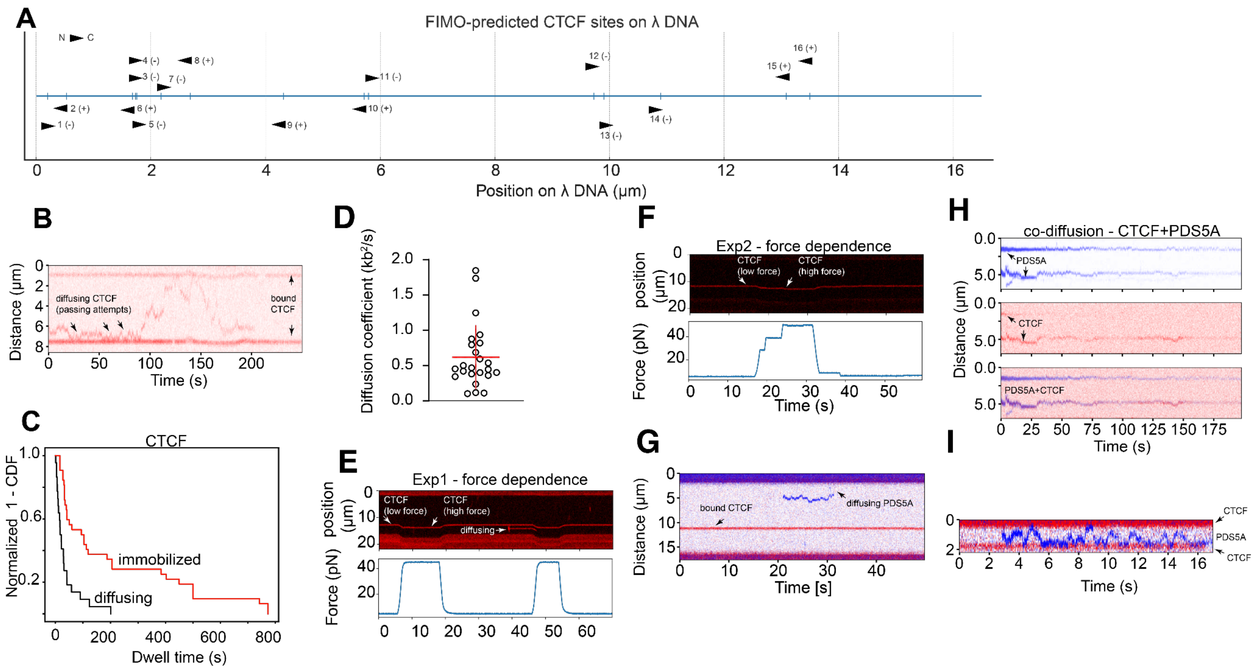


**Fig. S22. Confocal imaging of CTCF and PDS5A on optical tweezers. (A)** A sequence map of the λ DNA used for the following experiments. The location, strand (+ or -), and orientation of CTCF sites were predicted with FIMO and depicted as arrows pointing in the direction of the C-terminus. **(B)** A representative kymograph of 3 CTCF-JF646 molecules (red) bound to λ DNA under 5 pN tension. The top and bottom molecules exhibit minimal positional fluctuation and are considered motif-bound, while the middle molecule diffuses between them. **(C)** 1-CDF of CTCF dwell times when diffusive (black; median = 18.5s, n = 22) and immobilized (red; median = 98.5 s, n = 32). **(D)** Diffusion coefficients of each sliding CTCF-JF646 molecule (mean = 0.62 ± 0.45 $kb^2/s$, n = 24). **(E-F)** Representative kymographs of CTCF-JF646 molecules (red, above) under varying force (blue, below). Immobilized CTCF molecules remained bound through increases and decreases in force ranging from 5 to 45 pN, while a diffusive CTCF molecule in **(E)** dissociated when force was increased. **(G-I)** Representative kymographs of CTCF-JF646 (red) and PDS5A-GFP (blue) bound to λ DNA under 5 pN tension. In **(G)**, there is one observed immobile CTCF-JF646 molecule and one diffusive PDS5A-GFP molecule. In **(H)**, the 3 panels show, from top to bottom, the blue imaging channel (PDS5A-GFP visible), the red imaging channel (CTCF-JF646 visible), and an overlay of the two channels. In the panels, 2 sets of colocalized CTCF-JF646 and PDS5A-GFP molecules are seen co-diffusing. In **(I)**, a diffusive PDS5A-GFP molecule is seen constrained between two immobilized CTCF-JF646 molecules.

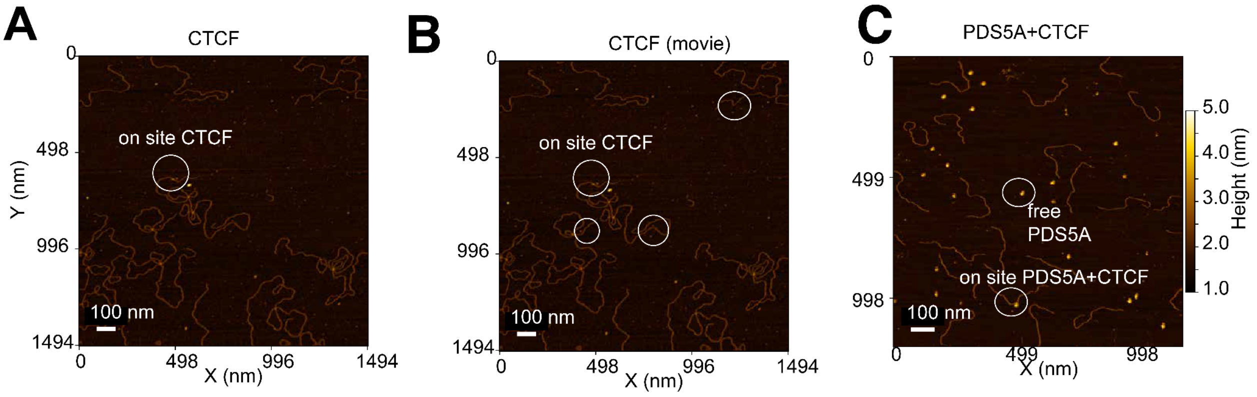

**Fig. S23. Structural examination of CTCF and PDS5A using AFM. (A-B)** Representative AFM images of CTCF and the 8 kb DNA substrate used in Fig. 3F. In **(A)**, density, located at the known CTCF binding motif, that resembles CTCF and potential IDRs is circled and labeled as "on site CTCF". **(B)** Is another frame of a movie from the same field of view as **(A)**, but with additional circles around other DNA-colocalized density that moves during the movie, interpreted as dynamic, DNA-bound CTCF molecules. **(C)** A representative AFM image of CTCF and PDS5A in the presence of ~1 kb DNA (1,006 bp) segment of the 8 kb DNA substrate with a central CTCF consensus motif (510 bp and 495 bp from motif center on each side). A complex that is localized to the known binding site is circled and labeled. Background density that has been identified as unbound PDS5A is also circled and labeled.

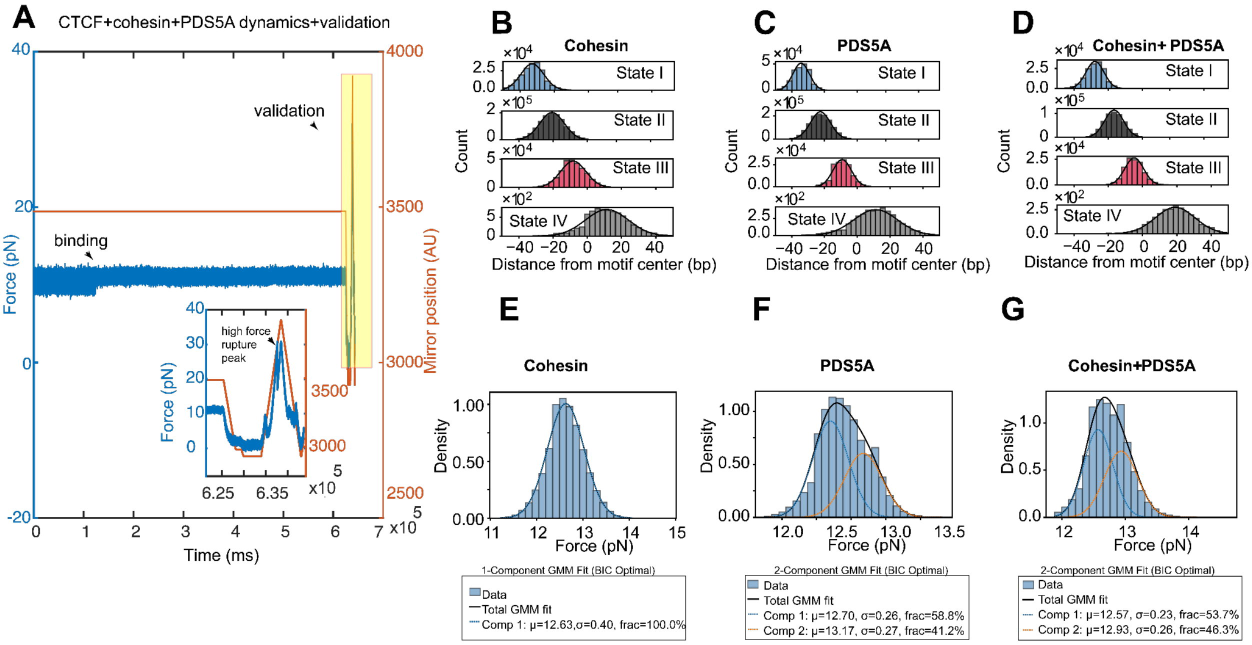


**Fig. S24: Dynamic measurements of cohesin and cohesin subunits. (A)** A representative dynamic measurement of CTCF from the N-terminal orientation bound to *PDGFRA* in the presence of PDS5A and cohesin. At the end of the measurement, an unzipping validation was performed (see Fig. S3), in which the resulting rupture force belonged to the highly stable population (see Fig. 4F). **(B–D)** Forces, partitioned into states by HMM models, were converted to base pairs from the CTCF motif center for N-terminal dynamic measurements of CTCF in the presence of **(B)** cohesin (6 representative traces), **(C)** PDS5A (4 representative traces), and **(D)** cohesin together with PDS5A (3 representative traces). Gaussian fits (black lines) provide the mean ± standard deviation and number of transitions for states I–IV, respectively: **(B)** −32.8 ± 6.7 bp (n = 188,728), −20.6 ± 7.1 bp (n = 1,201,162), −8.2 ± 7.4 bp (n = 288,988), 11.4 ± 12.5 bp (n = 7,313); **(C)** −34.3 ± 5.0 bp (n = 214,881), −22.5 ± 6.0 bp (n = 1,182,874), −9.2 ± 5.6 bp (n = 145,828), 11.2 ± 13.3 bp (n = 4,213); **(D)** −27.8 ± 5.2 bp (n = 148,350), −16.6 ± 5.5 bp (n = 526,857), −4.9 ± 5.5 bp (n = 120,276), 19.0 ± 11.5 bp (n = 2,645). **(E-G)** Histograms of state I forces across all time points of CTCF in the presence of **(E)** cohesin, **(F)** PDS5A, and **(G)** cohesin and PDS5A. As state I in some cohesin subunit conditions were the only distributions across states and conditions that did not fit single Gaussians well, we fit the histograms with a Gaussian mixture model to identify whether they contained 1 or 2 populations and the relative weight of each population. CTCF and cohesin contained 1 population (mean = 12.6 ± 0.4 pN), CTCF and PDS5A contained a lower force (58.8% weight, mean = 12.7 ± 0.3 pN, n = 31,210) and higher force population (41.2% weight, mean = 13.2 ± 0.3 pN, n = 21,856), and CTCF, PDS5A, and cohesin contained a lower force (53.7% weight, mean = 12.6 ± 0.2 pN, n = 167,349) and higher force population (46.3% weight, mean = 12.9 ± 0.3 pN, n = 144,136).

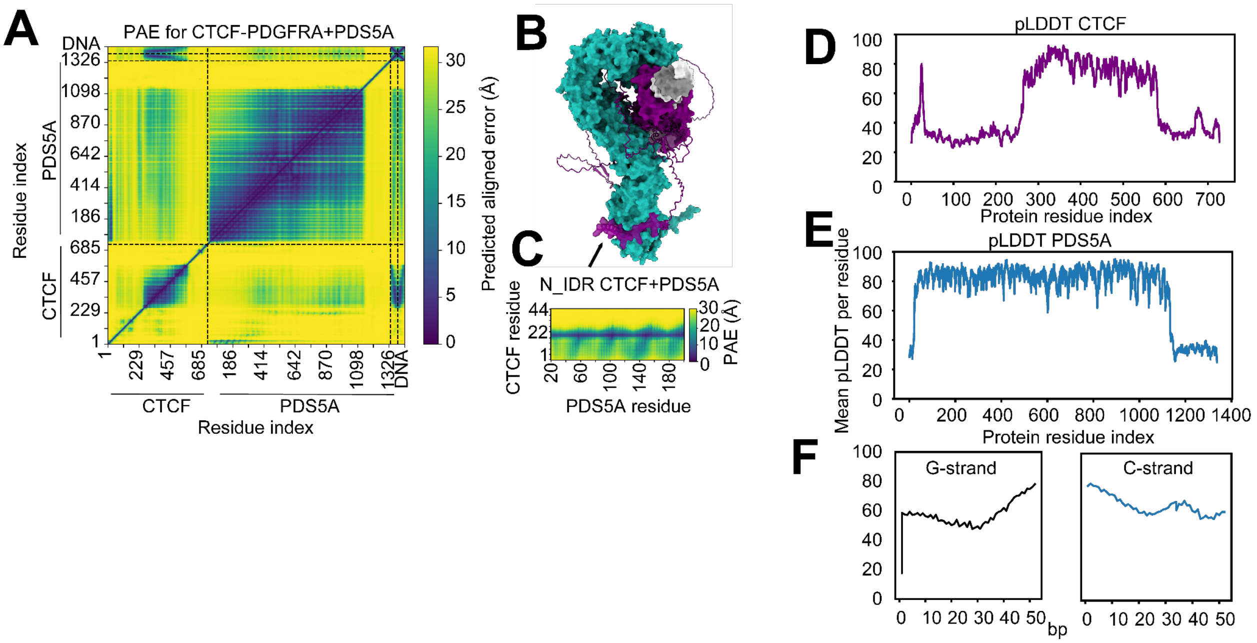


**Fig. S25. AlphaFold 3-predicted structure of CTCF-PDS5A-*PDGFRA*** **(A)** The CTCF-PDS5A-*PDGFRA* AlphaFold 3-predicted structure shown in Fig. 4A highlighting that the model recapitulated a previously identified (*38*, *39*) PDS5A-CTCF N-terminal IDR interaction. **(B)** A zoomed-in view of the PAE plot shown in **(C)**, focusing on the PDS5A-CTCF N-terminal IDR interaction shown in **(A)**. Residues 1-50 of CTCF and residues 20-200 of PDS5A are shown. **(D-F)** Plots of the per residue mean pLDDT value for each chain of the AlphaFold 3 model of *PDGFRA*-bound CTCF and PDS5A. **(D)** shows the pLDDT values for CTCF, **(E)** for PDS5A, and **(F)** for the *PDGFRA* G-strand (left) and C-strand (right).

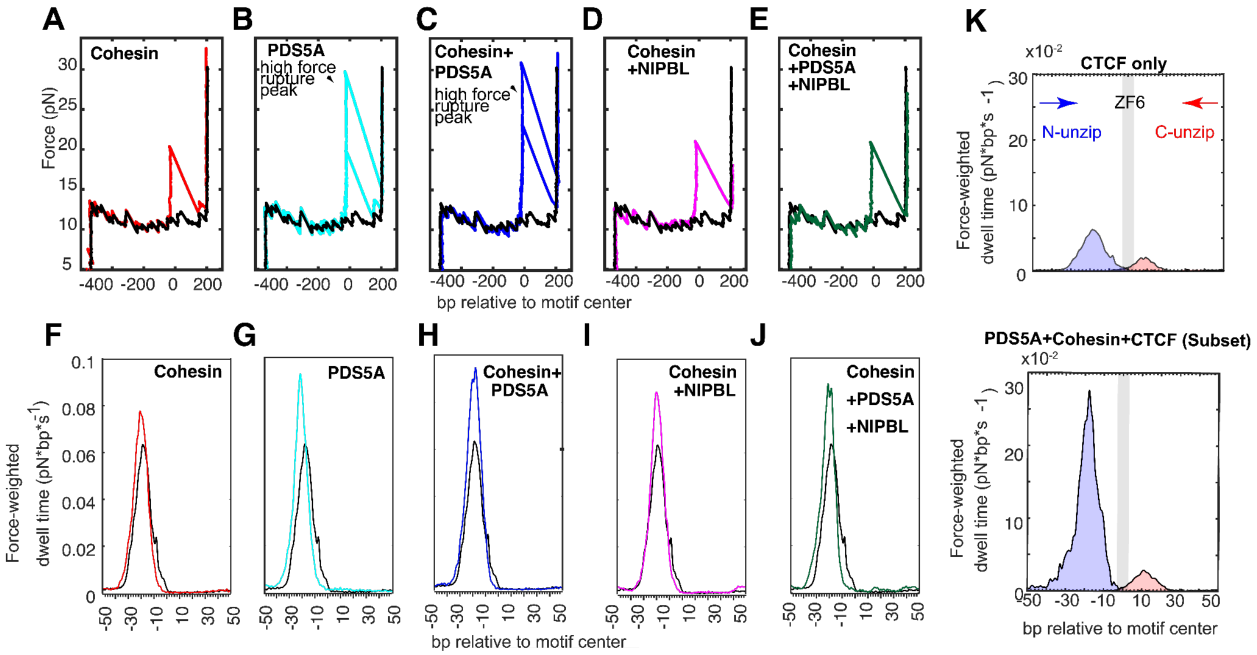


**Fig. S26. Optical tweezers measurements of cohesin and subunits. (A-E)** Representative unzipping traces from the N-terminal orientation of CTCF in the presence of **(A)** cohesin, **(B)** PDS5A, **(C)** cohesin and PDS5A, **(D)** cohesin and NIPBL, and **(E)** cohesin, PDS5A, and NIPBL. For conditions identified to have a highly-stable population of rupture forces (PDS5A with and without cohesin), a representative trace from both populations is shown. In all panels an unbound unzipping trace is shown in black. Violin plots of all rupture forces for each condition can be found in Fig. 4F. **(F-J)** Force-weighted dwell time maps from the N-terminal orientation of CTCF-only (red) and CTCF in the presence of **(F)** cohesin, **(G)** PDS5A, **(H)** cohesin and PDS5A, **(I)** cohesin and NIPBL, and **(J)** cohesin, PDS5A, and NIPBL (blue). **(K)** Force-weighted dwell time maps of CTCF bound to the *PDGFRA* motif (top) and the CTCF-PDS5A-cohesin complex bound to the *PDGFRA* motif (bottom). For the N-terminal orientation of the CTCF-PDS5A-cohesin complex, the higher rupture force population denoted in Fig. 4D, 4F is depicted. The location of ZF6 is overlaid as gray bars.

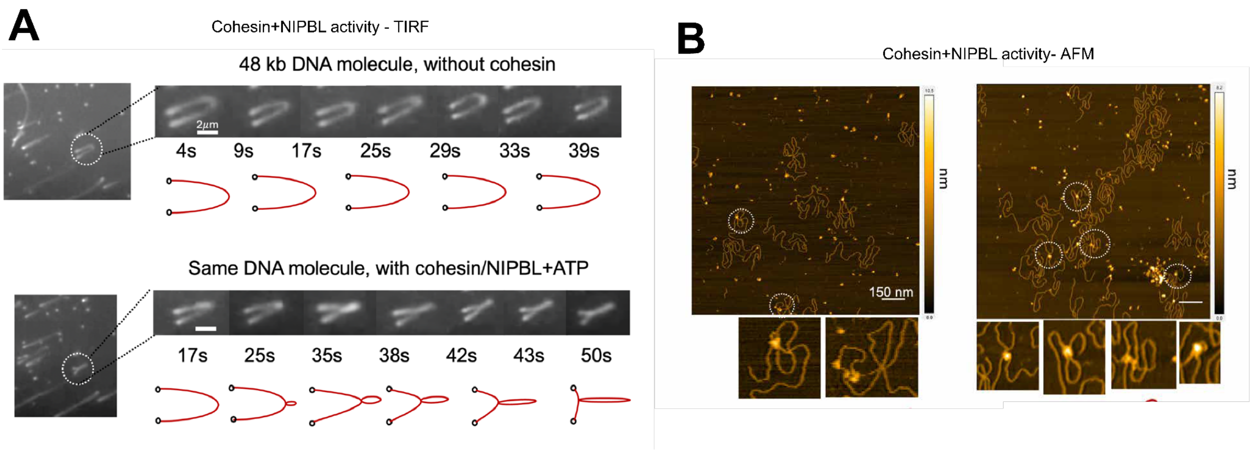


**Fig. S27. Activity of recombinant cohesin complexes (A)** Representative images of a single DNA molecule over time in a single-molecule TIRF loop extrusion assay (*11*, *19*) used to verify the activity of our cohesin proteins. Using flow to stretch and image immobilized DNA over time, we observed a consistent arc shape in the absence of cohesin (top) and an actively extruding loop in real-time in the presence of cohesin, NIPBL, and ATP. An observed loop extrusion rate of 1.6 kb/s matched previously reported values (*11*, *19*) **(E)** Representative AFM images of 8 kb DNA after incubation with CTCF, cohesin, NIPBL, and ATP at 37C. Structures resembling loops are circled and shown zoomed in below.

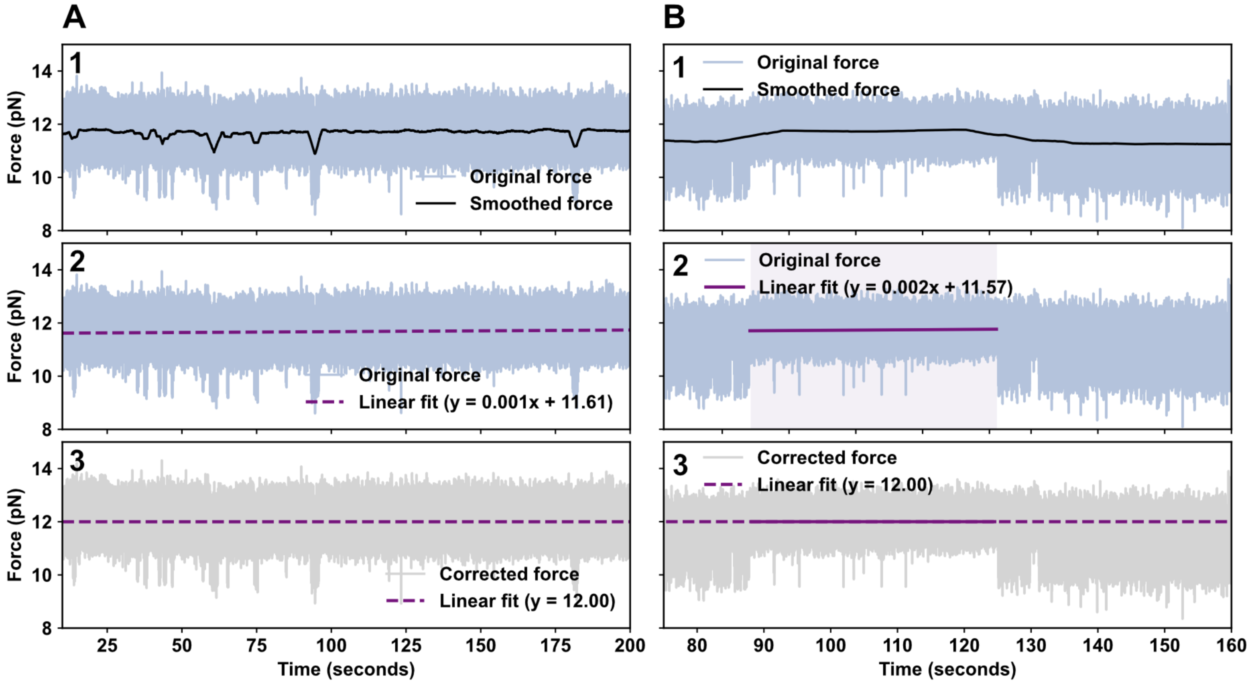


**Fig. S28. Correction of force over time traces of CTCF-DNA complexes. (A)** Homogeneous traces with little fluctuations to the open state were corrected by smoothing of the trace using a rolling window with size 5000. After smoothing it was fitted with a linear function and the trace was corrected to have a slope of zero and an intercept at 12.00 pN. **(B)** Heterogeneous traces with long periods in the open state were corrected in the same way but only to the parts in the closed states. This was done in order to ensure the same intercept for the high force (closed DNA) states.

**Table S1. Summary of duration and number of states for all experimental conditions.** The number of experiments and total duration of measurements across experimental conditions as well as how many HMM states were found to be ideal and the observed number of transitions between states.

| Sequence and additional proteins | CTCF | Direction | N.o. independent experiments | Total duration | N.o. HMM states | Total n.o. transitions |
|---|---|---|---|---|---|---|
| *PDGFRA* | WT | N | 10 | 21 min | 4 | 263,510 |
| *PDGFRA* | WT | C | 5 | 10 min | 4 | 145,367 |
| Consensus | WT | N | 9 | 30 min | 4 | 229,426 |
| *PDGFRA* | ΔN | N | 5 | 9 min | 4 | 158,670 |
| *PDGFRA* + Nucleosome | WT | C | 8 | 9 min | 4 | 15,099 |
| Naked *PDGFRA* | None | N | 5 | 2 min | 4 | 14,207 |
| Methylated *PDGFRA* | WT | N | 7 | 9 min | 4 | 175,577 |
| PDGFRA + PDS5A | WT | N | 8 | 14 min | 4 | 129,044 |
| PDGFRA + Cohesin | WT | N | 20 | 58 min | 4 | 538,958 |
| PDGFRA + PDS5A & Cohesin | WT | N | 9 | 24 min | 4 | 544,497 |

**Table S2.Lifetimes and occupancy of binding states for all experimental conditions**. Lifetimes and occupancies (fraction of time spent) of binding states of experimental conditions measured from the N-terminus and C-terminus of CTCF and variants. Errors of lifetimes represent the error on a single exponential fit of all data for experimental conditions pooled. Error bars of occupancy represent the standard deviation between replicates as shown in Table S1. Occupancy p-values were obtained by performing two-tailed Mann-Whitney tests for each condition compared against *PDGFRA* with WT CTCF probed from the same orientation as the experimental condition.

| Fraction of time spent / occupancies | | | | | | | | | |
|---|---|---|---|---|---|---|---|---|---|
| | **PDGFRA + WT N-term** | **PDGFRA + WT C-term** | **Cons. + WT N-term** | **PDGFRA + ΔN N-term** | **PDGFRA + nuc. C-term** | **Meth. PDGFRA N-term** | **PDGFRA+ WT & PDS5A N-term** | **PDGFRA+ WT & Cohesin N-term** | **PDGFRA + WT & PDS5A + Cohesin N-term** |
| **I** | 0.10 ±0.02 | 0.10 ±0.14 | 0.02 ±0.01 p=3E-4 | 0.12 ±0.05 p=0.86 | 0.17 ±0.11 p=0.08 | 0.01 ±0.01 p=1E-4 | 0.03 ±0.01 p=1E-3 | 0.10 ±0.01 p=0.98 | 0.11 ±0.02 p=0.71 |
| **II** | 0.76 ±0.05 | 0.46 ±0.19 | 0.82 ±0.02 p=8E-3 | 0.72 ±0.06 p=0.59 | 0.69 ±0.14 p=0.08 | 0.52 ±0.02 p=1E-4 | 0.83 ±0.02 p=7E-3 | 0.78 ±0.03 p=0.06 | 0.72 ±0.03 p=0.06 |
| **III** | 0.13 ±0.03 | 0.26 ±0.15 | 0.16 ±0.02 p=0.05 | 0.14 ±0.03 p=0.59 | 0.08 ±0.04 p=0.03 | 0.47 ±0.03 p=8E-4 | 0.13 ±0.01 p=0.96 | 0.11 ±0.02 p=0.24 | 0.17 ±0.01 p=2E-3 |
| **IV** | 0.01 ±0.02 | 0.17 ±0.06 | 2E-4 ±4E-4 p=8E-4 | 0.03 ±0.05 p=0.67 | 0.05 ±0.05 p=0.01 | 0.01 ±0.01 p=0.96 | 0.002 ±0.003 p=0.06 | 0.004 ±0.005 p=0.09 | 0.01 ±0.02 p=0.02 |
| **Lifetimes (ms)** | | | | | | | | | |
| **I** | 1.98 ±0.03 | 5.35 ±0.99 | 1.38 ±0.03 | 1.50 ±0.03 | 5.37 ±0.22 | 0.61 ±0.03 | 1.33 ±0.03 | 1.83 ±0.02 | 1.24 ±0.01 |
| **II** | 5.81 ±0.06 | 19.82 ±1.31 | 9.60 ±0.10 | 4.64 ±0.06 | 19.07 ±0.77 | 2.50 ±0.03 | 9.12 ±0.13 | 8.06 ±0.06 | 3.51 ±0.02 |
| **III** | 1.65 ±0.02 | 7.00 ±0.21 | 2.70 ±0.03 | 1.43 ±0.02 | 7.59 ±0.59 | 2.67 ±0.03 | 2.65 ±0.04 | 2.44 ±0.02 | 1.40 ±0.01 |
| **IV** | 10.63 ±1.00 | 5.00 ±0.13 | 5.07 ±8.35 | 7.61 ±1.21 | 12.65 ±1.55 | 0.72 ±0.029 | 3.59 ±0.60 | 6.37 ±0.62 | 12.49 ±1.54 |

**Data S1. (separate file)**
Supplementary Microsoft Excel spreadsheet for all materials, buffers, and sequences.
